# Global atlas of predicted functional domains in *Legionella pneumophila* Dot/Icm translocated effectors

**DOI:** 10.1101/2024.05.09.593423

**Authors:** Deepak T. Patel, Peter J. Stogios, Lukasz Jaroszewski, Malene Urbanus, Mayya Sedova, Cameron Semper, Cathy Le, Abraham Takkouche, Keita Ichii, Julie Innabi, Dhruvin H. Patel, Alexander Ensminger, Adam Godzik, Alexei Savchenko

**Affiliations:** Department of Microbiology, Immunology and Infectious Diseases, University of Calgary, Calgary, AB, T2N 4N1, Canada; BioZone, Department of Chemical Engineering and Applied Chemistry, University of Toronto, Toronto, ON, M5S 1A4, Canada; University of California Riverside School of Medicine, Biosciences Division; Department of Biochemistry, University of Toronto, Toronto, ON, M5S 1M1, Canada; Department of Molecular Genetics, University of Toronto, Toronto, ON, M5S 1M1, Canada

**Keywords:** Bacterial Effectors, Legionella pneumophila, Protein Modelling, Yeast Toxicity

## Abstract

*Legionella pneumophila* utilizes the Dot/Icm type IVB secretion system to deliver hundreds of effector proteins inside eukaryotic cells to ensure intracellular replication. Our understanding of the molecular functions of this largest pathogenic arsenal known to the bacterial world remains incomplete.

By leveraging advancements in 3D protein structure prediction, we provide a comprehensive structural analysis of 368 *L. pneumophila* effectors, representing a global atlas of predicted functional domains summarized in a database (https://pathogens3d.org/legionella-pneumophila). Our analysis identified 157 types of diverse functional domains in 287 effectors, including 159 effectors with no prior functional annotations. Furthermore, we identified 35 unique domains in 30 effector models that have no similarity with experimentally structurally characterized proteins, thus, hinting at novel functionalities.

Using this analysis, we demonstrate the activity of thirteen domains, including three unique folds, predicted in *L. pneumophila* effectors to cause growth defects in the *Saccharomyces cerevisiae* model system. This illustrates an emerging strategy of exploring synergies between predictions and targeted experimental approaches in elucidating novel effector activities involved in infection.

## Introduction

The Gram-negative bacterium, *Legionella pneumophila*, is an intracellular pathogen of freshwater protozoa (Abu Kwaik *et al*, 1998). The ubiquitous presence of this bacterium in human-made and natural freshwater reservoirs often leads to the accidental infection of humans from the inhalation of contaminated aerosolized water particles (Blatt *et al*, 1993; Muder *et al*, 1986). This can lead to a severe, life-threatening form of pneumonia, called Legionnaires’ disease, or a self-resolving, flu-like illness, known as Pontiac fever (Cordes & Fraser, 1980; Cunha *et al*, 2016).

Upon being phagocytosed by the eukaryotic host cell, *L. pneumophila* remodels the phagosome into a replication-permissive compartment - termed the *Legionella*-containing vacuole (LCV) (Horwitz & Maxfield, 1984; Roy *et al*, 1998; Tilney *et al*, 2001). The establishment of the LCV is dependent on the delivery of specific proteins, called “effectors”, into the host cell, which is mediated by the Dot/Icm (<u>d</u>efective in <u>o</u>rganelle trafficking/intra<u>c</u>ellular <u>m</u>ultiplication) type IVB secretion system (T4SS) - an essential molecular syringe-like complex that is conserved in all species of *Legionella* (Berger & Isberg, 1993; Ensminger & Isberg, 2009; Marra *et al*, 1992; Ninio & Roy, 2007; Segal & Shuman, 1997). The Dot/Icm effectors are involved in the manipulation of a wide variety of host cellular processes, including vesicle trafficking, protein translation, autophagy, vacuolar function, and the cytoskeleton to avoid lysosomal fusion and for the formation of the LCV (Horwitz & Maxfield, 1984; Shames, 2023; Swanson & Isberg, 1995; Tilney *et al*., 2001).

The recognition of Dot/Icm effectors for translocation is mediated through at least two mechanisms: either by a C-terminal glutamate-rich signal motif that is recognized by the T4SS component DotM or through the capture of effectors by the IcmSW proteins for delivery (Burstein *et al*, 2009; Meir *et al*, 2018; Meir *et al*, 2020). Over 360 Dot/Icm-translocated effectors have been identified in *L. pneumophila* through a variety of methods, including large-scale experimental screens (Huang *et al*, 2011; Zhu *et al*, 2011) and machine-learning approaches (Burstein *et al*., 2009). This represents the largest arsenal of bacterial effectors described to date, with effectors representing over 10% of the *L. pneumophila* proteome. Across the entire *Legionella* genus, the number of effectors is staggering, with over 18,000 unique effector-coding sequences identified to date (Gomez-Valero *et al*, 2019). This extensive arsenal of host-manipulating factors in *Legionella* species is attributed to the rapid evolution necessary for the successful survival and colonization of diverse protozoan species in the natural habitat of *Legionella* (Amaro *et al*, 2015; Gomez-Valero & Buchrieser, 2019; O’Connor *et al*, 2011; O’Connor *et al*, 2012; Park *et al*, 2020). The ability of *L. pneumophila* and other *Legionella* species to infect human macrophages, on the other hand, suggests that the effector arsenal targets conserved eukaryotic cellular processes. This hypothesis underscores how understanding the functions of individual *L. pneumophila* effectors could illuminate basic eukaryotic cellular processes and the evolutionary basis of crucial pathways required for intracellular bacterial pathogenesis. The size of this arsenal, however, presents its own experimental challenges: significant functional redundancy has been observed between effectors (O’Connor *et al*., 2011; O’Connor *et al*., 2012; Park *et al*., 2020), limiting the effectiveness of traditional forward genetic approaches to defining effector function. As a consequence, a significant number of *L. pneumophila* effectors remain functionally uncharacterized (Lockwood *et al*, 2022; Mondino *et al*, 2020b; O’Connor *et al*., 2012).

Insights into the molecular functions of *L. pneumophila* effectors have resulted primarily from analyses of their primary sequences and the detection of motifs and domains associated with known activities (Burstein *et al*, 2016; Gomez-Valero *et al*., 2019; Nachmias *et al*, 2019). A global evaluation of primary sequences revealed the prevalence of eukaryotic-like motifs/domains in *L. pneumophila* effectors, defined as predominantly (more than 75%) found in eukaryotic species (Gomez-Valero *et al*., 2019). This observation led to the hypothesis that *Legionella* acquired genes through horizontal gene transfer during co-evolution with its eukaryotic hosts (“inter-domain horizontal gene transfer”), co-opting eukaryotic genes as effectors for host manipulation (Cazalet *et al*, 2004; de Felipe *et al*, 2005; Lurie-Weinberger *et al*, 2010). Consequently, the presence of eukaryotic-like domains or sequence similarity (over 20%) to eukaryotic proteins in *Legionella* has been used to predict effector function (Gomez-Valero *et al*., 2019). This predictive method proved particularly successful for effectors sharing significant similarity with functional domains associated with eukaryotic-specific processes, such as protein ubiquitination and phosphorylation (Bruckert & Abu Kwaik, 2016; Ensminger & Isberg, 2010; Kubori *et al*, 2008; Kubori *et al*, 2010; Lee *et al*, 2020; Lee & Machner, 2018; Michard *et al*, 2015; Moss *et al*, 2019; Qiu & Luo, 2017; Quaile *et al*, 2015).

However, functional annotation of *Legionella* effectors based on their primary sequence analysis has limitations since a large proportion of these proteins do not share significant amino acid similarity with functionally characterized proteins (Gomez-Valero & Buchrieser, 2019; Mondino *et al*, 2020a). Consequently, experimental determination of effector 3D structures - primarily using X-ray crystallography as well as nuclear magnetic resonance spectroscopy, and, as of recent, cryogenic electron microscopy - proved to be instrumental in revealing otherwise cryptic domains and other molecular features indicative of their activity. For example, the structure of Lpg2511/SidC revealed the presence of an N-terminal domain with a unique fold, featuring a conserved amino acid arrangement similar to the Cys-His-Asp catalytic triad found in cysteine proteases (Hsu *et al*, 2014). Subsequent biochemical analysis showed that the catalytic triad in Lpg2511/SidC is involved in ubiquitin ligase activity via a non-canonical catalytic mechanism distinct from eukaryotic E3 enzymes (Hsu *et al*., 2014). Likewise, the crystal structure of Lpg2147/MavC revealed a domain similar in structure to ubiquitin-like protein-specific deaminase domains found in effectors of other bacterial pathogens (Samba-Louaka *et al*, 2009; Valleau *et al*, 2018). This discovery facilitated the demonstration that Lpg2147/MavC can deaminate human ubiquitin at Gln40 (Gan *et al*, 2020; Valleau *et al*., 2018) and catalyze the transglutamination of Ub via its Gln40 to Lys92 of the human UBE2N ubiquitin-conjugating enzyme (Gan *et al*., 2020; Mu *et al*, 2020). These examples underscore the advantages of molecular structural information for characterizing the activity of an effector in the host cell. Yet, the experimental determination of effector structures face numerous technical challenges (Benjin & Ling, 2020; McPherson & Gavira, 2014; Montelione *et al*, 2000); thus, the structural characterization of the *L. pneumophila* effector arsenal remains largely incomplete where more than 40% lack experimental structures and have no functional annotations associated with them in the UniProt database (UniProt, 2023).

Recently, new neural network (NN) and Large Language Model (LLM)-based computational approaches, such as AlphaFold2 (Jumper *et al*, 2021), RoseTTAFold (Baek *et al*, 2021), and Evolutionary scale modeling (ESM) Fold (Lin *et al*, 2023), have dramatically improved the accuracy of protein 3D structure prediction from primary sequences. These methods approach the accuracy of experimentally derived molecular structures for single proteins (Bertoline *et al*, 2023; Janes & Beltrao, 2024; Perrakis & Sixma, 2021), though they still face challenges in predicting multiple conformations, metal, co-factor, or ligand binding, as well as protein-protein interactions. Nevertheless, these methods allow for the analysis of structural models of large, previously uncharacterized protein families (Pinheiro *et al*, 2021) and entire proteomes (Tunyasuvunakool *et al*, 2021). Given that a large portion of the *L. pneumophila* effector arsenal remains both structurally and functionally uncharacterized, we conducted a global analysis of structural models generated by AI-based algorithms, particularly AlphaFold2, of all reported *L. pneumophila* effectors. We analyzed them for the presence of globular domains and functionally relevant structural motifs, which we then validated by leveraging the growth defect phenotype caused from the ectopic expression of effectors in the *Saccharomyces cerevisiae* model system. This approach allowed for the dramatic expansion of our functional predictions for the *L. pneumophila* effectorome, including newly predicted cysteine proteases, metalloproteases, kinases, ɑ/β hydrolases, ADP-ribosyltransferases, and glycosyltransferases. We also identified 30 effectors containing new, unique folds containing no structural similarity to any experimentally characterized 3D structure.

## Results

### *L. pneumophila* effectors carry an extensive repertoire of cryptic domains

To cast a wide net for this study, we compiled a list of *L. pneumophila* proteins from previously reported genome-wide evaluations of their translocation in a Dot/Icm-dependent manner into the host cell (Burstein *et al*., 2009; Huang *et al*., 2011; Zhu *et al*., 2011). This approach identified 368 *L. pneumophila* effector proteins (Supplementary Table 1) with 227 of them described by the UniProt database (UniProt, 2023) as “uncharacterized” or as “domain of unknown function (DUF)-containing” proteins and another 43 with a name, but no functional annotations.

A search using BlastP (Altschul *et al*, 1997) against the Protein Data Bank (PDB) repository showed that at the time of this analysis, the three-dimensional (3D) molecular structures of 41 *L. pneumophila* effectors have been experimentally characterized (Supplementary Table 1) to their complete or almost complete length (i.e. structure covered more than 90% of primary sequence with more than 90% identical residues). For an additional 44 effectors, molecular structures were available for a portion of the protein (i.e. structures covering less than 90% of primary sequence with more than 90% sequence identity) (Supplementary Table 1). Out of the remaining 283 effectors, 61 shared strong sequence similarity with structurally and/or functionally characterized proteins (Supplementary Table 1), leaving 221 effectors with no meaningful structural annotations. Consequently, more than half of the overall *L. pneumophila* effector repertoire showed no strong primary sequence similarity with any structurally and/or functionally characterized proteins. This scarcity of structural information about *L. pneumophila* effector proteins is even more apparent when related to the total length of these proteins. Only about one-third (33%) of the total effectors’ primary sequence length share strong similarity with structurally characterized proteins. We hypothesized that the gap in structural annotations can be now filled by high-quality structural predictions that can be used as a starting point for their functional characterization. In line with this hypothesis, our analysis of *L. pneumophila* effectors models generated by the AlphaFold2 algorithm (Jumper *et al*., 2021) showed confidence (pLDDT) above 70 for 78% of the primary sequence of these proteins. This level of confidence was shown to correspond to 100% correct fold assignments in numerous independent experiments, including those performed by our groups.

Hence, we analyzed Alphafold2 structural prediction models of all *L. pneumophila* effector proteins currently lacking structural coverage along with their primary sequences to identify potential functionally distinct domains (see the Methods section). Each of the predicted domains was classified based on their structural or sequence similarity to known domains in the ECOD and InterPro databases, respectively (Blum *et al*, 2021; Cheng *et al*, 2014) and, where applicable, based on previously reported experimental data. Predicted representatives of established functional domain families identified in effector protein models were further analyzed for the presence of specific functionally relevant molecular motives.

Including the AlphaFold2 models dramatically increased the number of predicted domain types identified in *L. pneumophila* effectors, as compared to previous analyses (Burstein *et al*., 2016; Gomez-Valero *et al*., 2019; Gomez-Valero *et al*, 2011), as well as public databases, such as UniProt (UniProt, 2023)(Fig. 1). We identified at least one distinct ECOD-classified domain type or structural motif in the models of 286 *L. pneumophila* effectors, with a maximum of six domains and motifs identified in the case of two effectors (Table 1). In the case of 82 effectors, we could not assign any specific ECOD-classified domain type to the predicted 3D structure. The models of these effectors were either unstructured (disordered), consisted of structural elements, such as ɑ-helical bundles and/or transmembrane helices that did not form an arrangement with significant structural similarity to a specific ECOD domain, or showed a unique, potentially novel, fold. Detailed analysis of several effector models composed of exclusively α-helical structures yielded some insights into their function, however, most of this group was excluded from the analysis based on ECOD domain assignments.

**Figure 1.**
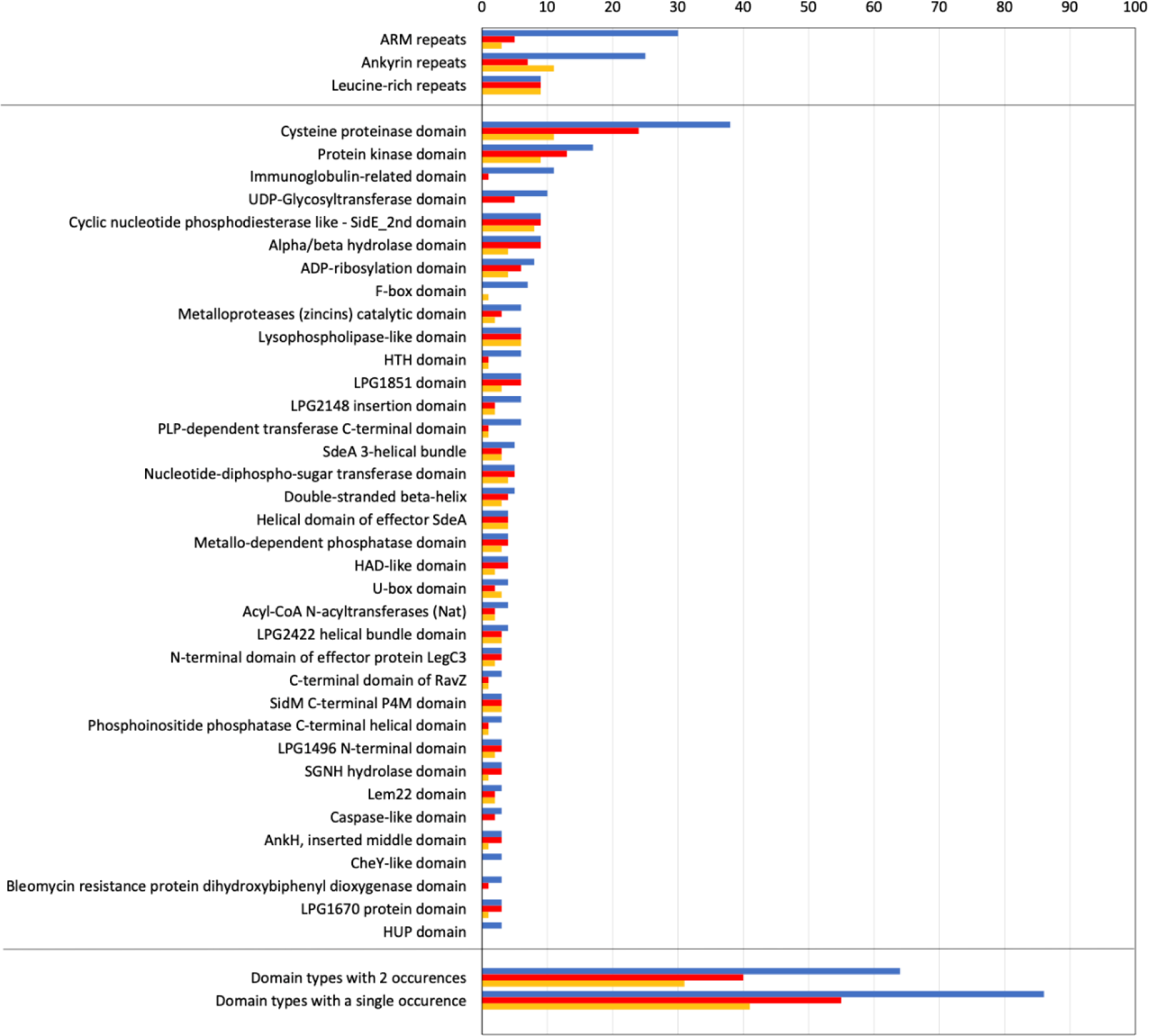
Functional Domain Occurrences from 368 *L. pneumophila* Effector Models. The number of domain instances from different ECOD domain types identified from 3D models (blue), the number of domain instances identified from remote homology (red), and by close homology (yellow). Domain instances identified based on homology are a subset of those identified in 3D models.

**Table 1.** The distribution of domain types identified in the 3D models of *L. pneumophila* effectors. Four types of protein regions that do not correspond to named ECOD domain types (unclassified helical regions, transmembrane helical regions, structurally disordered regions, and potentially novel domains) are also included.

While some models lacking defined secondary structure represented cases of the AlphaFold2 algorithm’s failure to generate a high-confidence model, most appear to be confidently predicted to be intrinsically disordered, which included both full-length disordered proteins as well as disordered regions interspaced with structured domains.

The prevalence of disordered regions in *L. pneumophila* effector models was notably higher than in the rest of *L. pneumophila* proteome. According to UniProt, 24% (90 out of 368) of effectors contain at least one disordered region, while the prevalence of such regions across non-effector *L. pneumophila* proteome is only 4.8 %. Notably, this value is more typical of eukaryotic than bacterial protein sets (Basile *et al*, 2019) and may constitute another “eukaryotic-like” feature of *L. pneumophila* effector arsenal (Hilbi & Buchrieser, 2022).

The overall repertoire of distinct domains identified in the analyzed effector models included 157 structurally diverse domain types associated with known enzymatic activities, protein-protein and protein-nucleic acid interaction functions, as well as eukaryotic-specific post-translational modification cascades (Table 1). As it could be anticipated, effector regions that corresponded to identified structural domains matched the regions modeled with high confidence. In contrast, only 5% of residues outside of identified structural motifs and domains were modeled with confidence above 50. Among categories of domains present in multiple effectors, the cysteine protease domain represents the most abundant globular domain type – identified in 37 effector models (Fig. 1B). The next largest group are protein kinase domains – found in 17 effector models. Overall, a total of 66 domain types were identified in more than one effector model, while 91 of the predicted domain types were present only in a single *L. pneumophila* effector model (Table 1). We interpret this observation as an indication of high functional diversity across the *L. pneumophila* effector repertoire. Finally, our analysis identified 35 predicted domains that showed no strong structural similarity to experimentally characterized protein structures, thus, suggesting potential novel structural folds (Table 1).

In line with previous global analyses of *L. pneumophila* effector primary sequences (Burstein *et al*., 2016; Gomez-Valero *et al*., 2019), the analysis of effector models confirmed the presence of significant number of so-called tandem repeat motifs, including armadillo (ARM) identified in 27 models, ankyrin (ANK) identified in 24 models, or leucine-rich repeats (LRR) identified in 9 models (Table 1, Fig. 1), all of which are typically associated with protein-protein interactions. Interestingly, ANK and ARM repeats were usually found in effector models that also contained other domains. In contrast, LRRs all represented the only functional element identified in effector models (Table 1). While the presence of most of these structural motifs was also predicted in previous reports using primary sequence-based tools (Burstein *et al*., 2016; Cazalet *et al*., 2004), including the 3D models as a starting point significantly expanded the number of effectors predicted to contain these structural elements (Table 1, Fig. 1).

Taken together, our analysis using the 3D models of *L. pneumophila* effectors expanded the number of predicted domains previously found in these proteins but also identified a significant number of previously unrecognized or cryptic domains, including the ones that appear to share no structural similarity with experimentally defined structures. In particular, in 199 of the 270 effector proteins annotated as uncharacterized (uncharacterized, DUF-only domains, or disordered-only regions), we have identified a domain that allowed us to provide at least a partial functional annotation. Furthermore, in the case of 15 effector models, we identified unique folds in uncharacterized domains, which were the only domain in a given protein, hence we could not assign functions based on structural similarity to a known protein.

To facilitate the follow-up functional characterization of identified functional domains, we have captured our analysis on a publicly-accessible web page https://pathogens3d.org/legionella-pneumophila. Furthermore, we initiated model-guided analysis of the potential activity of predicted domains using the budding yeast *Saccharomyces cerevisiae* model cell system. Heterologous expression of individual *L. pneumophila* effectors in this eukaryotic cell often causes severe growth defects that can be exploited for elucidation of effector biochemical activity (Belyi *et al*, 2012; Campodonico *et al*, 2005; de Felipe *et al*, 2008; Fu *et al*, 2022; Gaspar & Machner, 2014; Guo *et al*, 2014; Heidtman *et al*, 2009; Nevo *et al*, 2014; Shohdy *et al*, 2005; Urbanus *et al*, 2016; Viner *et al*, 2012; Xu *et al*, 2010). Our analysis identified 42 distinct domain types in 64 *L. pneumophila* effectors causing toxicity in yeast model system. Previous studies have linked this phenotype to the function of a distinct domain in 27 of these effectors. Among the remaining proteins, we prioritized effectors with cryptic functional domains that could not be readily recognized by primary sequence analysis tools. Below we discuss in detail the main types of cryptic domains identified in 3D models of *L. pneumophila* effectors triggering toxicity in yeast before probing predicted active sites of 14 such domains in this model system by mutagenesis.

### Predicted cysteine protease domains in *L. pneumophila* effectors show functional diversity

Representatives of the cysteine protease domain have been characterized across all kingdoms of life and have been the subject of extensive analysis due to their critical roles in diverse biological processes, including protein degradation, cell signaling, and the immune response (Lopez-Otin & Bond, 2008; Verma *et al*, 2016). As per their family name, these enzymatic domains contain a conserved cysteine residue typically paired with a histidine and an aspartate, or asparagine, residues arranged into a catalytic triad (Rawat *et al*, 2021). Some cysteine proteases contain a catalytic dyad formed by cysteine and histidine residues (Rawat *et al*., 2021). The active site containing these catalytic residues are usually located in a cleft between the two lobes of the ɑ/β domain (Hofer *et al*, 2020; Verma *et al*., 2016).

The presence and function of cysteine protease domains have been previously reported for 14 *L. pneumophila* effectors, and the 3D structures of several of these cysteine protease domains have been experimentally defined (Table 2). Expanding on these previous studies, our analysis of effector structural models suggested the presence of cysteine protease domains in 21 additional effectors (Table 2). Notably, cysteine protease domains were identified in the Lpg0284-Lpg0285 (∼23% sequence similarity) and Lpg1965-Lpg1966 (26% sequence similarity) effector pairs. Primary sequence similarity and co-localization of corresponding genes in each pair are suggestive of a tandem gene duplication event (Fig. 2A-B). In a similar case, the genetically co-localized effectors, Lpg2147/MavC and Lpg2148/MvcA were shown to be functionally diverse despite their significant primary sequence and structural similarity (Valleau *et al*., 2018).

**Figure 2.**
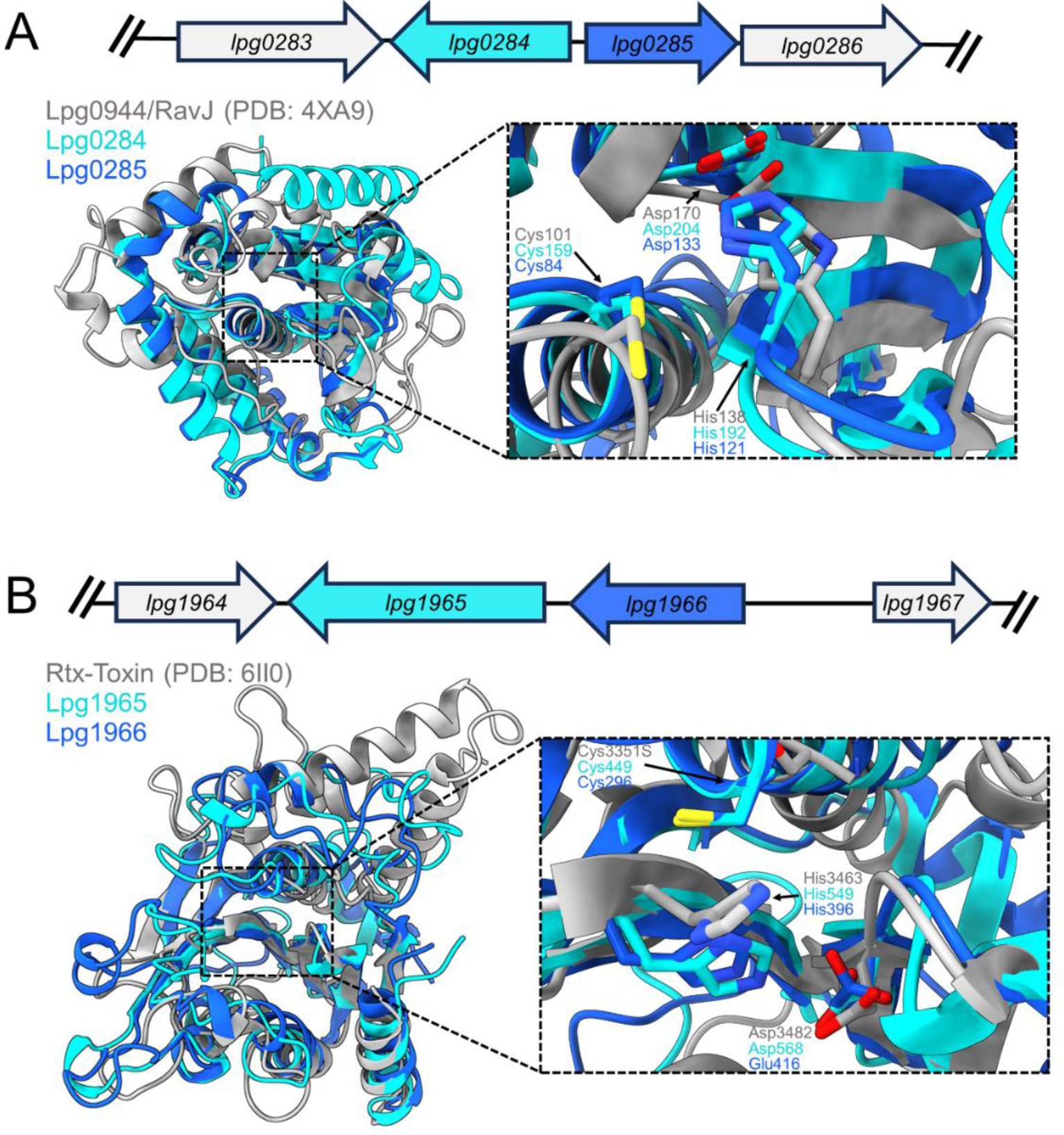
Gene co-localization reveals two pairs of paralogous effectors with cysteine protease domains. A. Genomic organization of *lpg0284* and *lpg0285* in the *L. pneumophila* strain Philadelphia (Top). Superimposition of Lpg0284 (residues 66-290) and Lpg0285 (residues 1-181) models onto the crystal structure of Lpg0944/RavJ (residues 1-228) (Urbanus *et al*., 2016), followed by a zoom-in of the putative active sites and their catalytic residues (shown in sticks) of these paralogs (Bottom). B. The gene diagram of *lpg1965* and *lpg1966* from the genome of the *L. pneumophila* strain Philadelphia (Top). The structural alignment of Lpg1965 (residues 390-611) and Lpg1966 (residues 251-470) with the Rtx-Toxin from *Vibrio vulnificus* (residues 3309-3562) (Lee *et al*, 2019). Next to the structural alignment is a zoomed-in image of the potential active site pocket and residues that form the catalytic triad (shown in sticks) identified in these paralogous effectors (Bottom).

**Table 2.** Effectors previously characterized to have a cysteine protease domain and the expansion of this domain and the putative catalytic residues identified from the 3-D model analysis of *L. pneumophila* effectors against the top structurally and functionally characterized hit from the FATCAT server.

| Effector | Potential, or Previously Described, Catalytic Residues | Reference |
| --- | --- | --- |
| Lpg0056 | Cys19-His182-Asp200 | This Study |
| Lpg0126 | Cys21-His228-Asn226 | This Study |
| Lpg0160/RavD | Cys12-His94-Ser111 | (Wan *et al.*, 2019b) |
| Lpg0196 | Cys320-His32-Ser89 | This Study |
| Lpg0227/Ceg7/LotD | Cys13-His256-Asp6 | (Kang *et al.*, 2023) |
| Lpg0234/SidE | Cys117-His64-Asp80 | (Sheedlo *et al.*, 2015) |
| Lpg0284 | Cys159-His192-Asp204 | This Study |
| Lpg0285 | Cys84-His121-Asp133 | This Study |
| Lpg0403 | Cys73-His217-Asp55-Asn232 | This Study |
| Lpg0944/RavJ | Cys101-His138-Asp170 | (Urbanus *et al.*, 2016) |
| Lpg1110 | Cys109-His137-Asp148 | This Study |
| Lpg1120 | Cys23-His205-Asn228 | This Study |
| Lpg1148/LupA | Cys-252-His183-Asp207 | (Urbanus *et al.*, 2016) |
| Lpg1355/SidG | Cys623-His57-Asp158-Glu162 | This Study |
| Lpg1387 | Cys403-His38-Ser104 | This Study |
| Lpg1621/Ceg23/LotB | Cys29-His270-Asp29 | (Ma *et al.*, 2020) |
| Lpg1683/RavZ | Cys258-His176-Asp197 | (Horenkamp *et al.*, 2015) |
| Lpg1797 | Cys314-His45-Ser102 | This Study |
| Lpg1909 | Cys166-His22 | This Study |
| Lpg1949 | Cys174-His105-Asp124 | This Study |
| Lpg1965 | Cys449-His549-Asp568 | This Study |
| Lpg1966 | Cys296-His396-Glu416 | This Study |
| Lpg2143 | Cys138-His259-Asp274 | This Study |
| Lpg2147/MavC | Cys74-His231-Gln252 | (Valleau *et al.*, 2018) |
| Lpg2148/MvcA | Cys83-His244-Gln265 | (Valleau *et al.*, 2018) |
| Lpg2153/SdeC | Cys118-His64-Asp80 | (Sheedlo *et al.*, 2015) |
| Lpg2156/SdeB | Cys123-His69-Asp85 | (Sheedlo *et al.*, 2015) |
| Lpg2157/SdeA | Cys118-His64-Asp80-Asn114 | (Sheedlo *et al.*, 2015) |
| Lpg2215 | Cys33-His145-Asn162 | This Study |
| Lpg2248/LotA/Lem21 | Domain 1 (residues 1-294): Cys13-His237-Asn239 and Domain 2 (residues 296-544): Cys303-Asn296-His535 | (Takekawa *et al.*, 2022; Warren *et al.*, 2023) |
| Lpg2433 | Cys26-His205-Asp229-Asn226 | This Study |
| Lpg2529/LotC/Lem27 | Cys24-His304-Asp17 | (Shin *et al.*, 2020) |
| Lpg2538 | Cys299-His231 | This Study |
| Lpg2586 | Cys106-His291-Asn320-Gln100 | This Study |
| Lpg2907 | Cys263-His192-Asp215 | This Study |
| Lpg2952 | No functional residues could be assigned | This Study and (Hermanns *et al.*, 2020) |

In 11 out of 21 effectors with predicted cysteine protease domain, we were able to identify an active site cavity with cysteine, aspartate, and histidine residues arranged in a potential catalytic triad (Table 2). In five additional predicted cysteine protease domains, the putative catalytic site featured an asparagine instead of a catalytic aspartate residue (Table 2).

For the Lpg1949, Lpg2538, and Lpg2907 effector models, the predicted cysteine protease domains showed similarity to members of the YopJ effector family. The members of this family found in such human pathogens, such as *Yersinia* species, and in several plant pathogens have been only associated with type 3 secretion system (T3SS) (Lewis *et al*, 2011; Ma & Ma, 2016; Meinzer *et al*, 2012; Mukherjee *et al*, 2006; Orth *et al*, 2000; Xia *et al*, 2021). Cysteine protease domains in YopJ effectors demonstrate acetyltransferase activity, which is activated by the eukaryote-specific co-factor inositol hexakisphosphate (IP_6_) (Mittal *et al*, 2010). This co-factor binds to a conserved positively charged pocket on the effector surface (Mittal *et al*., 2010). Along with the identification of active site pockets made of either a Cys-His-Asp/Glu triad or Cys-His dyad (Mukherjee *et al*., 2006; Orth *et al*., 2000; Tomar *et al*, 2023; Zhang *et al*, 2017), the analysis of the corresponding predicted domains in Lpg1949, Lpg2538, and Lpg2907 effectors (Fig. 3A) suggested the presence of a positively charged pocket typical of IP_6_ binding (Appendix Fig. S1). A previous report determined that Lpg1949 functions as an acetyltransferase rather than a protease (Hermanns *et al*, 2020). However, the host substrate and biological significance of this effector during infection remains to be determined. Accordingly, based on our analysis, we hypothesize that along with Lpg1949, Lpg2538, and Lpg2907 effectors may also demonstrate YopJ-like acetyltransferase activity; therefore, potentially expanding the YopJ enzyme family into effectors translocated by the *L. pneumophila* T4SS.

**Figure 3.**
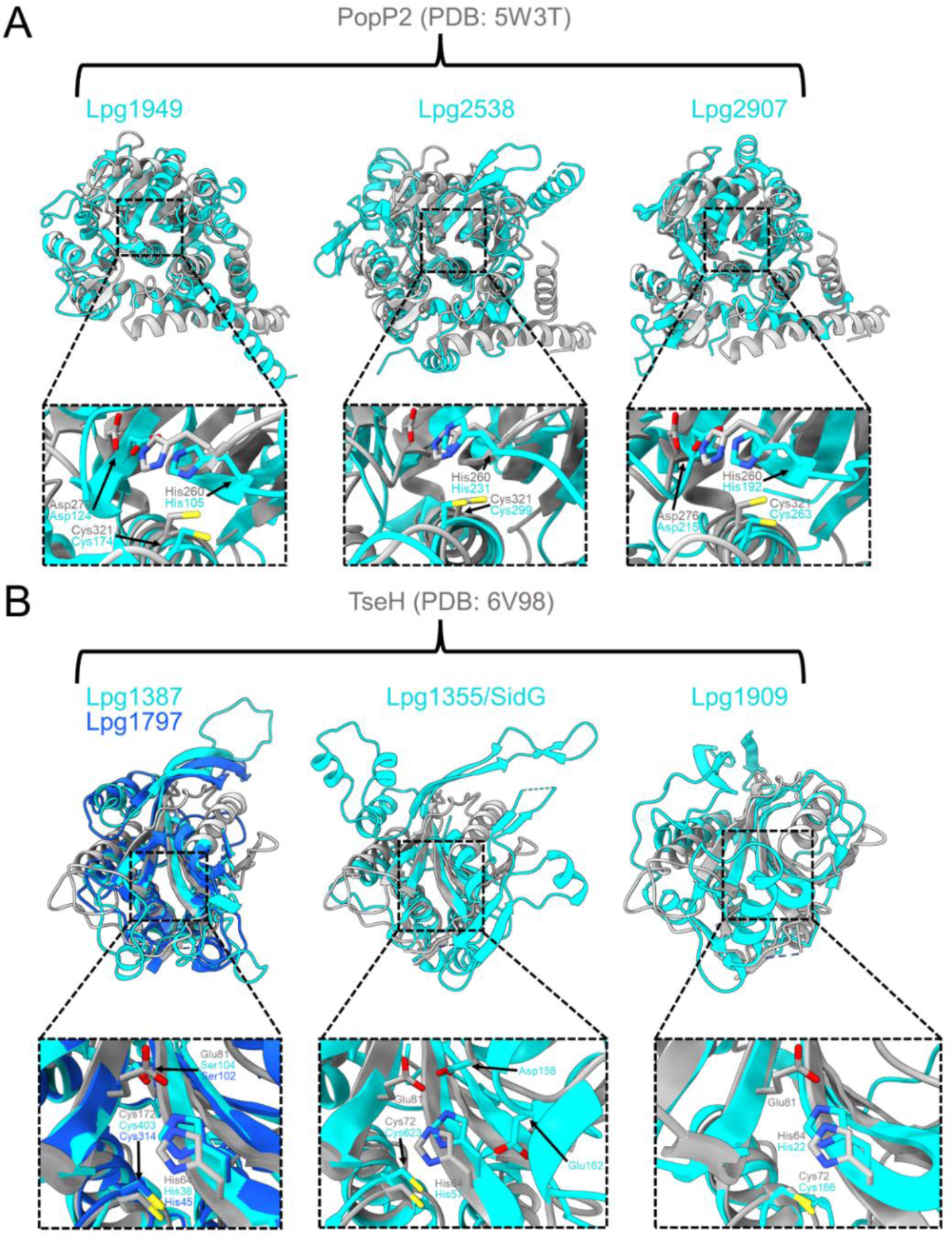
*L. pneumophila* effectors that display structural similarities to members of the YopJ-like T3SS effector family and the T6SS effector, TseH, from *Vibrio cholerae*. A. Superimpositions of the Lpg1949 (residues 6-327), Lpg2538 (residues 25-463), and Lpg2907 (residues 61-370) models onto the YopJ-like effector, PopP2 (PDB: 5W3T, residues 81-420), from *Ralstonia solanacearum* (Zhang et al., 2017) (Top). The zoom-in panels show the putative active site and residues that form the catalytic dyad or triad. B. Structural alignments of the Lpg1355/SidG (residues 1-195 and 563-665), Lpg1387 (residues 1-139 and 359-462), Lpg1797 (residues 8-126 and 226-384), and Lpg1909 (residues 1-215) models onto the TseH crystal structure (PDB: 6V98, residues 22-223) (Hersch *et al*., 2020). The pop-out is a view of the potential active site harboring residues that could be involved in catalysis (sticks).

Previous studies have identified Lpg0227, Lpg2248, Lpg1621, and Lpg2529 effectors as novel bacterial deubiquitinases, which, due to their primary sequence and structural similarity to eukaryotic ovarian-tumor deubiquitinases (Kang *et al*, 2023; Kubori *et al*, 2018; Schubert *et al*, 2020), have been termed the *<u>L</u>egionella* <u>OT</u>U-like DUBs (Lot-DUBS). Typically, Lot-DUBs harbor a catalytic triad consisting of Cys-His-Asp residues. Based on structural similarity, our analysis suggested that a similar domain may also be present in Lpg2952. However, the Lpg2952 model did not reveal any suitable candidates for the catalytic residues typical of Lot-DUB enzymes (Appendix Fig. S2). This observation is in line with a previous report that was not able to demonstrate DUB activity for Lpg2952 (Hermanns *et al*., 2020), raising the possibility that the activity of Lpg2952 may have deviated from Lot-DUBs.

Cysteine protease domains in the structural models of Lpg1355/SidG, Lpg1387, Lpg1797, and Lpg1909 share structural features with the NlpC/P60 cysteine endopeptidase family, which include known peptidoglycan (PG) degraders (Griffin *et al*, 2023; Hersch *et al*, 2020). Notably, the structurally characterized effector TseH from *Vibrio cholerae* translocated by type 6 secretion system (T6SS) (Altindis *et al*, 2015; Hersch *et al*., 2020) also belongs to this protein family (Fig. 3B). While the specific enzymatic activity of TseH remains enigmatic, the TseH structure features two lobes that form a pocket housing glutamate, histidine, and cysteine residues shown to be essential for its activity (Hersch *et al*., 2020). Similarly, we identified a histidine and a cysteine at similar positions in the interlobar pocket of Lpg1355/SidG, Lpg1387, Lpg1797, and Lpg1909 models (Fig. 3B). Furthermore, the Lpg1387 and Lpg1797 models also include a serine residue (Ser104 and Ser102, respectively) corresponding to the catalytically important glutamate in TseH (Fig. 3B). In the case of Lpg1355/SidG, we identified an adjacent ɑ-helix in the predicted catalytic pocket that harbors two residues, Asp158 and Glu162, orientated in a position that suggests an involvement in the biochemical activity of this effector (Fig. 3B). Within the Lpg1909 model, we were unable to assign an appropriate third residue to match the catalytic triad found in TseH or other structurally characterized cysteine proteases, suggesting that this effector potentially relies on a Cys166-His22 dyad for catalysis (Fig. 3B).

Finally, Lpg2586, also identified as a T4SS effector containing a potential cysteine protease domain, shares high sequence similarity (∼30%) with the *L. pneumophila* effector Lpg2622. The latter effector has been associated with the type II secretion system (T2SS) of this bacterium and was characterized as a member of the C1 peptidase family (Gong *et al*, 2018). Along with the experimentally defined structure of Lpg2622 (PDB 6A0N), the model of Lpg2586 also contains a unique hairpin-turn-helix motif, which was shown to be essential for Lpg2622 protease activity (Appendix Fig. 3A-B) (Gong *et al*., 2018). Lpg2586 also possesses an N-terminal β-sheet that was shown to be involved in regulating the activity of Lpg2622 (Gong *et al*., 2018). Our analysis also suggested that residues Cys106, His291, Asn320, and Gln100 in Lpg2586 may be important for catalysis based on the role of equivalent residues demonstrated for Lpg2622 (Appendix Fig. S3A, Table 2).

### Identification of three additional metalloprotease domain-containing effectors in the *L. pneumophila* arsenal

Metalloproteases are important components of multiple biological processes in eukaryotes. These enzymes facilitate the degradation of extracellular membrane proteins, glycoproteins, growth factors, cytoskeletal proteins, and cytokines – which, in turn, regulate apoptotic, cellular differentiation, and proliferation pathways (de Almeida *et al*, 2022; Parks *et al*, 2004; Sternlicht & Werb, 2001). When compared to cysteine proteases, metalloproteases have several distinct structural features essential for their functionality. Typically, these enzymes contain an ɑ-helical pro-domain that regulates the protease activity and a catalytic domain consisting of roughly five β-strands and three ɑ-helices (Laronha & Caldeira, 2020). The catalytic pocket of metalloproteases is made of an ɑ-helix harboring a highly conserved sequence motif: His-Glu-X-X-His-X, where “X” represents any amino acid (Laronha & Caldeira, 2020; Sternlicht & Werb, 2001). Of importance are the histidine residues in this motif that are required for the coordination of a divalent cation, typically a Zn^2+^, that is involved in peptide bond hydrolysis (Ra & Parks, 2007).

Only two *L. pneumophila* effectors (Lpg0969/RavK and Lpg2999/LegP) have been identified as metalloproteases based on primary sequence analysis (de Felipe *et al*., 2005; Liu *et al*, 2017). Lpg0969/RavK was shown to cleave actin to prevent the formation of actin polymers when ectopically expressed in HEK293T cells and during *L. pneumophila* infection. However, the biological significance of this cleavage remains unclear (Liu *et al*., 2017). In contrast, the function of Lpg2999/LegP as a canonical metalloprotease has not been experimentally validated and the host substrate also remains unknown (de Felipe *et al*., 2005). Therefore, we confirmed the presence of the catalytic motif in Lpg2999/LegP, which has not been previously described (Fig. 4A). Furthermore, our analysis suggested three additional effectors (Lpg0041, Lpg1667, and Lpg2461) harboring the metalloprotease catalytic domain (Fig. 4A-B). Each of the models of these three effectors contains the His-Glu-X-X-His-X motif typical of the metalloprotease fold (Fig. 4A-B, Table 3). Notably, among the five metalloprotease domain-containing effector models, only Lpg0041 and Lpg1667 contained an additional structural element. The Lpg0041 model contains an additional C-terminal structural element composed of the three tandem beta-sandwiches, whereas Lpg1667 is predicted to contain a single β-sandwich spanning residues 57 to 176. We hypothesize that this structural element contributes to the recognition of the host substrate for these effectors.

**Figure 4.**
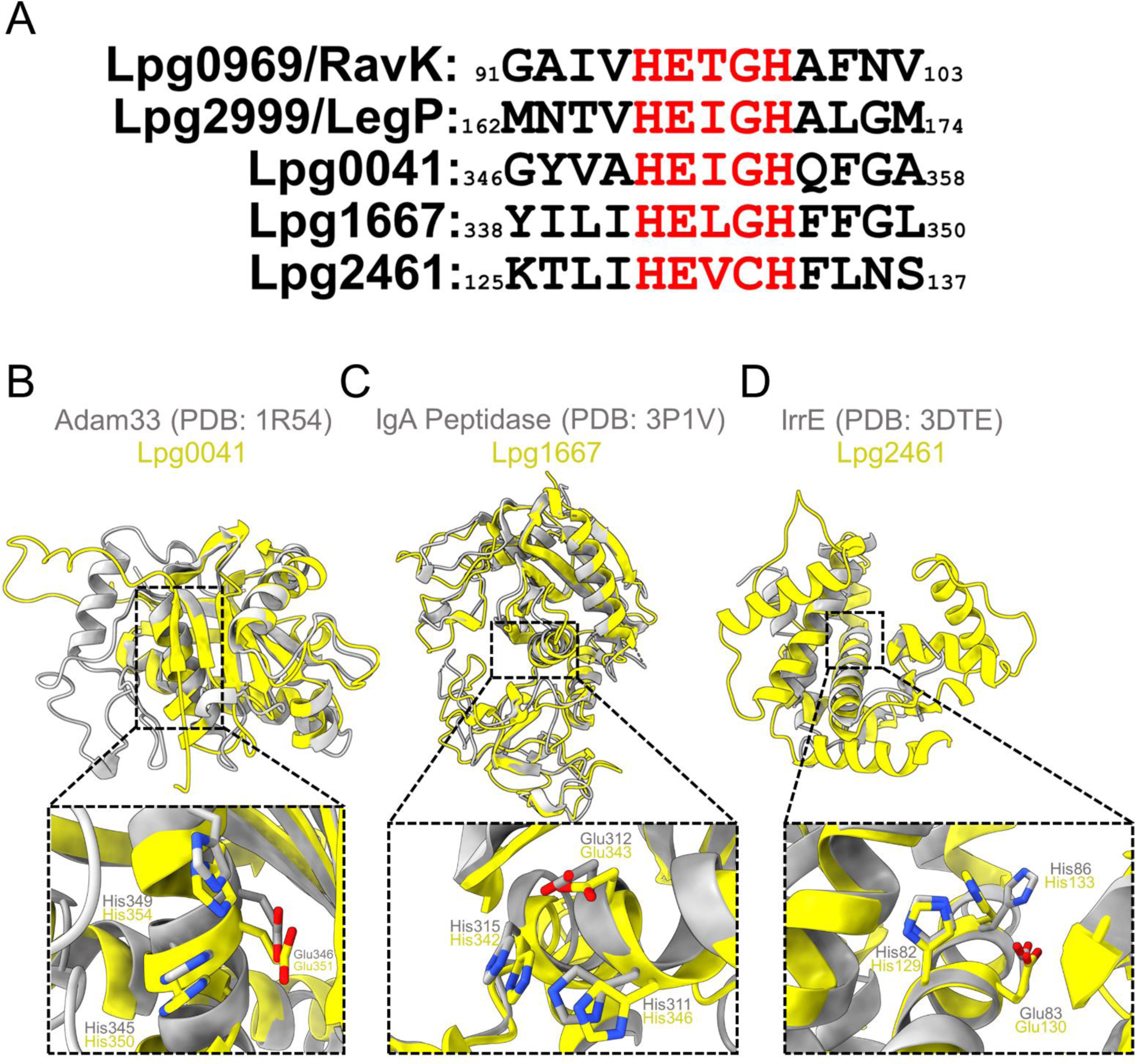
Expansion of effectors harboring a metalloprotease fold. A. Sequence alignment of the metalloprotease motif in *L. pneumophila* effectors that were previously identified and found in this study. B-D. Superimposition of Lpg0041 (residues 174-351) with human Adam33 (residues 209-409) (Orth *et al*, 2004), Lpg1667 (residues 177-455) with IgA peptidase from *Bacteroides ovatus* (residues 159-425), and Lpg2461 (residues 1-205) with IrrE from *Deinococcus derserti* (residues 1-281) (Vujicic-Zagar *et al*, 2009). Below is a close view of the potential catalytic residues (yellow sticks) of each effector model determined by their top structural hit (grey sticks) from our FATCAT analysis.

**Table 3.** Summary of the catalytic motif of previously characterized *L. pneumophila* metalloprotease domain-containing effectors and those identified from the 3D model analysis.

| Effector | Potential, or Previously Described, Catalytic Residues | Reference |
| --- | --- | --- |
| Lpg0041 | His350-Glu351-Ile352-Gly353-His354 | This Study |
| Lpg0969/RavK | His95-Glu96-Thr97-Gly98-His99 | (Liu *et al.*, 2017) |
| Lpg1667 | His342-Glu343-Leu344-Gly345-His346 | This Study |
| Lpg2461 | His129-Glu130-Val131-Cys132-His133 | This Study |
| Lpg2999/LegP | His166-Glu167-Ile168-Gly169-His170 | This Study |

### Expansion of the repertoire of *L. pneumophila* effector kinases

Kinases are important mediators of multiple biological functions, such as signal transduction, protein function modulation, and modification of small molecules, including lipids and carbohydrates (Fabbro *et al*, 2015; Oruganty *et al*, 2016; Pereira *et al*, 2011; Rauch *et al*, 2011). There are multiple molecular folds associated with this biochemical activity, but the most prevalent is the Protein Kinase fold that adopts a bi-lobed structure comprised of a smaller, all β-strand N-terminal domain and a larger C-terminal mixed ɑ/β domain, with the two domains connected via a short, linear region called the hinge (Arter *et al*, 2022). The ATP binding site is localized close to the hinge in the cavity formed between the two lobes, with its adenine ring nestled in a hydrophobic pocket and also forming hydrogen bonds between its purine nitrogens and residues in the hinge (Arter *et al*., 2022). In accordance with the catalytic mechanism, the substrate binding site is also found in this cavity, placing it in proximity to the ATP γ-phosphate (Arter *et al*., 2022). Other salient features of the Protein Kinase fold include the “catalytic loop,” which harbors aspartate residues that coordinate Mg^2+^ ions interacting with ATP phosphate oxygens; a glycine-rich loop (also called the “P-loop”) that interacts with the ATP phosphate oxygens; and an “activation segment” containing an Asp-Phe-Gly (DFG) sequence and often a tyrosine residue, both of which are important in the regulation of kinase activity as the activation segment is often disordered in the non-phosphorylated state (Arter *et al*., 2022; Leipe *et al*, 2003; Nolen *et al*, 2004; Reinhardt & Leonard, 2023). ATP grasp is another common molecular fold associated with kinases which usually act on small molecules (Fawaz *et al*, 2011). This fold is founded on two ɑ/β domains that bind ATP in the interdomain cleft (Fawaz *et al*., 2011). We identified 17 effector kinases in *L. pneumophila*, with 15 of these falling into the ECOD T group “Protein kinase” and two into the ECOD T group “ATP-grasp” (Table 4 and 5)

**Table 4.** A summarization of kinase domain-containing effectors from *L. pneumophila* identified from predicted models and their putative catalytic residues.

| Effector | Potential, or Previously Described, Catalytic Residues |  | Reference |
| --- | --- | --- | --- |
|  | **Lys/Asp - Glu pair** | **DFG motif** |  |
| Lpg0208/LegK4 | Lys110-Glu125 | Asp219-Phe220-Gly221 | (Flayhan *et al.*, 2015) |
| Lpg1316 | Lys93-Glu103 | Asp203-His204-Glu205 | This Study |
| Lpg1317 | Lys62-Glu73 | Asp169-His170-Glu171 | This Study |
| Lpg1408 | Arg101-Glu116 | Asp259-Trp260-Glu261 | This Study |
| Lpg1483/LegK1 | Lys121-Glu137 | Asp244-Phe245-Gly246 | (Ge *et al.*, 2009) |
| Lpg1924/LegK7 | Arg209-Glu219 | Asp324-Arg325-Lys326 | (Lee *et al.*, 2020) |
| Lpg2050 | Lys57-Glu87 | Asp178-Leu179-Asp180 | This Study |
| Lpg2137/LegK2 | Lys112-Glu128 | Asp223-Ala224-Gly225 | (Hervet *et al.*, 2011; Michard *et al.*, 2015) |
| Lpg2155/SidJ | Lys367-Glu381 | Asp542-Leu543-Gly544 | This Study |
| Lpg2322/AnkK/LegA5 | Lys43-Glu56 | Asp201-His202-Asp203 | This Study |
| Lpg2508 | Lys305-Glu319 | Asp480-Leu481-Gly482 | This Study |
| Lpg2556/LegK3 | Lys87-Glu110 | Asp207-Tyr208-Gly209 | This Study |
| Lpg2603/Lem28/SdmB | Lys114-(missing) | Asp225-Leu226-Asp227 | (Sreelatha *et al.*, 2020) |
| Lpg2975/MavQ | Lys46-(missing) | Asp160-Phe161-Asp162 | (Hsieh *et al.*, 2021) |

Three *L. pneumophila* effector kinases, Lpg0208/LegK4, Lpg1924/LegK7, and Lpg2975/MavQ, have been structurally characterized (Black *et al*, 2019; Flayhan *et al*, 2015; Hsieh *et al*, 2021; Lee *et al*., 2020; Sreelatha *et al*, 2020; Sulpizio *et al*, 2019). Two additional effectors, Lpg1483/LegK1 and Lpg2137/LegK2, although lacking structural characterization have been shown to exhibit kinase activity (Ge *et al*, 2009; Hervet *et al*, 2011; Michard *et al*., 2015). Our analysis suggested that 9 additional effectors contain domains falling into the T group of the “Protein kinase” domain category. With the exception of Lpg1684, Lpg1924, and Lpg1925, all these predicted domains contain the essential structural and catalytic motifs characteristic of kinase enzymatic activity. Comparative analysis of the effector kinase models with experimentally characterized kinase structures in the PDB suggested that the models of Lpg1483/LegK1 (Ge *et al*., 2009), Lpg2050 and Lpg2556/LegK3 contain the necessary features of *bona fide* protein kinases (Fig. 5A), while the models of Lpg1316/RavT, Lpg1317/RavW, Lpg2322/LegA5/AnkK (Ledvina *et al*, 2018) are more structurally similar to kinases targeting lipids (Fig. 5B). The Lpg2322/LegA5/AnkK effector has been experimentally shown to be a phosphatidylinositol 3-kinase (PI3K) (Ledvina *et al*., 2018). Another effector we identified as containing a kinase-like domain - Lpg2508/SdjA – has been shown to regulate the activity of other effectors through glutamylation activity and is thus considered a pseudokinase (Song *et al*, 2021).

**Figure 5.**
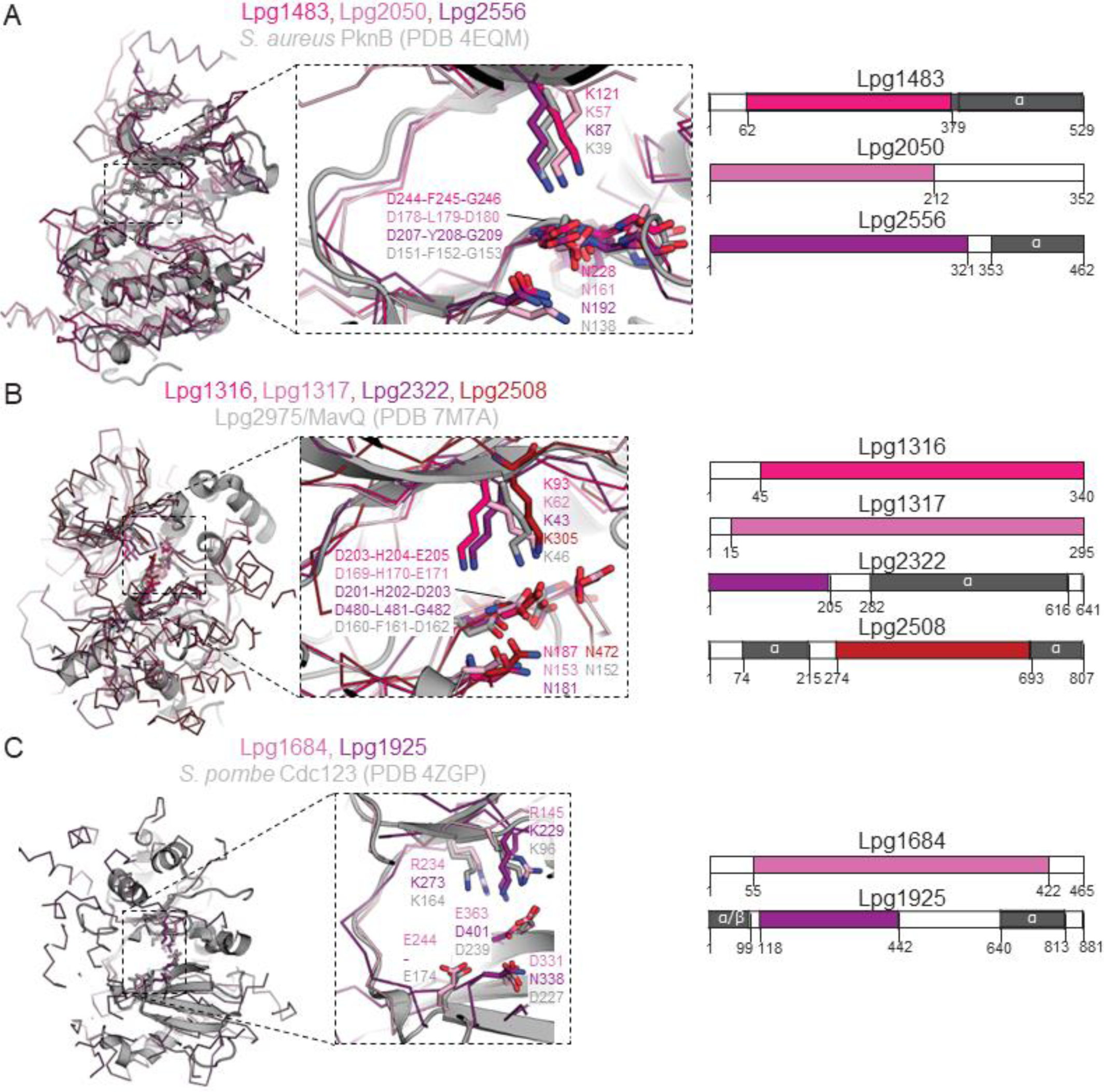
*L. pneumophila* effector kinases are the second largest functional group of effectors in the arsenal. A. Putative protein kinases, B. putative small molecule kinases, and C. putative ATP grasp fold kinases. For each panel, left = overlay of effector kinases (shades of purple/pink) with a characterized, reference structure from the PDB in grey. Right = Primary domain architecture, with kinase domain colored in shades of purple/pink and alpha-helical regions colored grey.

In our analysis of the remaining effector models with the protein kinase domain, we observed significant variation from the canonical motifs established for functional kinases, particularly in the composition of the activation loop and P-loop (Table 4). For example, the models of Lpg1316, Lpg1408, and Lpg2322 featured DHE (residues 206-208), DWE (residues 262-264), and DHD (residues 201-203) sequences, respectively, instead of the DFG motif typical of the activation loop sequence in canonical kinases. Also, in the Lpg1316 model, the P-loop contained a stretch of three serine residues (SSS, residues 77-79), while typical kinases contain at least one glycine at these positions. Lpg1316 shares 45% of sequence identity with Lpg1317 and is predicted to adopt a very similar structure. As in the case of other effector pairs which appear to originate from tandem gene duplication that were mentioned above, follow-up functional analysis will establish whether these paralogous effectors have a functionally redundant role when translocated in eukaryotic cells or have diversified in their host targets.

The kinase domain identified in the Lpg2050 model is very similar to the structure of the *Shigella* effector OspG (Grishin *et al*, 2014b; Pruneda *et al*, 2014) or the *E. coli* effector NleH (Grishin *et al*, 2014a), both of which require interaction with other proteins to trigger their protein kinase activities.

The model of Lpg1408/LicA is most similar to the structures of the choline kinase LicA from *Streptococcus pneumoniae* (Wang *et al*, 2015), as well as to aminoglycoside phosphotransferases (Wang *et al*, 2014), which act on small molecule substrates. Notably, Lpg1408/LicA was shown to interact with a component of the host cell ubiquitination machinery (i.e. the Skp1 protein) through its F-box domain, but not with a full E3 ligase complex, illustrating that the role of this effector and its kinase domain requires further analysis (Ensminger & Isberg, 2010).

The models of Lpg1684 and Lpg1925 effectors show similarity to “ATP-grasp” kinases. In this fold, the ATP binding site is composed of multiple charged residues, including Glu/Asp and Lys/Arg that interact with magnesium ions or the phosphate oxygens of ATP (Fawaz *et al*., 2011) (Table 5). Both of these effector kinase models contain such charged residues and are most similar in structure to the kinases targeting protein substrates (Fig. 5C). Notably, the predicted kinase domain in the latter effector contains an ɑ-helical insertion in its N-terminal lobe, corresponding to residues 122 to 202 and 235 to 267. The Lpg2147/MavC effector, while not a kinase, contains an insertion within its Cif domain that contributed to the recognition of a specific host protein target, illustrating the potential functional implications of insertions in the enzymatic domains of effectors (Valleau *et al*., 2018).

**Table 5.** The identification of two ATP grasp kinase domain-containing effectors from *L. pneumophila* and their catalytic residues from this study.

| Effector | Potential, or Previously Described, Catalytic Residues | Reference |
| --- | --- | --- |
| Lpg1684 | Arg145-Arg234-Glu244-Asp331-Glu363 | This Study |
| Lpg1925 | Lys229-Lys273-Asn388-Asp401 | This Study |

Structural elements beyond the “core” kinase domain often contribute to the recognition and positioning of kinase substrates or regulation of kinase activity (Pereira *et al*., 2011). Specifically, interactions between the Lpg1924/LegK7 kinase effector with host protein MOB1A involve the N-terminal α-helical domain in addition to the “core” kinase domain (Lee *et al*., 2020). Therefore, we analyzed the structural models of effectors with predicted kinase domains for the presence of such additional structural elements. Each of Lpg0208/LegK4, Leg1483/LegK1, Lpg2322, Lpg2508/SdjA, and Lpg2556 are predicted to harbor their kinase domains in the N-terminal portion of the protein, followed by ɑ-helical bundles; this region in Lpg0208/LegK4 may adopt an ARM fold, as mentioned above. Lpg2137/LegK2 model also features the kinase followed by an ɑ-helical bundle, plus a small ɑ/β domain preceding the kinase domain at the N-terminus. According to the model, this N-terminal domain is predicted to pack against the β-lobe of the kinase. The model of another effector kinase, Lpg1925, also reveals a multi-domain composition, with an ɑ/β structure at its N-terminus, the kinase domain including an ɑ-helical insert followed by a long ɑ-helical hairpin that packs against the kinase domain, another ɑ/β domain, and finally an ɑ-helical bundle.

Overall, our analysis expanded on the previous prediction of effectors with kinase domains based on the analysis of effector primary sequences (Gomez-Valero *et al*., 2019). This analysis also expands on the list of kinase effectors identified as part of the global survey of *Legionella* kinases (Krysinska *et al*, 2022) by identifying three additional effectors with potential kinase domains, specifically Lpg1317/RavW, Lpg1684, and Lpg2508/SdjA. Notably, this latter report suggested three additional kinases encoded by Lpg1930, Lpg2380, and Lpg2490/LepB to be part of the kinase effector arsenal. The structure of the Lpg2490/LepB effector has been previously characterized and its function was determined to be a GTPase activating protein (GAP) acting on host Rab1 (Mihai Gazdag *et al*, 2013). For the Lpg1930 or Lpg2380 proteins, we were not able to identify any report of experimentally confirmed Dot/Icm translocation and thus these proteins were not included in our analysis set.

### ɑ/β hydrolase domains are recurring in *L. pneumophila* effectors

The specific activities of ɑ/β hydrolases can vary widely between members of this superfamily (Kourist *et al*, 2010; Nardini & Dijkstra, 1999). The general structure of ɑ/β hydrolases comprises a central six-stranded central β-sheet surrounded by α-helices, with the ligand binding site found at the “top” of the β-sheet where the C-terminal ends of each β-strand align. Members of this family typically harbor a catalytic triad formed of a serine residue fulfilling the role of the catalytic nucleophile, a histidine and an acidic residue (almost always an aspartate), localized to loops between β-strands. An important distinguishing feature of some ɑ/β hydrolases is the presence of a “lid” subdomain inserted into the central β-sheet, which varies in size and structural features (Kourist *et al*., 2010; Nardini & Dijkstra, 1999). This subdomain often forms part of the ligand binding cleft and can create solvent-excluded pockets for binding hydrophobic compounds such as lipids. A subgroup called SGNH hydrolases share a common three-layer α/β/α structure and unites a group of enzymes with diverse specific activities, including carbohydrate esterase, thioesterase, protease, arylesterase, and lysophospholipase (Anderson *et al*, 2022).

Our analysis indicated that nine *L. pneumophila* effector models contained the ɑ/β hydrolase domain (ECOD T group “alpha/beta-hydrolases”), and three effector models contained domains reminiscent of SGNH hydrolases (ECOD T group “SGNH hydrolase”) (Table 6). Only three of these effectors - Lpg1642/SidB, Lpg1907, and Lpg2911 - have been previously reported to possess an ɑ/β hydrolase domain (Gomez-Valero *et al*., 2011; Gomez-Valero *et al*, 2014; Luo & Isberg, 2004), The structure of Lpg2422 has been experimentally determined (PDB 4m0m), confirming the presence of this domain; however, to the best of our knowledge, none of these effector proteins have been experimentally characterized to possess hydrolase activity. Along the same lines, we did not find any previous reports describing *Legionella* effectors possessing the SGNH hydrolase domain.

**Table 6.** The α/β-Hydrolase domain-containing effectors from *L. pneumophila* and their potential catalytic residues which were identified in this study.

| Effector | Potential, or Previously Described, Catalytic Residues | GDSL motif | Reference |
| --- | --- | --- | --- |
| Lpg0275/SdbA | His470-Cys300-Asp406 | N/A | This Study |
| Lpg1108 | His235-Ser125-Asp206 | N/A | This Study |
| Lpg1642 | His378-Ser190-Asp302 | N/A | This Study |
| Lpg1907 | His378-Ser239-Asp296 | N/A | This Study |
| Lpg1959 | His529-Cys321-Asp449 | N/A | This Study |
| Lpg2391 | His339-Ser188-Asp270 | N/A | This Study |
| Lpg2422 | His235-Ser144-Glu203 | N/A | This Study |
| Lpg2482/SdbC | His351-Cys187-Asp273 | N/A | This Study |
| Lpg2911 | His399-Ser165-Asp341 | N/A | This Study |
| Lpg0788 | His343-Ser73-Asp340 | Gly71-Asp72-Ser73-Leu74 | This Study |
| Lpg1354 | Glu242-Ser10-Ala239 | Gly8-Asp9-Ser10-Leu11 | This Study |
| Lpg2587 | His344-Ser17-Asp341 | Gly15-Asp16-Ser17-Leu18 | This Study |

Of the nine putative ɑ/β hydrolase domains, the ones found in six effectors (Lpg1108, Lpg1642, Lpg1907, Lpg2391, Lpg2422, Lpg2911) contain potential serine-aspartate-histidine catalytic triads, while the domains predicted in Lpg0275, Lpg1959, and Lpg2482/SdbC feature potential cysteine-aspartate-histidine triads (Fig. 6A). Our analysis indicates a diversity of lid subdomain structures among these putative ɑ/β hydrolases, while the models of Lpg1108, Lpg2391, and Lpg2422 lack the lid subdomain. The Lpg2911 effector shares significant similarity (33% identity) with human cathepsin A (PDB 4AZ0) (Ruf *et al*, 2012), which suggests that this effector may possess carboxypeptidase activity. The remaining 8 putative ɑ/β hydrolases show very low sequence identity with structurally and functionally characterized proteins. Some of the predicted features that may provide a cue to the function of these effectors include a long, deep, hydrophobic cleft in the model of Lpg1642 that is reminiscent of the ligand binding site in the monoacylglycerol lipase from *Palaeococcus ferrophilus* (PDB 6QE2) (Labar *et al*, 2021), and the model of Lpg2391 shows a broad, wide, and deep cleft.

**Figure 6.**
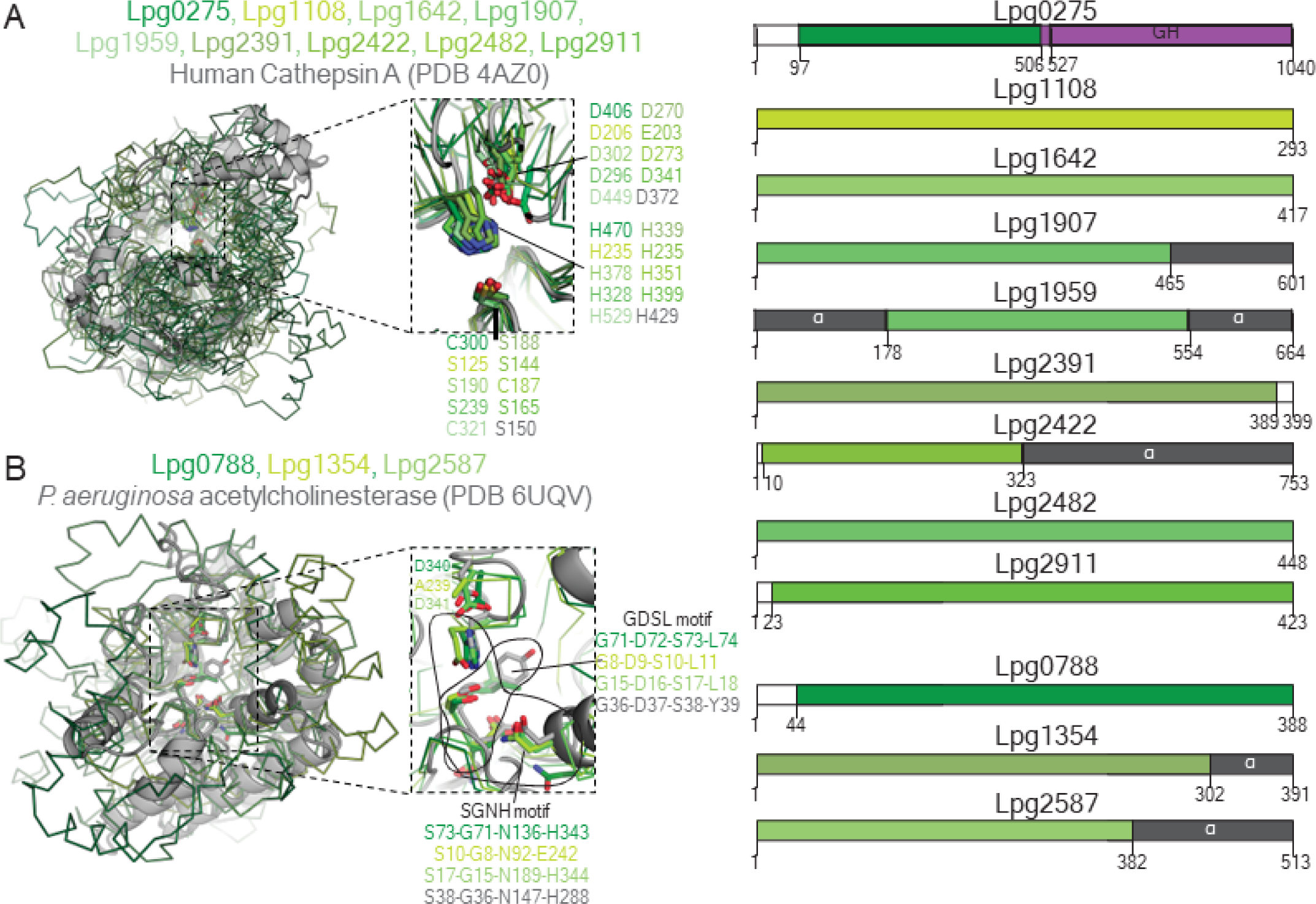
Model of *L. pneumophila* effector hydrolases. A. Putative effector hydrolases and (B) putative effector hydrolases in the SGNH class. For each panel, left = overlay of effector hydrolases (shades of green) with a characterized, reference structure from the PDB in grey. Right = Primary domain architecture, with the hydrolase domain colored in shades of green, alpha-helical regions colored grey, and glycoside hydrolase (GH) domain in Lpg0275 colored purple.

Lpg0788 and Lpg2587 contained the serine-glycine-asparagine-histidine tetrad typical for the catalytic site of the SGNH-type hydrolase enzymes (Fig. 6B). Notably, while the model of Lpg1354 adopts the ɑ/β hydrolase fold, the histidine in the catalytic site is replaced by aspartate-242, which brings into question whether this enzyme functions as a hydrolase. SGNH hydrolases are further subcategorized into GDSL esterases based on the presence of the corresponding sequence motifs in the N-terminal portion of the protein (Anderson *et al*., 2022). Accordingly, we identified such motifs for all three of these effectors represented by: residues 71-74 (GSDI) in Lpg0788; residues 8 to 12 (GDSTL) in Lpg1354; and 36 to 39 (GDSY) in Lpg2587. However, reliable prediction of the specific activity for effectors in this group is obscured by low similarity to characterized proteins.

### Predicted ADP-ribosyltransferase domains extend beyond LarT1, Ceg3, and the SidE family in the *L. pneumophila* effector arsenal

Adenosine diphosphate-ribosyltransferases, commonly referred to as ADP-ribosyltransferases, are a group of enzymes with crucial roles in various eukaryotic cellular processes (Luscher *et al*, 2018). These enzymes catalyze the transfer of ADP-ribose moieties from nicotinamide adenine dinucleotide (NAD^+^) onto target proteins (Mikolcevic *et al*, 2021; Suskiewicz *et al*, 2023). This post-translational modification can modulate protein activity, localization, and interactions within the cell (Suskiewicz *et al*., 2023). Similar domains have also been characterized as part of bacterial toxins and pathogenic factors, which harness this activity to disrupt critical host cell processes (Dean, 2011; Simon *et al*, 2014).

In *L. pneumophila,* ADP-ribosyltransferase domains have been characterized primarily as part of the paralogous SidE effector family (Lpg0234/SidE, Lpg2153/SdeC, Lpg2156/SdeB, and Lpg2157/SdeA). These effectors utilize their ADP-ribosyltransferase to ADP-ribosylate (ADPR) the Arg42 residue of ubiquitin, which is then conjugated onto the host substrate the effector’s phosphodiesterase (PDE) domain, forming a host protein-phosphoribosylate-ubiquitin complex (Bhogaraju *et al*, 2016; Dong *et al*, 2018; Kalayil *et al*, 2018; Kotewicz *et al*, 2017; Qiu *et al*, 2016; Song *et al*., 2021), as part of a two-step unorthodox ubiquitination mechanism. In addition to the SidE effector family, ADP-ribosyltransferase domains were also identified in Lpg0080/Ceg3 and Lpg0181. Lpg0080/Ceg3 is an ADP-ribosyltransferase involved in the modification of the Arg236 of the human adenine nucleotide translocase 2 (ANT2) - which is a membrane-spanning protein required for the exchange of ADP and ATP across the mitochondria inner membrane (Kubori *et al*, 2022). The modification of ANT2 has also been shown to be reversed by the metaeffector Lpg0081 (Kubori *et al*., 2022). Lpg0181 was reported to target a conserved Arg residue located within the NAD^+^ binding pocket of the 120 kDa glutamate dehydrogenase enzyme family that is present in both fungi and the protist hosts of *Legionella* (Black et al, 2021).

Based on our analysis, the models of two additional effectors - Lpg0796 and Lpg2523/Lem26 - also contain domains similar to an ADP-ribosyltransferase fold (Table 7). A previous study suggested that Lpg2523/Lem26 contains a C-terminal PDE domain based on primary sequence similarity to SdeA (Wan *et al*, 2019a). However, this study failed to demonstrate Lpg2523/Lem26’s ability to hydrolyze ADPR-Ubiquitin, suggesting that this effector’s PDE domain may have different substrate specificity (Wan *et al*., 2019a). Furthermore, the N-terminal (residues 5-328) of Lpg2523 has limited primary sequence identity to Lpg0080/Ceg3 where the sequence alignment implied the potential presence of functionally relevant residues involved in ADP-ribosyltransferase activity (Kubori *et al*., 2022). Our analysis of the Lpg2523 model revealed that apart from similarity to the PDE domain of SdeA, the N-terminal domain is reminiscent of the T3SS effector ExoT from *Pseudomonas aeruginosa* (Karlberg et al, 2018) and protein-arginine ADP-ribosyltransferase Tre1 (Ting *et al*, 2018) that was characterized as part of the T6SS arsenal in the insect pathogen, *Serratia proteamaculans* (Fig. 7A). In line with this analysis, the N-terminal domain of the Lpg2523 model features several functionally relevant elements identified in these bacterial effectors, including a catalytic triad consisting of an Arg222-Ser257-Glu294 residues, an ADP-ribosylating turn-turn (ARTT) loop harboring the Glu292-X-Glu294 motif essential for catalysis, followed by a β-sheet involved in the binding and stabilization of NAD^+^ (Fig. 7A and B). Overall, the structural analysis of the N-terminal domain of Lpg2523/Lem26 corroborates with the ADP-ribosyltransferase fold prediction.

**Figure 7.**
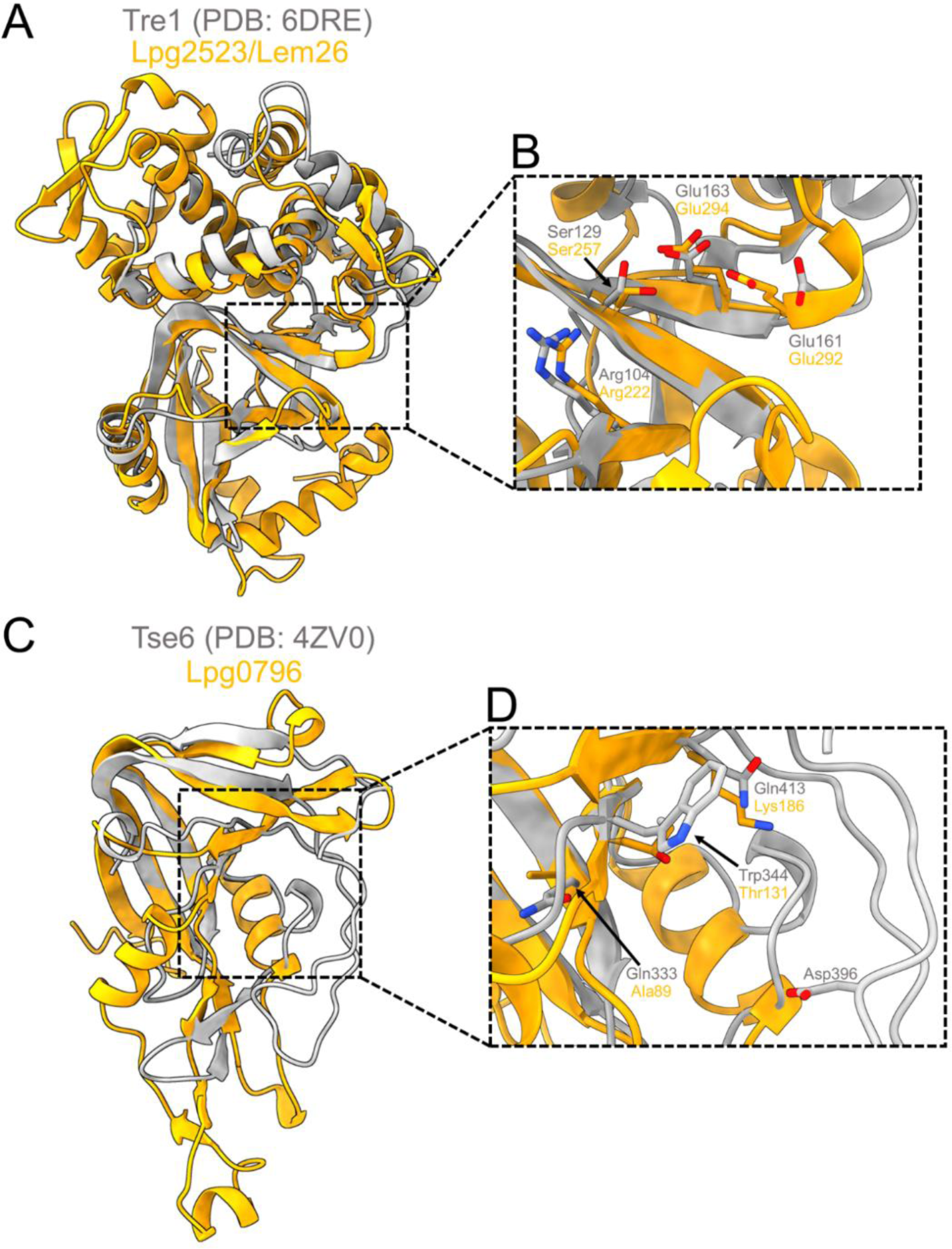
Uncovering cryptic ADP-ribosyltransferase folds in two *L. pneumophila* effectors. A. An overall structural alignment of Lpg2523/Lem26 (residues 5-325) with the structurally characterized ADP-ribosyltransferase effector Tre1 from *S. proteamaculans* (residues 8-192) (Ting *et al*., 2018). B. Closer view of the corresponding putative catalytic residues (gold sticks) of Lpg2523/Lem26 assigned by the structural alignment with Tre1 (grey sticks) from (A). C. Structural alignment between Lpg0796 (residues 71-208) and the T6SS effector, Tse6 (residues 318-427), from *P. aeruginosa* (Whitney et al., 2015). D. Inset view comparing the catalytic residues of Tse6 (grey sticks) with the corresponding residues of Lpg0796 (gold sticks).

**Table 7.**
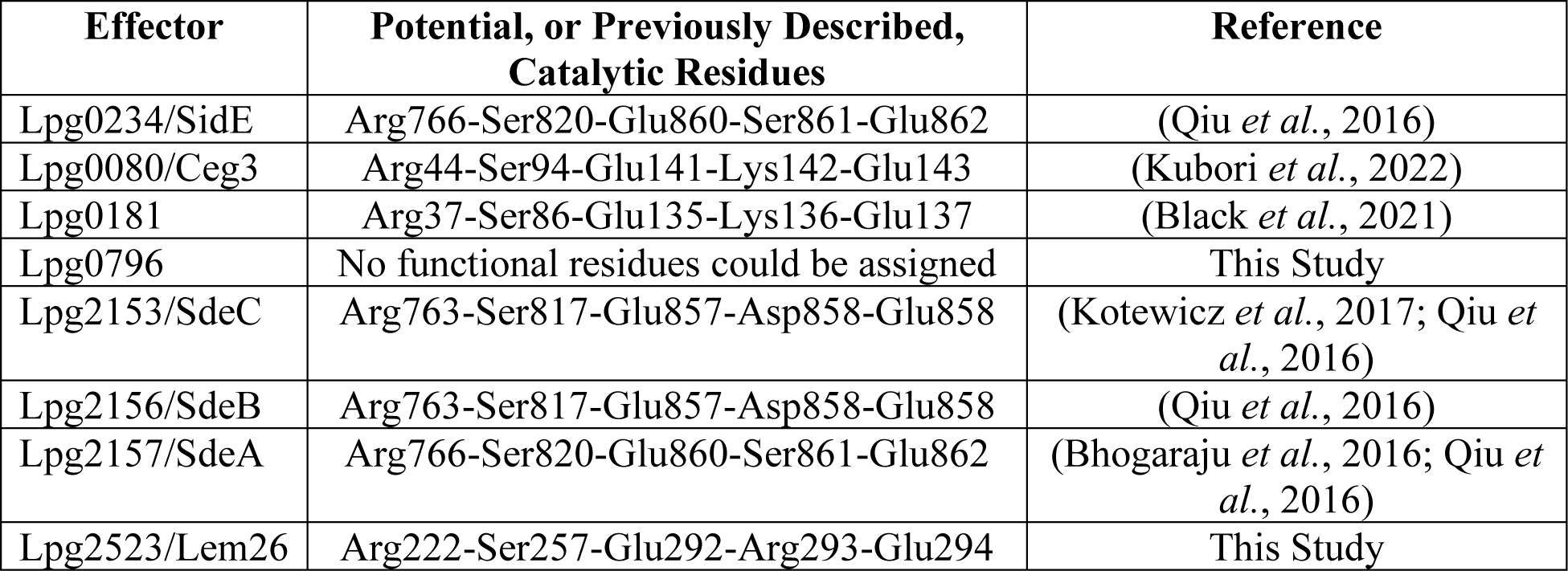
Identification of *L. pneumophila* ADP-ribosyltransferase domain-containing effectors and their catalytic residues that were identified in this study and previous ones.

The analysis of the predicted ADP-ribosyltransferase domain in the Lpg0796 model suggested a structural resemblance to Tse6 - a T6SS effector from *P. aeruginosa* (Fig. 7C). Tse6 is an effector that is structurally similar to the catalytic domain of ADP-ribosyltransferase toxins released by human bacterial pathogens - such as the diphtheria toxin from *Corynebacterium diphtheriae* and Exotoxin A from *P. aeruginosa* (Whitney et al, 2015). However, despite having this enzymatic domain, Tse6 has diverged in function by acting as a NAD(P)+ glycohydrolase (Whitney *et al*., 2015). Our structural analysis of the Lpg0796 model suggests that it possesses a similar conserved ꞵ-sheet core involved in NAD+ binding. However, we were not able to identify corresponding residues in the ꞵ-sheet core that could contribute to the interaction of this co-factor. Furthermore, the ability of Tse6 to hydrolyze NAD(P)^+^ is facilitated by Asp396 in an activation loop - a structural element that is absent in Lpg0796 (Whitney *et al*., 2015) (Fig. 7D). Taken together, we suggest that Lpg0796 lacks catalytically important residues typical of ADP-ribosyltransferases and NAD(P)^+^ glycohydrolases, suggesting a possible diversification of its biochemical activity.

### Five additional *L. pneumophila* effectors contain potential glycosyltransferase domains

Glycosyltransferases facilitate the transfer of a glycosidic sugar moiety from a donor co-substrate to an acceptor co-substrate, such as nucleic acids, lipids, and proteins (Zhang *et al*, 2020). Based on the features of their primary sequence, glycosyltransferases form more than a hundred distinct families and adopt one of the three different structural folds, known as GT-A, GT-B, and GT-C (Rosen *et al*, 2004). Enzymes classified as GT-A possess a Rossman-like fold with a conserved aspartate-X-aspartate motif required for the coordination of divalent cations and transferase activity (Persson *et al*, 2001; Taujale *et al*, 2020). In the case of the GT-B domain, such enzymes contain two Rossman-like folds that form a central cleft, serving as the site for catalysis (Breton *et al*, 2006). Furthermore, GT-B enzymes do not require divalent cations for catalysis, thus, lacking the aspartate-X-aspartate motif (Both *et al*, 2011; Li *et al*, 2007). Instead, a single negatively charged (aspartate or glutamate) residue, typically found in the N-terminal Rossman-like fold, has been demonstrated to be essential for catalysis. The C-terminal Rossman-like fold is involved in the recognition and binding to the donor co-substrate. GT-C fold-containing enzymes have a single Roseman-like fold connected to multiple transmembrane helices and use lipid-linked sugars as the donor co-substrate (Alexander & Locher, 2023; Rini *et al*, 2009).

Several *L. pneumophila* effectors have been demonstrated as GT-A and GT-B glycosyltransferases, including the members of the Lgt effector family (Lpg1368/Lgt1, Lpg2862/Lgt2, and Lpg1488/Lgt3). The structural characterization of Lpg1368/Lgt1 revealed that this effector family harbors a GT-A domain involved in the glycosylation of the human elongation initiation factor A1, which causes protein translation to be inhibited (Belyi *et al*, 2009; Belyi *et al*, 2008; Lu *et al*, 2010). Lpg1978/SetA also contains a GT-A domain that was shown to have activity against the human transcription factor EB, histones H3.1 and H4 in *in cellulo* and *in vitro* assays; however, the exact role of this effector during infection remains unclear (Beck *et al*, 2020; Jank *et al*, 2012). Another GT-A domain-containing effector is Lpp0356/LtpM which has an atypical active site architecture, consisting of an aspartate-X-asparagine motif required for catalysis (Levanova *et al*, 2019). The substrate specificity of this effector remains unknown; however, it has been postulated that LptM is involved in hijacking the microtubule vesicle trafficking pathway (Levanova *et al*., 2019). Lpg2504/SidI is an effector with a GT-B fold that acts as a mannosyltransferase on host ribosomes, resulting in the inhibition of protein translation and the activation of host stress response kinases that promote the transcription of genes involved in cell death (Joseph *et al*, 2020; Subramanian *et al*, 2023).

Our analysis of the *L. pneumophila* effector models identified potential glycosyltransferase domains in five additional effectors (Lpg0275/SdbA, Lpg0402/LegA9, Lpg0770, Lpg1151, and Lpg1961) (Table 8). Lpg1961 has a structural resemblance to the GT-A domain containing PaToxG toxin from the insect and human bacterial pathogen, *Photorhabdus asymbiotica* (Costa et al, 2009; Jank et al, 2013), which is reflected by the superimposition having an RMSD of 2.89 Å over 205 Cα equivalent positions. The model of Lpg1961 has an active site strikingly similar to PaToxG (Jank *et al*., 2013) and contains a typical aspartate-X-aspartate motif, including several residues (Trp33, Asp130, and Arg133) possibly involved in the interaction and transfer of the donor co-substrate (Fig. 8A). For the Lpg0275/SdbA, Lpg0402/LegA9, Lpg0770, and Lpg1151 effectors, we determined that these effectors are likely members of the GT-B family with two Rossman-like folds indicative of glycosyltransferase activity (Fig. 8B). We also defined that these effectors likely harbour potential catalytic residues in their N-terminal Rossman-like fold and use their C-terminal fold for nucleotide binding, which is consistent with the functionality of most GT-B enzymes (Table 8). Overall, we have determined new effectors with glycosyltransferase folds; however, a limitation of our modeling analysis prevents us from determining the specific donor co-substrate involved in the activity of these effectors.

**Figure 8.**
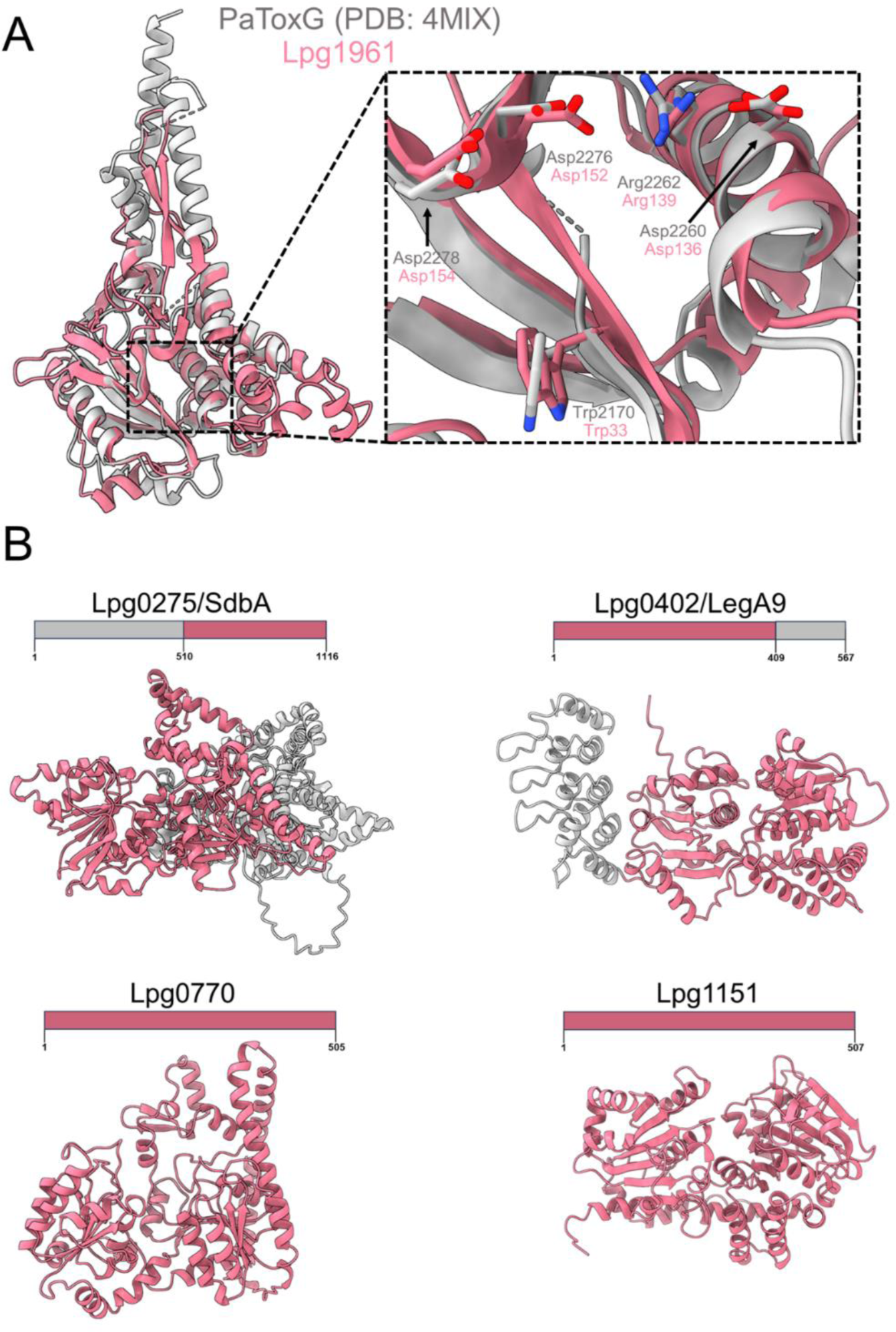
Identification of five more members of *L. pneumophila* effectors with glycosyltransferase domains. A. Structural alignment of Lpg1961 GT-A domain (residues 27-321) with the *Photorhabdus asymbiotica* toxin PaToxG (residues 2131-2421) (Jank *et al*., 2013). A close view of the potential active site of Lpg1961 where pink sticks indicate putative catalytic residues determined by the PaToxG alignment (grey sticks). B. Structural domain architecture of *L. pneumophila* effectors with a GT-B domain (pink) while other structural features not involved in potential glycosyltransferase activity are highlighted in grey.

**Table 8.**
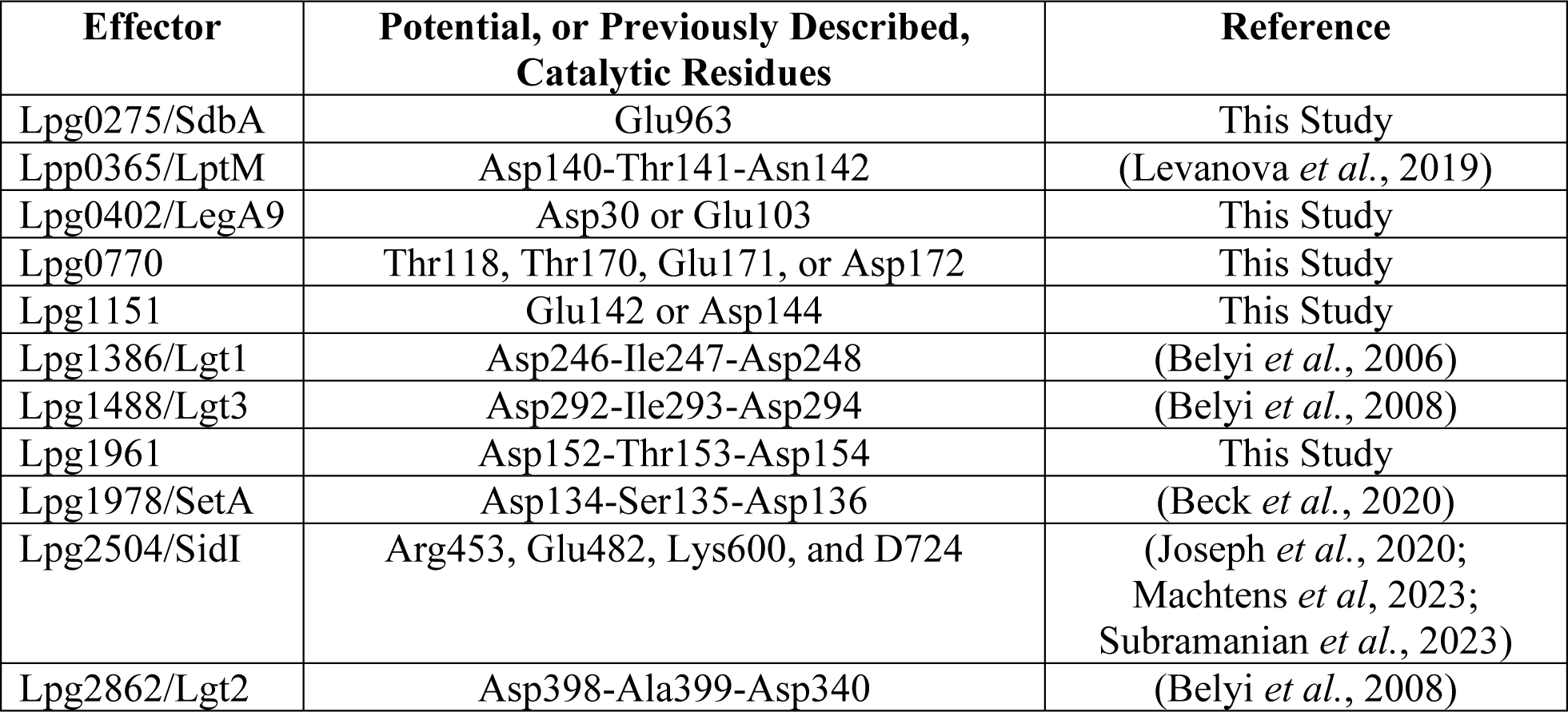
List of effectors with a glycosyltransferase domain and their catalytic residues determined from our 3D modeling analysis in this study or others. The effectors are either from *L. pneumophila* (Lpg) or *L. pneumophila* Paris strain (Lpp).

| Effector | Potential, or Previously Described, Catalytic Residues | Reference |
| --- | --- | --- |
| Lpg0275/SdbA | Glu963 | This Study |
| Lpp0365/LptM | Asp140-Thr141-Asn142 | (Levanova *et al.*, 2019) |
| Lpg0402/LegA9 | Asp30 or Glu103 | This Study |
| Lpg0770 | Thr118, Thr170, Glu171, or Asp172 | This Study |
| Lpg1151 | Glu142 or Asp144 | This Study |
| Lpg1386/Lgt1 | Asp246-Ile247-Asp248 | (Belyi *et al.*, 2006) |
| Lpg1488/Lgt3 | Asp292-Ile293-Asp294 | (Belyi *et al.*, 2008) |
| Lpg1961 | Asp152-Thr153-Asp154 | This Study |
| Lpg1978/SetA | Asp134-Ser135-Asp136 | (Beck *et al.*, 2020) |
| Lpg2504/SidI | Arg453, Glu482, Lys600, and D724 | (Joseph *et al.*, 2020; Machtens *et al.*, 2023; Subramanian *et al.*, 2023) |
| Lpg2862/Lgt2 | Asp398-Ala399-Asp340 | (Belyi *et al.*, 2008) |

### Cryptic domain activity of *L. pneumophila* effectors manifests in the yeast model system

Our and others’ previous work has demonstrated that the ectopic expression of individual *L. pneumophila* effector proteins in the *Saccharomyces cerevisiae* model system can result in different degrees of growth defects (Urbanus *et al*., 2016). This phenomenon has been exploited for the elucidation of the biochemical activity of several effectors in *L. pneumophila* (Belyi et al., 2012; Bhogaraju et al., 2016; Campodonico et al., 2005; de Felipe et al., 2008; Fu et al., 2022; Gaspar & Machner, 2014; Guo et al., 2014; Qiu et al., 2016; Shohdy et al., 2005; Urbanus et al., 2016; Viner et al., 2012), but for most effectors, the cause of their toxicity for *S. cerevisiae* remains unknown. Identification of potential cryptic functional domains in the 3D models of these uncharacterized effectors provided us with the opportunity to experimentally test the role of these domains in the observed yeast growth defect phenotype.

Our analysis identified cryptic functional domains in eleven *L. pnemophila* effectors that were previously demonstrated to cause significant toxicity in yeast (Fig. EV1A-K). Correspondingly, we probed the role of individual residues in the active sites of these cryptic domains by site-directed mutagenesis (Fig. 9A-D, Appendix Fig. S4-5). The expression of Lpg1290/Lem8 was used as a control for this experiment. The toxicity of this effector to yeast was linked to its cysteine protease domain and was shown to be alleviated by Cys280Ser, His391Ala, or Asp412Ala substitutions of catalytic triad residues (Song *et al*, 2022).

**Figure 9.**
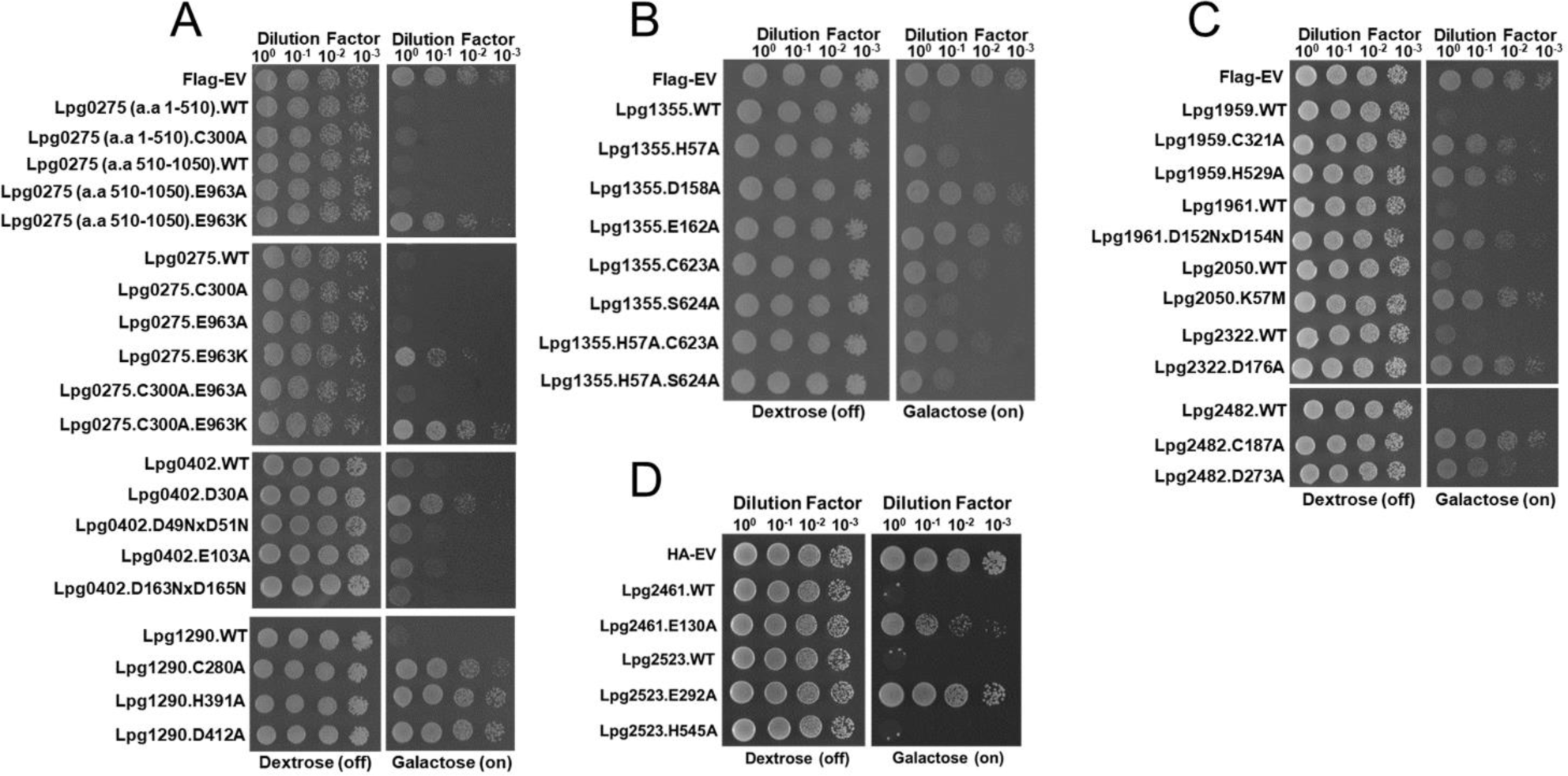
Yeast toxicity panel of *L. pneumophila* effectors harboring cryptic enzymatic domains. A-C. The overexpression of *L. pneumophila* effectors (Lpg0275 to Lpg2482) in *S. cerevisiae* strain BY4741 using the wild-type and mutant constructs with point mutations targeting a specific cryptic domain in Supplemental Table 2 and Figure EV1. D. Overexpression of Lpg2461 and Lpg2523 and their respective mutants using the low copy number pGAL416-HA plasmid in the *S. cerevisiae* strain BY4741. Yeast toxicity panel experiments in A-D were spotted with the indicated serial dilutions on synthetic defined (SD) media containing either dextrose (repressing) or galactose (inducing). Experiments were also repeated three times independently with similar results. Each mutant in A-D was tested for expression, solubility, and stability via a western blot using an antibody targeting the respective epitope tag (Appendix Figures S4 and S5)

The first effector we tested was Lpg0275/SdbA, where the model suggested the presence of two distinct functional domains connected by a flexible linker (Fig. EV1A). Residues 1 to 510 of Lpg0275/SdbA are predicted to form an ɑ/β hydrolase domain, while residues 511 to 1050 are predicted to form a domain adopting a fold reminiscent of GT-B glycosyltransferases (Fig. EV1A). To test if each of these predicted domains contributed to toxicity in yeast, we expressed the corresponding fragments of Lpg0275 individually in yeast along with the variants carrying substitutions in putative catalytic residues in each of the predicted domains. Based on our results, the individual expression of each of the two predicted domains in Lpg0275 causes toxicity in yeast (Fig. 9A) suggesting that both domain activities contribute to this phenotype. The Cys300Ala substitution of the ɑ/β hydrolase domain significantly reduced the toxicity compared to the wild-type expression (Fig. EV1A and Fig. 9A). The lysine substitution of Glu963 also partially alleviated the growth defect caused by the expression of Lpg0275 [511-1050] fragment (Fig. 9A). Notably, the alanine substitution of this residue was insufficient to alleviate the toxicity of this fragment. In agreement with these results, the Glu963Lys mutation in the context of full-length Lpg0275 also partially alleviated toxicity, whereas the double substitution of Cys300Ala and Glu963Lys restored the growth of yeast comparable to that of the FLAG-only yeast control (Fig. 9A).

Next, we targeted potential catalytic residues in the glycosyltransferase (GT) domains predicted in Lpg0402 and Lpg1961 effectors. In the case of the former effector, we identified Asp30 and Glu103 as residues that correspond to the catalytically important residues in experimentally characterized members of this protein family (Fig. EV1B and Fig. EV1F). Furthermore, our analysis suggested that Lpg0402 Asp49/Asp51 or Asp163/Asp165 pairs can form catalytically important Asp-X-Asp motif typically found in GT enzymes (Fig. EV1B). The yeast toxicity assay suggests that the alanine substitution of Asp30 completely alleviates the toxicity of Lpg0402 which is in agreement with the suggested role of this residue in the predicted GT-B domain of this effector (Fig. 9A). In contrast, Lpg0402 variants carrying Asp49Gln/Asp51Gln or Asp163Gln/Asp165Gln double mutations had a yeast toxicity profile comparable to the wild type effector.

Similarly, our analysis of the Lpg1961 model suggested residues Asp152 and Asp154 as candidates for theAsp-X-Asp motif essential for the catalytic activity of its predicted GT-A domain (Fig. EV1F). The double substitution of these residues to asparagine led to complete alleviation of toxicity, thus, corroborating our structural prediction analysis and suggested role of this predicted domain in the observed phenotype (Fig. 9C).

We next tested the role of the ɑ/β hydrolase domains predicted in the Lpg1959 and Lpg2482/SdbC effectors (Fig. EV1E and Fig. EV1J). In the case of Lpg1959, we predicted that the residues Cys321, Asp346, and His529 to fulfill the role of a catalytic triad (Table 7). In support of this, alanine substitution of either Cys321 or His529 resulted in the recovery of yeast growth, clearly implicating the activity of this cryptic domain in toxicity (Fig. 9C and Figure EV1E). Similarly, alanine substitution of either Cys107 or Asp273 identified as potential catalytic residues in Lpg2482/SdbC also resulted in the alleviation of toxicity (Fig. 9C, Fig. EV1J). The observed effect was stronger in the case of the former residue substitution (Fig. 9C).

Our analysis suggested the presence of a potential kinase domain in Lpg2050 and Lpg2322/AnkK effectors causing toxicity in yeast (Fig. EV1G-H). Based on our analysis, we suggested that Lys57 is catalytically important for Lpg2050 (Fig. 9C and Fig. EV1G). Substitution of Lys57 to methionine restored the yeast growth to that of the FLAG-only control yeast (Fig. 9C). A previous study used primary sequence analysis to suggest that His178 makes part of the kinase activation loop of Lpg2322/AnkK (Ledvina *et al*., 2018). Extending on this analysis, we tested the role of Asp176 also predicted to be part of the activation loop in the model’s kinase domain (Fig. EV1H). In line with its suggested role, the Asp176Ala substitution led to the abrogation of Lpg2322/AnkK toxicity in yeast (Fig. 9C).

The model of the Lpg1355/SidG effector suggested the presence of a cryptic cysteine protease domain, with residues Cys623, Asp158, and His57 forming the catalytic triad that is typical of such enzymes (Fig. 3B). The alanine substitution of Asp158 led to the complete alleviation of toxicity, whereas similar substitutions of His57 or Cys623 led to a partial alleviation of the yeast growth defect (Fig. 9B). Furthermore, our analysis of the Lpg1355/SidG model suggested that the residues Glu162 and Ser624 may also be important for catalytic activity of this domain (Fig. 3B and Fig. EV1D). In line with our hypothesis, the alanine substitution of Glu162 led to complete alleviation of Lpg1355/SidG’s toxicity, while a similar substitution to the latter residue failed to restore yeast growth (Fig. 9B). Next, we tested the effect of double His57Ala/Cys623Ala or His57Ala/Ser624Ala substitutions on the toxicity of Lpg1355/SidG. In the case of the His57Ala/Ser624Ala variant, we observed partial restoration of growth comparable to the effect observed for the His57Ala variant. For the His57Ala/Cys623Ala Lpg1355 variant, we also observed restoration of yeast growth which appeared stronger than what we observed in the case of the His57Ala or Cys623Ala Lpg1355/SidG variants (Fig. 9B).

We identified a cryptic domain reminiscent of zinc metalloproteases in the model of the Lpg2461 effector, forming the classical catalytic motif present in these enzymes. Confirming this analysis, alanine substitution of Glu130 in Lpg2461 resulted in the complete restoration of yeast growth (Fig. EVJ and Fig. 9D).

The model of Lpg2523/Lem26 revealed two distinct domains connected by a central helical bundle. The N-terminal domain spanning residues 1 to 337 resembles an ADP-ribosyltransferase domain, while the model of the C-terminal portion of this effector that spans residues 494 to 779 shares structural similarity to the PDE domain of the SidE effector family (Fig. EV1K). To test if either of these domains contributes to the toxic effect of Lpg2523 in yeast cells, we targeted Glu294 residue, which is suggested to be part of the Glu-X-Glu motif in the ARTT loop of the ADP-ribosyltransferase domain, and His545 as a putative catalytic residue in the PDE domain (Fig. EV1K). The Glu294Ala mutation resulted in restored yeast growth, while the expression of His545Ala Lpg2523/Lem26 variant showed toxicity similar to that of the wild-type effector. These results suggested that while the function of PDE domain of Lpg2523/Lem26 remains enigmatic, the predicted ADP-ribosyltransferase domain is responsible for the toxicity phenotype in yeast (Fig. 9D).

### *L. pneumophila* effector models contain significant number of domains with unique folds, some of which are responsible for toxicity in yeast

A number of predicted structural domains in *L. pneumophila* effector models demonstrate no significant structural similarity to the experimentally defined protein structures deposited to either the ECOD domain or the PDB databases, when analyzed with standard structure comparison tools, such as Dali (Holm, 2022) and FATCAT (Ye & Godzik, 2003). Accordingly, we classify these domains as having “unique folds” (Table 9), even that manual analysis can identify some commonalities with known folds. We discuss such cases in detail below. Overall, we identified 35 such domains in 30 effectors, with models of five effectors –Lpg1426/VpdC, Lpg1978/SetA, Lpg1925/CegL1, Lpg1963, and Lpg1964, – containing two unique domains each (Table 9). In fourteen effectors, the unique folds were the only globular domain identified in the models, while in the case of six effector models the unique fold was accompanied by an additional ɑ-helical domain that could not be confidently matched to any similar element in experimentally structurally characterized proteins (Table 9 and Supplemental Table 1).

**Table 9.** Residue boundaries of the unique folds in *L. pneumophila* effectors and the primary sequence similarity to the closest structural homolog outside of the *Legionella* genus.

| **Effector** | **Amino acid boundary of the Unique fold (s)** | **Domains classified in ECOD (for details see Table 1 and Sup. Table 1)** | **Closest Homolog to the Unique Fold** | **Sequence Similarity to Homolog (%)** |
| --- | --- | --- | --- | --- |
| Lpg0012 | 1-474 | - | GGR11_001436 from *Brevundimonas mediterranea* | 20.7 |
| Lpg0172 | 1-109 | Hom (152-211) | BHQ10_008315 from *Talaromyces amestolkiae* | 24.0 |
| Lpg0519 | 179-285 | - | AQUSIP_07400 from *Aquicella siphonis* | 48.8 |
| Lpg0717 | 24-153 | - | AQUSIP_07790 from *Aquicella siphonis* | 52.5 |
| Lpg0733 | 159-272 | LPG2148ins (273-364) | N/A | N/A |
| Lpg1083/SidN | 1-227 | - | CULFYP111_01230 from *Campylobacter ureolyticus* | 53.3 |
| Lpg1154/RavQ | 56-360 | - | AQUSIP_02370 from *Aquicella siphonis* | 55.5 |
| Lpg1426/VpdC | 1-239 and 271-299+623-678 | PhosphoLip (300-622) | FDP41_009687 from *Naegleria fowleri* | 16.2 |
| Lpg1489/RavX | 93-288 | - | N/A | N/A |
| Lpg1551 | 143-241 | - | N/A | N/A |
| Lpg1588 | 165-479 | - | N/A | N/A |
| Lpg1692 | 283-421 | Fbox (1-63), Ank (70-140+268-282), and LPG2148ins (141-267) | orf 309 from *Coxiella burnetii* | 33.0 |
| Lpg1716 | 1-103 | - | ROA7023_03573 from *Roseisalinus antarcticus* | 6.50 |
| Lpg1925 | 110-320 and 321-455 | DnaD (1-109) and Ank(529-821) | GRS96_03500 from *Rathayibacter* sp. VKM Ac-2803 | 11.7 |
| Lpg1963 | 187-320 and 522-681 | - | N/A | N/A |
| Lpg1964 | 60-225 and 226-404 | - | N/A | N/A |
| Lpg1978/SetA | 282-450 and 515-628 | NucDiStran (1-223) |  |  |
| Lpg2073 | 250-380 | Imm (5-249) and HEPN(381-535) | BGO78_14705 from *Chloroflexi* bacterium 44-23 | 44.6 |
| Lpg2160 | 1-332 | - | IMCC3317_17790 from *Kordia antarctica* | 14.9 |
| Lpg2207 | 263-407 | LDtrans (43-233) | A3F10_03715 from *Coxiella burnettii* | 65.9 |
| Lpg2223 | 1-384 | - | N/A | N/A |
| Lpg2239 | 513-730 | LPG2422hbun (1-153), SidE2 (158-460), Ank (731-1000), and Ank(1001-1285) | N/A | N/A |
| Lpg2372 | 279-422 | CheY (107-278) | N/A | N/A |
| Lpg2385 | 1-156 | PInPhC (466-595) | N/A | N/A |
| Lpg2410/vpdA | 23-51+404-459 | PhosphoLip (53-403) and WYR (513-655) | COA94_02160 from *Rickettsiales* bacterium (unclassified) | 22.7 |
| Lpg2452/LegA14 | 1-202+309-450 | EF (208-299) and Ank (485-921) | N/A | N/A |
| Lpg2504/SidI | 52-137+228-247 | UDPGluco(1-51+427-568) | N/A | N/A |
| Lpg2527/LnaB | 94-311 | - | A7J50_5968 from *Pseudomonas antarctica* | 26.8 |
| Lpg2638 | 1-280 | - | N/A | N/A |
| Lpg2912 | 1-122 | AcylC (123-343) | N/A | N/A |

Primary sequence analysis of effectors with predicted unique folds suggested that these are not exclusive to the *L. pneumophila* proteome. Most of the effectors from this category are conserved across *Legionella* species, with six effectors also carrying strong similarity to proteins present in the intracellular pathogens from *Coxiellacae* family, such as *Aquiclla siphonis*, *Coxiella burnetii*, and *Rickettsiella* spp (Table 9).

Importantly, the structures of three effectors – Lpg1083/SidN, Lpg1978/SetA and Lpg2504/SidI with predicted unique folds were experimentally determined and deposited in the PDB database (PDB: 7YJI, PDB: 7TOD, and PDB: 8BVP, respectively) during the course of our analysis, confirming the structural predictions (Gao *et al*, 2023; Subramanian *et al*., 2023) (Beck *et al*, 2022) and novel fold assignments. Interestingly, manual analysis of these effector structures identified potential distant structural similarities, providing the first indication of their molecular function (Table 9). In the case of Lpg1978/SetA, the unique fold is present in the C-terminal domain and consists of an ɑ-helical bundle connected to a β-sheet that forms a positively charged pocket important for interactions with phosphoinositol-3-phosphate (Beck *et al*., 2022). This domain with a unique fold is essential for the localization of Lpg1978/SetA on the surface of the LCV (Beck *et al*., 2022). The Lpg1083/SidN structure was revealed to be a unique domain where the N-terminal region, described to be “paw-like”, aids in the localization of the effector to the nucleus (Gao *et al*., 2023). Furthermore, Lpg1083/SidN is shown to disrupt the lamina complex that, in turn, leads to the destabilization of the nuclear envelope (Gao *et al*., 2023).

Awaiting structural characterization of remaining effector proteins with unique predicted structural elements, we used the yeast toxicity model to investigate their functional relevance in Lpg1154/RavQ, Lpg1426/VpdC, Lpg1489/RavX, or Lpg2527/LnaB (Li *et al*, 2022; Urbanus *et al*., 2016). Detailed analysis of the structural models combined with primary sequence conservation suggested residues that may be part of the active site in these proteins (Fig. 10A-D, Fig. EV2-5). Accordingly, we probed their relevance for toxicity using site-directed mutagenesis (Fig. 10E).

**Figure 10.**
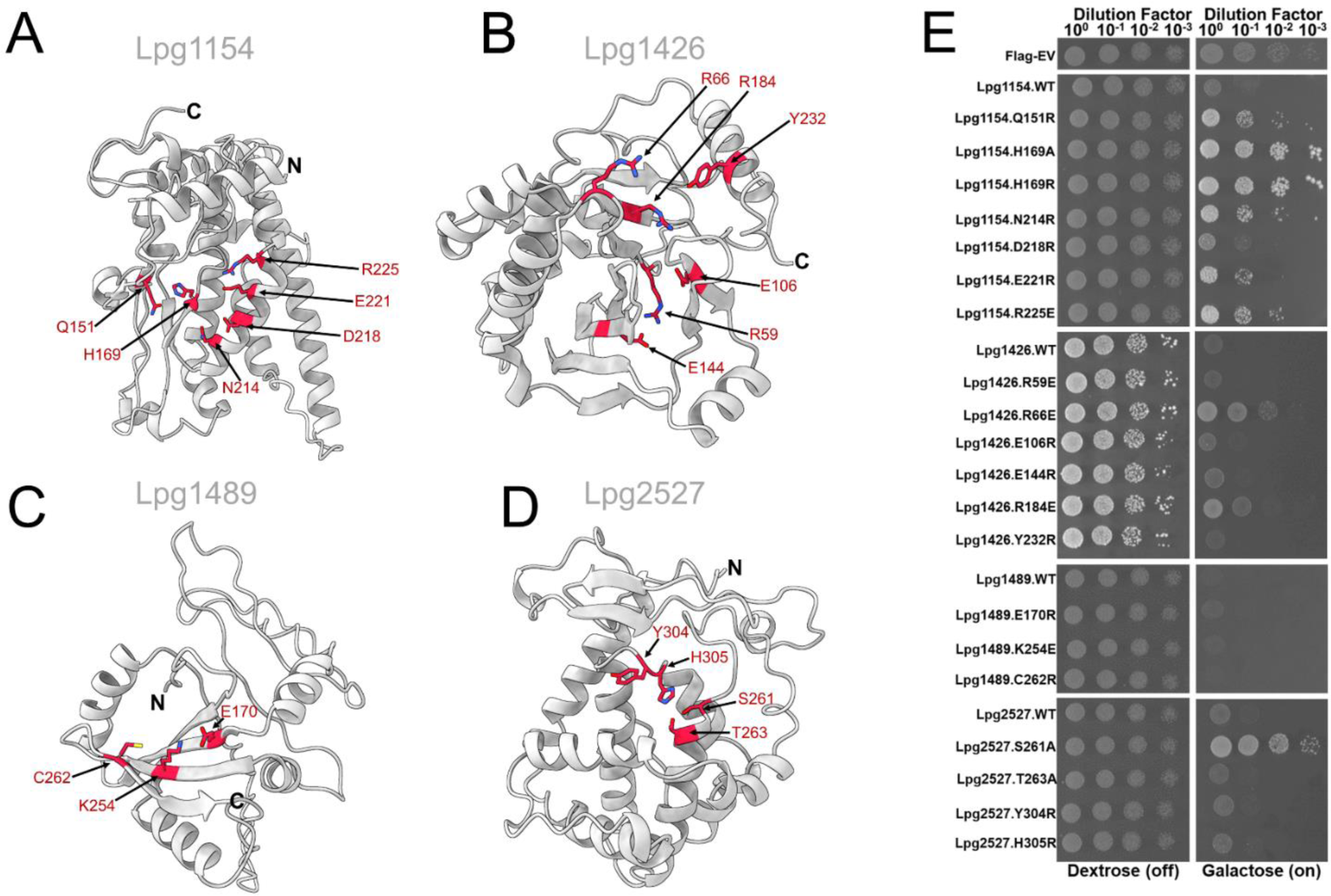
Unique folds in *L. pneumophila* effectors are capable of contributing to the yeast toxicity phenotype. A-D. Structural models of the unique fold from *L. pneumophila* effectors that are toxic to yeast when expressed ectopically (grey). Potentially important residues in the unique folds that were mutated and tested in (E) were shown in sticks (red). E. Constructs of the novel fold wild-type effectors and their respective mutants that were overexpressed in *S. cerevisiae* strain BY4741. Serial dilutions were spotted on synthetic defined media containing either dextrose (repressing) or galactose (inducing). The yeast toxicity panel was performed in triplicate with similar results. The expression levels of each of the mutants tested were analyzed using western blot (Appendix Figure S6).

Based on the structural model of Lpg1154/RavQ, the residues 59 to 349 form a unique α/β domain, while the preceding N-terminal portion of this effector is predicted to be disordered. This unique domain is predicted to contain a three-stranded β-sheet with a two-stranded β-sheet packed onto one face. The other face of the three-stranded β-sheet is packed against a four-helix bundle, and the model also contains five other ɑ-helices. The overall shape of the model is a “T” shape, with the base of the shape formed by an N-terminal ɑ-helix in one direction and the C-terminal three ɑ-helices in the other direction (Fig. EV2A). One of the vertices of the “T” shape is lined up with negatively and positively charged residues at its base and on its side, respectively (Fig. EV2A). According to the comparative sequence analysis using ConSurf server (Ben Chorin *et al*, 2020), the residues forming this groove show complete conservation across orthologs found in other *Legionella* species (Fig. EV2B). Specifically, a highly conserved histidine residue (His169 in Lpg1154/RavQ) is positioned at the center of the groove surrounded by other conserved residues, including Asn141, Gln151, Asp218, Glu221, and Arg225 (Fig. EV2B). Accordingly, we tested all six highly conserved residues identified to be potentially relevant to Lpg1154/RavQ activity by mutagenesis. Substitution of His169 to alanine, or arginine, led to complete alleviation of yeast toxicity (Fig. 10D and Fig. 10E). The substitution of Glu151 and Asn214 to arginine residues partially alleviated the toxicity of Lpg1154/RavQ (Fig. 10E). In contrast, substitution of Asp218 and Glu221 to arginine did not rescue yeast growth, suggesting that these substitutions are not detrimental to effector’s activity in this model system (Fig. 10E). Notably, we also identified orthologs of Lpg1154 in more distant members of the *Legionellales* order; for instance, *Aquicella siphonis* from the *Coxiellaceae* family but also intracellular pathogens from the *Chlamydiales* order, such as *Waddlia chondrophila* or *Estrella lausannensis* (Supplemental Table 3).

The model of Lpg1426/VpdC effector contained three distinct domains: a unique fold spanning residues 1 to 299, a lysophospholipase domain (E-Cod T group FabD/lysophospholipase-like, Patatin Family) spanning residues 299 to 621, and a small ɑ/β structure that packs onto the second unique domain (residues 622-677) followed by a predicted disordered region (residues 678-719), and a C-terminal helical bundle corresponding to residues 720 to 853 (Fig. EV3A). A recent report indicated that this effector relies on the C-terminal helical bundle to bind ubiquitin, which, in turn, results in a conformational change to activate its phospholipase domain to facilitate the conversion of phospholipids into lysophospholipids (Li *et al*., 2022). The activity of Lpg1426/VpdC was shown to be important for LCV expansion during the infection of U937 human macrophages (Li *et al*., 2022). The predicted domain with a unique fold in Lpg1426/VpdC contains a central seven-stranded β-sheet which is bounded by eight ɑ-helices on one face of the β-sheet, and a small region containing a two-stranded β-sheet and one ɑ-helix on the other face (Fig. EV3A). Further analysis of this unique domain suggested it may be distantly homolog with the Ntox11 domain, a structurally uncharacterized putative toxin domain that is broadly distributed in bacterial and some eukaryotic pathogens (Fig. EV3B) (Zhang *et al*, 2012). Interestingly, the latter group includes *Naegleria fowleri* amoeba species, which has been identified as one of the natural protist hosts of *Legionella* (Fig. EV3B) (Boamah *et al*, 2017; Fields, 1996; Newsome *et al*, 1985). Our analysis of this unique fold domain in Lpg1426/VpdC suggests the formation of a negatively-charged groove formed by highly conserved residues - Arg59, Arg66, E106, E144, Arg184, and Tyr232 – present in Lpg1426/VpdC orthologs encoded by the other *Legionella* species (Fig. EV3C and Fig. EV3D). Our mutational analysis demonstrated that glutamate substitutions of Arg66 or Arg184 partially attenuate the toxic effect of Lpg1426/VpdC in yeast which is in line with the suggested role in the predicted active site of the unique fold domain (Fig. 10B and Fig. 10E).

With Lpg1489/RavX, a previous report identified this protein to be a member of a large group of effectors (Lpg0103/VipF, Leg0208/LegK4, Lpg0437/Ceg14, Lpg1368/Lgt1, Lpg1488/Lgt3, Lpg2504/SidI, and Lpg2862/Lgt2) shown to manipulate eukaryotic protein translation (Barry *et al*, 2013; Belyi *et al*, 2006; Belyi *et al*., 2008; Fontana *et al*, 2011; Joseph *et al*., 2020; Moss *et al*., 2019; Subramanian *et al*., 2023; Syriste *et al*, 2024). However, the structural and biochemical basis of this process remains undefined. Our analysis of the structural model of Lpg1489/RavX suggested the presence of a domain with a unique fold encompassing residues 89 to 267. However, the AlphaFold2 model of Lpg1489/RavX had low (<50%) confidence, which can be explained by the fact that this effector is found only in *L. pneumophila*. To overcome the potential limitation of our modelling approach using Alphafold2, we generated another structural model using ESMFold, an LLM-based algorithm that does not rely on MSA for the prediction (Lin *et al*., 2023). The ESMFold-generated model of Lpg1489/RavX was similar to the unique fold suggested by Alphafold2 but had higher confidence (Fig. EV4A), and thus was selected for further analysis. Interestingly, the structural assembly predicted in this model showed distant similarity to the structure of *E. coli* enterotoxin (PDB: 1lta) (Merritt *et al*, 1994) with an RMSD of 3.05 Å over 105 equivalent positions (Fig. EV4B). According to this comparative analysis, we were unable to assign equivalent residues to the active site residues in enterotoxin. Therefore, we elected residues Glu107, Lys254, and Cys262, which based on the model, form a potential catalytic triad in an exposed pocket (Fig. 10C). However, individual substitutions of these residues were unable to alleviate the yeast toxicity phenotype, suggesting that these mutations are insufficient to alter activity of this effector (Fig. 10E).

Lpg2527/LnaB is an effector involved in the activation of the NF-κB pathway during infection of HEK293T cells and bone marrow macrophages (Losick *et al*, 2010). The molecular mechanism of this activation remains unknown. However, it has been demonstrated that residues 361 to 410 which are suggested to form a potential helical bundle are important for protein-protein interactions (Losick *et al*., 2010). The full-length model of Lpg2527/LnaB features a helical bundle (residues 1-110) that leads into a compact domain with a unique fold spanning residues 111 to 333 (Fig. EV5A). Additional structural elements in the model include a long ɑ-helix (residues 339-393), a small three-helical bundle backed against the long helix (residues 397-497), and an extended disordered region to its C-terminus. The domain with a unique fold predicted in Lpg2527/LnaB consists of two helical bundles packed against each other and interrupted by an eighty-residue insert that appears mostly unstructured except for a small β-sheet region, packed against the base of the two helical bundles (Fig. EV5A). Phylogenetic analysis shows that this potential domain is broadly distributed in bacteria and present in over 400 proteins with different architectures. In *L. pneumophila* alone, it is found in four effectors (Lpg2527/LnaB, Lpg0437/SidL, Lpg0208, and Lpg0209) (Fig. EV5B). Similar sequences are found in other pathogens, such as *Coxiella burnetii*, several species of *Pseudomonas,* and *Vibrio*, including several strains of *V. cholerae* (Supplemental Table 4). Within this predicted unique fold, we were able to distinguish several highly conserved residues arranged in a potential catalytic triad (Ser261-His305-Glu309) co-localized between the base of the helical bundle and the β-sheet (Fig. EV5C). The substitution of Ser261 to an alanine residue rescued the yeast toxicity phenotype; however, in comparison, the expression of the Lpg2527 His305Arg variant caused toxicity like the wild type (Fig. 10D-E). The Glu309Ala substitution had a dramatic reduction of protein expressed in yeast cells, thus, resulting in this variant being excluded from our experimental panel. Other conserved residues in the Lpg2527/LnaB model that may be part of this putative active site include Thr263 and Tyr304 (Fig. EV5C); however, individual substitutions of these residues to an alanine and arginine, respectively did not lead to alleviation of this effector’s toxicity.

In total, our structural model-guided approach combined with the use of the yeast model system led to the identification of specific residues in three out of four *L. pneumophila* effectors with predicted unique folds. Combined with studies of cryptic domain activity in this model system, these results yet again demonstrated the power of such a combinatorial approach and provided a roadmap to further functional characterization of novel activities across this effector arsenal.

## Discussion

Over ten percent of the *Legionella pneumophila* proteome accounts for effector proteins that are translocated by the Dot/Icm secretion system into the eukaryotic host. Dissecting the individual functions of the largest arsenal of secreted pathogenic factors represents a significant challenge that hampers our understanding of this pneumonia-causing bacterium’s infection strategy. The recent dramatic improvements in protein 3D structure prediction have provided valuable new tools for globally assessing the potential functional domain repertoire in *L. pneumophila* effectors. By combining advanced protein 3D structure modeling with a model system phenotypic assay, we present an expansive overview of predicted functional domains in over 360 *L. pneumophila* effectors. Our comprehensive analysis has unveiled a remarkably diverse repertoire of predicted folds and functions. This range of predicted functional domains not only enriches our understanding of *L. pneumophila* pathogenicity, but, more importantly, provides a foundation for the functional characterization of potential novel functional entities within previously uncharacterized effectors.

Previous large-scale studies of *L. pneumophila* effectors, based on primary sequence analysis, highlighted the presence of a significant number of structural motifs typically present in the protein of eukaryotic organisms, defined as 75% and above of protein sequences being encoded in eukaryotic genomes (Burstein *et al*., 2016; Gomez-Valero *et al*., 2019). Among such motifs, so-called tandem repeat motifs, including the ARM, ANK, and LRRs, were identified as the most recurrent in *L. pneumophila* effector proteins. Primarily recognized for their role in protein-protein interactions, these structural elements are now acknowledged as more versatile molecular recognition modules that can also be involved in protein-lipid and protein-sugar interactions (Islam *et al*, 2018). Confirming the prevalence of tandem repeats in *L. pneumophila* effectors, the analysis of their predicted 3D models suggested an even larger presence of these structural elements, particularly for ARMs. The identification of tandem repeat motifs in conjunction with other cryptic functional domains in an effector model may indicate their combined role in interactions with the appropriate host substrate, which, in many cases, awaits identification and functional characterization. Notably, all nine effector models with predicted LRRs lacked other functional determinants. This observation may suggest the unique role of LRRs as autonomous protein regulator molecules in the effector arsenal.

Another eukaryotic-like feature of *L. pneumophila* effectors that became apparent in our analysis is the overrepresentation of proteins with intrinsically disordered regions (IDRs). IDRs have been increasingly recognized as an integral part of a cellular proteome that does not fold into a specific 3D structure, but rather performs their function while maintaining a range of alternative conformations (Wright & Dyson, 1999). IDRs are estimated to be present in over 40% of a given eukaryotic proteome and their role in different cellular processes is only starting to emerge (Latysheva *et al*, 2015). In the case of analyzed *L. pneumophila* effector models, we estimated IDRs to be present in at least 24% of these predicted effector models, as compared to 4.8% estimated for the remainder of this bacterium’s proteome. While these estimates are based on sequence-based IDR predictions, they are further supported or even extended by AlphaFold2 predictions, which are accepted to have high accuracy for the identification of such regions (Zhao *et al*, 2023). Notably, 8 *L. pneumophila* effector models do not contain any recognizable structural elements, suggesting these proteins to be full-length IDRs. While this may be the result of the Alphafold2 algorithm’s limitation in predicting structural elements in these effectors, it also raises an intriguing question of IDR effector potential function during infection and possible functional mimicry with host cell IDR counterparts. Such an observation further supports and expands on the previous hypothesis about the evolutionary importance of interkingdom horizontal gene transfer for the acquisition of “eukaryotic-like *L. pneumophila* effectors (de Felipe *et al*., 2005; Gomez-Valero *et al*., 2011).

Our analysis also highlighted the prominence of transmembrane helices (TMs) in effector models, thus indicative of membrane localization to host cell organelles or the LCV. Effectors possessing such structural elements remain one of the most understudied categories of the *L. pneumophila* effector arsenal. TMs were identified in 62 of the effector models with the number of them varying from 1 to 11. Notably, in the case of 22 effectors, TM helices were the only structural element recognized in their model often accompanied by only disordered regions or helical bundles. In the remaining effectors in this category, models contained other recognizable structural elements or cryptic functional domains with potential enzymatic activity, such as cysteine protease or ɑ-β hydrolase.

Our analysis suggested the presence of numerous cryptic domains that share structural features with established protein families. This included new members of prominent protein families, such as cysteine proteases and kinases, two families which have been already suggested as having the most representatives in the *L. pneumophila* effector arsenal. Less expectedly, we also identified representatives of other protein families, such as osmotin-like domains and the RIFT-related alanine racemase family (Supplementary Table 1), that were, according to our knowledge, never associated with any effector arsenals. The specific roles of such domains in the effectors’ functions remain to be determined.

In the case of cysteine proteases, which represent the most abundant functional domain identified in *L. pneumophila* effectors, our analysis has doubled the estimated number of effectors expected to carry such a domain. These included several effectors with predicted similarity to acetyltransferase domains previously described in YopJ effectors translocated by the T3SS system. Along with other *L. pneumophila* effectors, such as Lpg2147/MavC and Lpg2148/MvcA, which contain structural domains previously identified only in the T3SS effector arsenals (Valleau *et al*., 2018), and other examples discussed below, these observations highlight the phenomenon of shared host manipulation strategies between bacterial pathogens with drastically diverse pathogenic lifestyles and effector translocation systems.

As mentioned above, many of the cryptic functional domains that we identified were accompanied in effector models by additional structural motifs - such as helical bundles, TMs, or tandem repeats - which may contribute to their cellular function as localization signals or substrate binding determinants. The presence of such additional functional elements would be particularly important for of kinase domains where predominantly *α*-helical elements outside of the main enzymatic domain were shown to be responsible for the recruitment of a specific substrate (Lee *et al*., 2020). Several effector models with cryptic functional domains also included *L. pneumophila* effector-specific substrate recruitment elements. One such example is the helical insertion domain characterized in the aforementioned Lpg2147/MavC and Lpg2148/MvcA effectors to recognize their host targets (Puvar *et al*, 2020). Our analysis suggests the presence of a similar structural element in several effector models, including that of Lpg3000, which is conserved across all sequenced *Legionella* species (Gomez-Valero *et al*., 2019).

A significant number of structural domains predicted in *L. pneumophila* effector proteins lacked similarity with experimentally characterized protein structures. In 133 of these effectors, the domains consisted of multiple α helices assembled in various α helical bundles. In 35 effectors, these helical bundles were the only recognized structural motifs. Such structural motifs appear common in a given proteome and are particularly challenging for functional annotation due to the lack of obvious functionally related structural signatures. However, two recently characterized *L. pneumophila* effectors provide an indication of how these structural motifs can be adapted to a specific function in the host cell. The most striking example is Lpg2829/SidH, which represents the largest effector in the arsenal and the second-largest protein encoded by *L. pneumophila*. Recently, the structure of Lpg2829/SidH was determined using cryogenic electron microscopy, which revealed the effector to be exclusively α-helical (Sharma *et al*, 2023). The arrangement of the helical bundles in Lpg2829/SidH does not share significant structural similarity with any proteins in the PDB (Sharma *et al*., 2023). Its atypical structural architecture enables Lpg2829/SidH to bind with human t-RNA, which was shown to be associated with the effector’s toxic phenotype when overexpressed in HEK293T cells (Sharma *et al*., 2023). Another *L. pneumophila* effector, Lpg2327/Lug15 - which is predicted to have an exclusively α-helical structure - has been demonstrated to possess E3 ubiquitin ligase activity (Ma *et al*, 2023). Given the recurrent prediction of unique α-helical structures in *L. pneumophila* effectors, we anticipate more specific activities to be associated with such structural arrangements.

A subset of *L. pneumophila* effector models contained domains with unique folds. Notably, some of these proteins shared significant sequence similarity with effectors encoded by other bacterial pathogens suggesting shared functionality. Using the yeast model system, where the ectopic expression of an individual effector often results in significant growth defect phenotype, we were able to test the role of conserved, co-localized, and surface-exposed residues of these domains with predicted unique folds. In three out of four tested effectors with predicted domains of unique folds, this approach identified specific amino acids critical for these proteins’ activity in the yeast system. Using the same model cell assay approach, we were also able to confirm the role of active site residues in ten predicted functional domains as causing yeast growth defects. Along with these new members of the established protein families, the effectors with predicted unique folds represent an exciting basis for the discovery of new eukaryotic cell manipulation mechanisms that are evolved by *L. pneumophila* and potentially shared by other bacterial pathogens. The additional 27 domains with predicted novel folds that are not lethal in the yeast model system provide an interesting set for further exploration.

To conclude, by combining advanced 3D structural predictions of all reported *L. pneumophila* effectors with functional assays, we have converged on a full catalog of predicted functions and mechanisms of how this pathogen takes control and thrives inside eukaryotic phagocytic cells. With the ability to analyze both sequence and 3D structural models, we were able to precisely pinpoint functional residues that are essential for several cryptic domains by experimentally confirming their importance for the activity of the *L. pneumophila* effector in yeast cells. Many more still await a full experimental verification. By putting together all the annotations and analyses in the interactive web-based database, we provide a centralized starting point for the functional studies of specific *L. pneumophila* effectors with a clear map of predicted and experimentally characterized functionally relevant 3D elements.

## Methods

### 3D modeling of *L. pneumophila* effectors and assignment of domain types based on ECOD hierarchy

Primary sequences and 3D structural models of all 368 *L. pneumophila* effector proteins were analyzed to assign their domain architectures and, when possible, predict potential biochemical functions.

The 3D models calculated with AlphaFold v2.0 (Jumper *et al*., 2021) were retrieved from the AlphaFold DB database (Varadi *et al*, 2022), where available or built locally. For a small number of the longest effectors, which exceeded AlphaFold length limits, sequences were split into overlapping fragments that were modeled individually.

The unstructured regions of the models were removed and the structured regions were divided into compact fragments, corresponding to potential structural domains. These putative domains were then compared to structures of domains of experimentally characterized proteins retrieved from the ECOD database version 288 (Cheng *et al*., 2014) and reduced by clustering by sequence identity with a 40% cut-off. The structural comparison was performed using the FATCAT (Ye & Godzik, 2003) program without allowing for twists in the aligned structures. The results of FATCAT searches were analyzed, and the putative domains were assigned to domain types using the ECOD topology (T) level when possible (132 domain types are assigned at this level). In cases when structural similarities did not allow for clear assignment at the topology level, the assignment was done at the higher ECOD homology (H) level (19 domain types are assigned at this level). In some cases, strong sequence and/or structural similarity made it possible to assign effector domains to ECOD at the family level (F) (6 of the assigned domain types correspond to ECOD families). In cases when, according to the FATCAT results, the fragments of effector models did not correspond to complete ECOD domains, the initial division of models into potential domains was revised, and FATCAT searches were repeated.

Independently, the primary sequences of the effectors were compared to primary sequences of entries from the ECOD and PDB databases using Blastp (Altschul *et al*., 1997) and HHPred (Gabler *et al*, 2020) algorithms. The primary sequence similarities, if significant, were used to verify structure-based assignments of regions to the ECOD domain topologies and to resolve some cases where assignments to ECOD topologies were difficult to make based on structural similarity. The data about the presence and locations of signal peptides and transmembrane helices were downloaded from the Uniprot database (UniProt, 2023). Regions with at least two consecutive transmembrane helices were labeled as “transmembrane regions” and included in the annotations of domain architectures unless it was possible to assign them to specific ECOD domain types. Long stretches of structural disorder regions were indicated by the lack of structured domains as predicted with AlphaFold, were classified as “disordered regions” and also included in the descriptions of domain architectures.

In over 120 effector proteins, we found regions modeled by AlphaFold as a series of helical hairpins or bundles without unambiguous matches to structurally characterized proteins. If their primary sequences also did not show any similarity to experimentally characterized structures, we labeled these fragments as “helical regions”.

### Identification of eukaryotic-like domains in *L. pneumophila* effectors

Microbial protein families and domains with mostly eukaryotic homologs are regarded as eukaryotic-like domains. They are more likely to be involved in interference with the signaling and metabolism of the host’s cell. Here, the eukaryotic-like domains were identified among effector domains as follows:

1. The sequences of ECOD representatives of a given domain (clustered by sequence identity with a 40% cut-off) were used to start Blastp searches against the set of 22925 representative proteomes downloaded from the UniProt database.
2. The percentage of significant (e-values < 0.001) unique Blastp hits (presumed homologs) which came from eukaryotic organisms was calculated for each domain type.
3. Domain types with more than 75% of eukaryotic homologs were labeled as eukaryotic-like.

According to this criterion, we labeled 29 out of the 153 identified ECOD domain types as eukaryotic-like (Table 1).

### Identification of *L. pneumophila* effector domains present in effectors from other species

The effector domains with homologs in other species were labeled as follows:

1. The sequences of all ECOD representatives of a given domain type (clustered by sequence identity with a 40% cut-off) were used to start Blastp searches against the SecretEPDB database (An *et al*, 2017) (the *L. pneumophila* effectors themselves were excluded from the set of SecretEPDB sequences).
2. The effector domains with significant (e-values < 0.001) Blastp hits in the SecretEPDB database were labeled as being present in effectors from species other than *L. pneumophila*.

With the above procedure, we labeled 46 out of the 153 identified domain types as present in known effectors from other organisms.

### Cloning and DNA manipulations

The pDONR221-effector constructs used in this study were obtained from the pDONR221-Dot/Icm substrate library from a previous report (Losick *et al*., 2010; Urbanus *et al*., 2016). Point mutations of each WT effector in pDONR221 were performed using Quik change, as previously described (Liu & Naismith, 2008; Popov *et al*, 2023). Primers used for the generation of point mutations can be found in Supplemental Tables 5 and 6. The plasmid DNA of each construct was obtained using a PRESTO mini prep plasmid extraction kit (Geneaid, Taiwan), and then validated by sequencing. All pDONR221 constructs containing the WT effectors and their variants were then cloned into the pGAL423-FlAG, or pGAL416-HA (in the cases of Lpg2461 and Lpg2523/Lem26), *S. cerevisiae* overexpression construct using the Gateway LR clonase kit (Cat#11791100, Thermo Fischer).

### Yeast Spotting Assays

All the overexpression constructs were transformed into the *S. cerevisiae* BY4741 strain using a previously described lithium acetate procedure (Salomon & Sessa, 2010). The spotting assays of all toxic effectors and their respective variants in this study were performed as previously described with minor modifications (Salomon & Sessa, 2010). In brief, cultures of toxic effectors and their mutants were grown in an SD selective medium lacking uracil and supplemented with 2% dextrose overnight. Following overnight growth, all the cultures were normalized to an OD_600_ of 1 to make a fourfold dilution series. Each effector and its mutants were spotted on a selective SD media plate lacking uracil containing either 2% dextrose (non-inducing) or 2% galactose (inducing). After spotting, plates were incubated for three days at 30°C and then imaged.

### Western Blot Analysis to Validate Effector Expression in Yeast

The expression and solubility of the toxic effectors and their mutants were performed using western blot analysis with a previously described methodology where only a few modifications were made (Salomon & Sessa, 2010; Urbanus *et al*., 2016). Overnight cultures (3 ml) in selective SD medium lacking uracil and supplemented with 2% dextrose were spun down and washed three times with ddH2O water. Cultures were then diluted to an OD_600_ of 1 in selective SD liquid media lacking uracil and supplemented with either 2% dextrose or 2% galactose, and then grown overnight. On the following day, the overnight cultures were spun down and the yeast cell pellets were resuspended in 100 µl of ice-cold lysis buffer (4% v/v 5M NaOH and 0.5% v/v β-mercaptoethanol). Cells were mixed and then incubated on ice for 40 minutes. After incubation, lysed cells were supplemented with 1 µl of 6N HCl and 50 µl of 3x sample loading buffer (0.05% [v/v] bromophenol blue, 30% [v/v] glycerol, 37.5% [v/v] 500 mM Tris-HCl pH 6.8, 0.15% [w/v] sodium dodecyl sulfate [SDS], and 500 mM Dithiothreitol) and then vortex mixed. Samples were then boiled on a heat block at 95°C for five minutes and spun down for one minute at 12000 *xg*. 20 µl of an effector construct grown in either dextrose or galactose was loaded onto a 12% SDS-PAGE gel for immunoblot analysis. Proteins were transferred onto a nitrocellulose membrane and then blocked with 5% non-fat milk at room temperature for two hours. After blocking, the membrane was then washed three times for 15 minutes with tris-buffered saline-0.1% Tween 20 detergent (TBST) buffer, and then incubated overnight with a primary antibody that was specific for the N-terminal fusion tag of each over-expression construct (α-FLAG [Cat#8146S, Cell Signalling Technology] or α-HA [Cat#3724S, Cell Signalling Technology]) at a dilution of 1:1000. Following primary antibody incubation, the nitrocellulose membrane was washed with TBST three times for 15 minutes. After the washing, the membranes were incubated with the appropriate horse radish peroxidase-conjugated secondary antibody for 1 hour at room temperature. The signal was detected with an Immobilon Western chemiluminescent HRP substrate (Cat#WBKLS0500, Millipore Sigma, Canada). Immunoblots were visualized using a BioRad ChemiDoc machine. The loading control western blots were performed with an α-GADPH antibody (Cat#97166S, Cell Signalling Technology) using the previously mentioned procedure.

## Acknowledgments

We would like to thank Dr. Nobuhiko Watanabe and Dr. Elina Karimullina from the Savchenko Laboratory for their insightful discussions. The work from this manuscript was funded in whole or in part with U.S. Federal funds from the National Institute of Allergy and Infectious Diseases, National Institutes of Health Department of Health and Human Services, which is under Contract numbers: HHSN272201700060C and 75N93022C00035. These funds were awarded to the Center for Structural Biology of Infectious Diseases (CSBID, http://csbid.org). The functional data in this manuscript was supported by a Canadian Institutes of Health Research Project Grant (A.W.E, A.S) (PJT-162256) and a National Science and Engineering Research Council of Canada (NSERC) Discovery program grant that was awarded to the principal investigator, A.S. Computations were performed using the computer clusters and data storage resources of the UCR High Performance Computer Cluster (HPCC), which were funded by grants from NSF (MRI-2215705, MRI-1429826) and NIH (1S10OD016290-01A1) D.T.P was supported by the Doctoral Alberta Graduate Excellence Scholarship.

## Data Availability Section

Analysis of all effector models can be viewed in the following database: https://pathogens3d.org/legionella-pneumophila.

## Expanded View Figure Legends

**Figure EV1.**
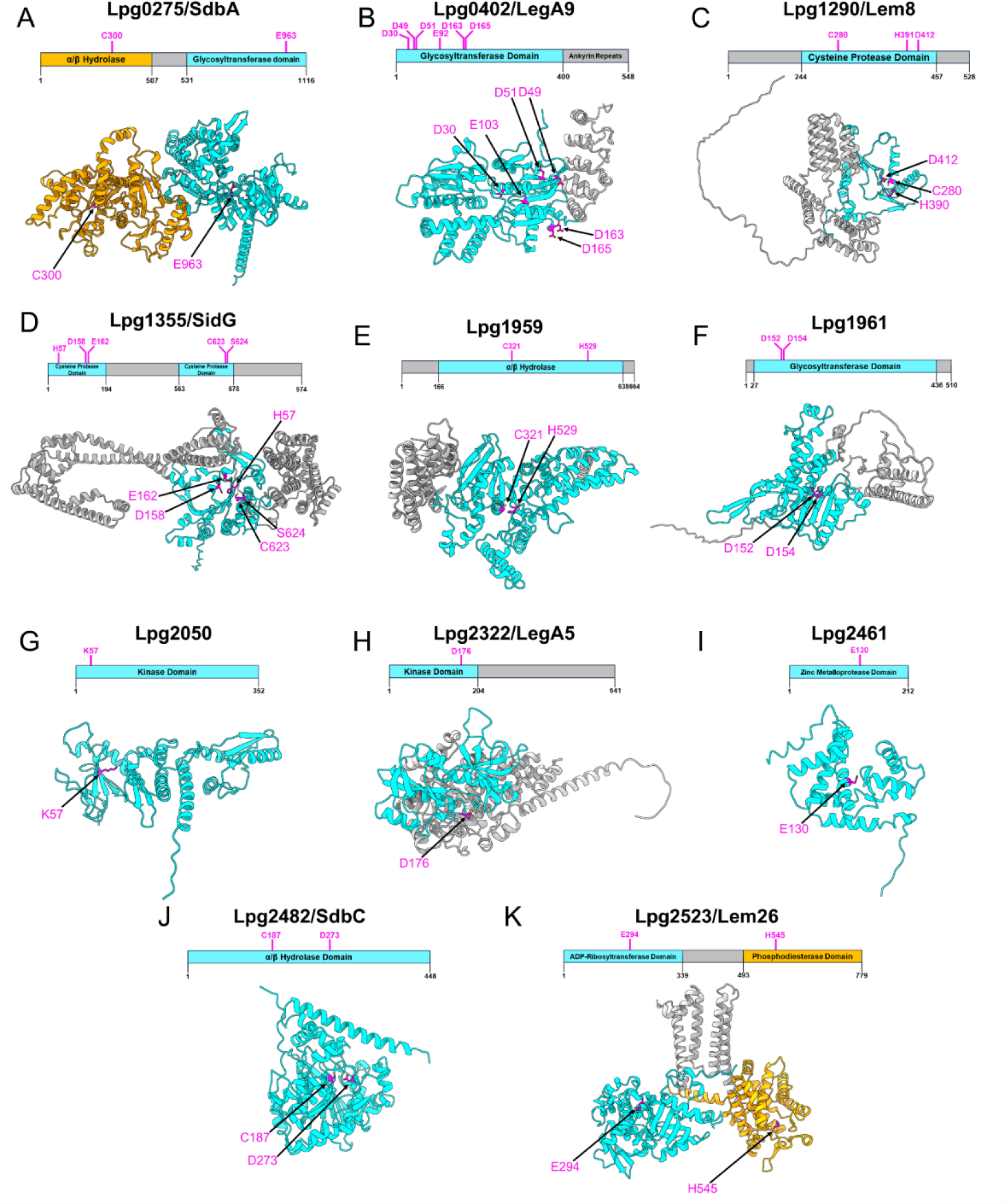
Alphafold2 models of *L. pneumophila* effectors of uncharacterized effectors with cryptic domains. A-K. Molecular architecture of effectors harboring cryptic domain, which displays residues (magenta) that were tested in Figure 9. Cryptic domains were highlighted in cyan while other structural elements in the models were highlighted in grey. The secondary cryptic domains in Lpg0275/SdbA and Lpg2523/Lem26 that led to very little to no alleviation of toxicity in Figure 9 were highlighted in gold.

**Figure EV2.**
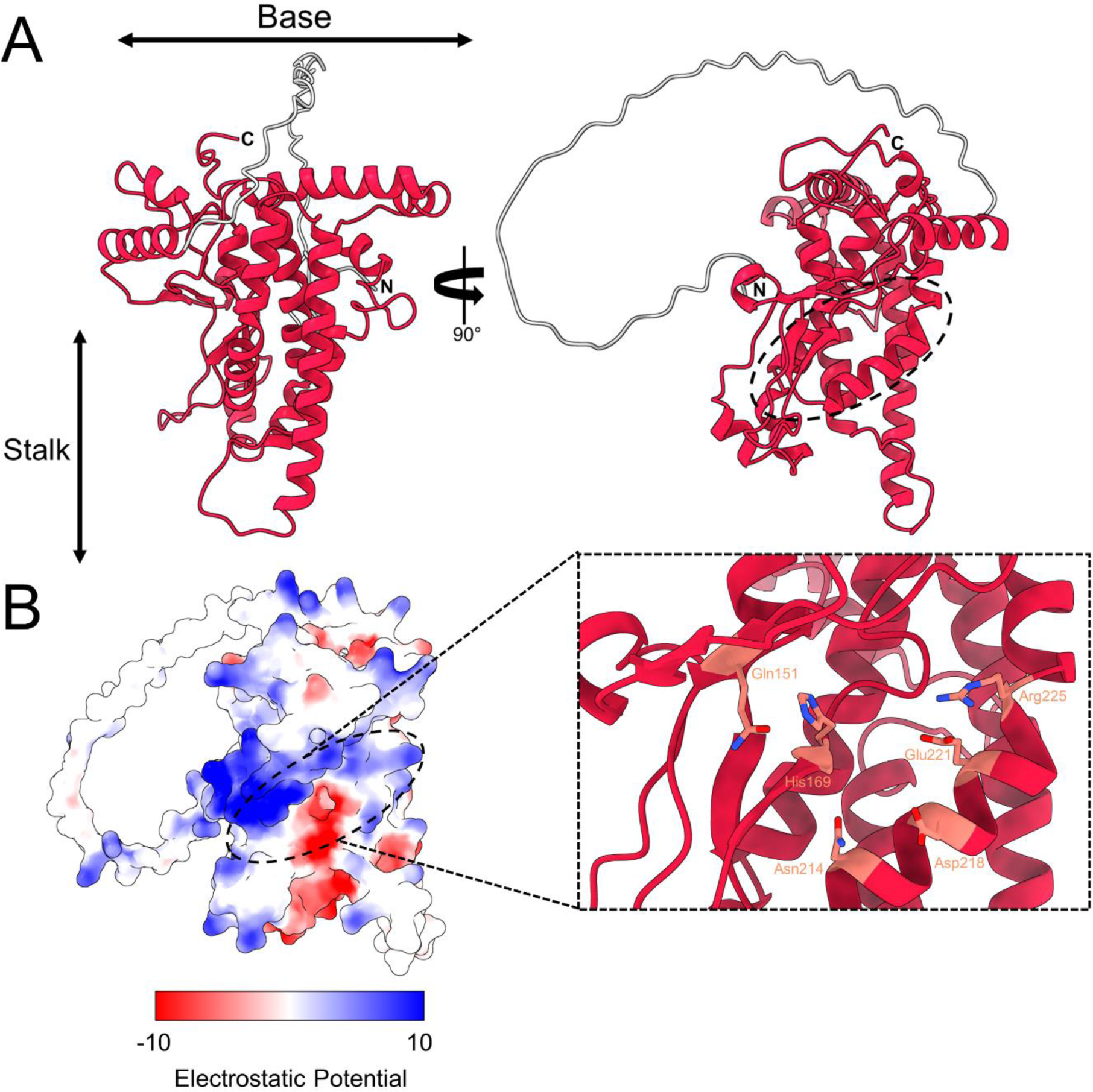
Lpg1154/RavQ forms a unique “T” shape containing a highly conserved groove that may serve as an active site. A. The Lpg1154/RavQ model (residues 59-389, red) is followed by a view of the potential active site cavity of Lpg1154/RavQ with an electrostatic potential surface representation of that site. B. Conserved residues identified in Lpg1154/RavQ orthologs from the *Legionella* genus that are arranged in a potential active.

**Figure EV3.**
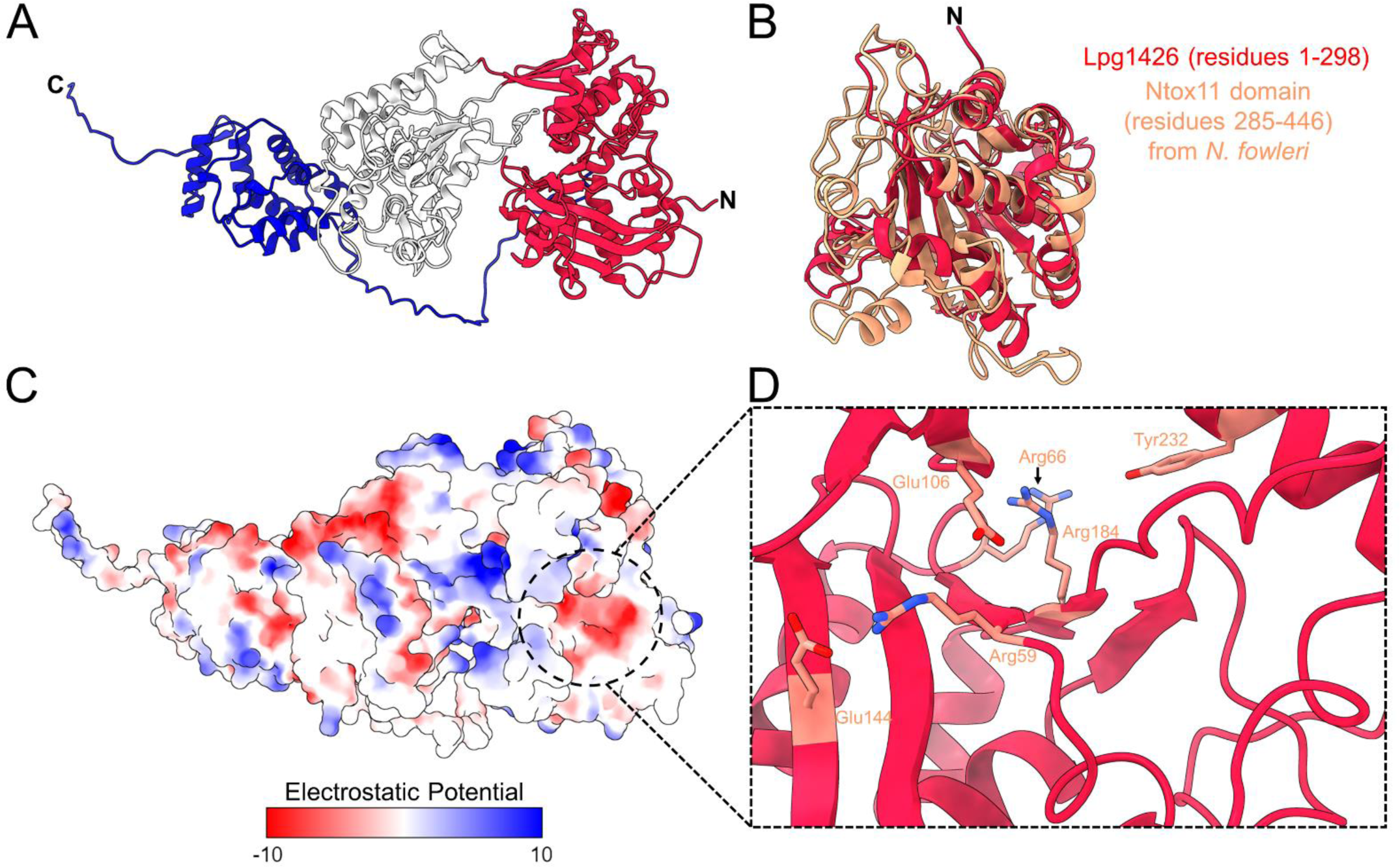
Lpg1426/VpdC has a unique domain on the N-terminus that has structural similarity to the Ntox11 putative toxin found in human pathogenic amoeba. A. Cartoon representation of the Lpg1426/VpdC Alphafold2 model. The unique fold is found on the N-terminus (red), followed by a central phospholipase domain (white) and the C-terminal helical bundle involved in interactions with ubiquitin. B. Structural alignment of the Lpg1426/VpdC unique fold (residues 1-298, red) onto the Ntox11 Alphafold2 model (residues 285-446, salmon) from *N. fowleri*. C. Surface representation of the electrostatic potential of the Lpg1426/VpdC model that also shows a conserved negatively charged pocket (dotted circle) is present in the unique fold. D. Zoom in on the positively charged region where highly conserved residues (salmon sticks), which are present in the *Legionella* orthologs of Lpg1426/VpdC, form this pocket.

**Figure EV4.**
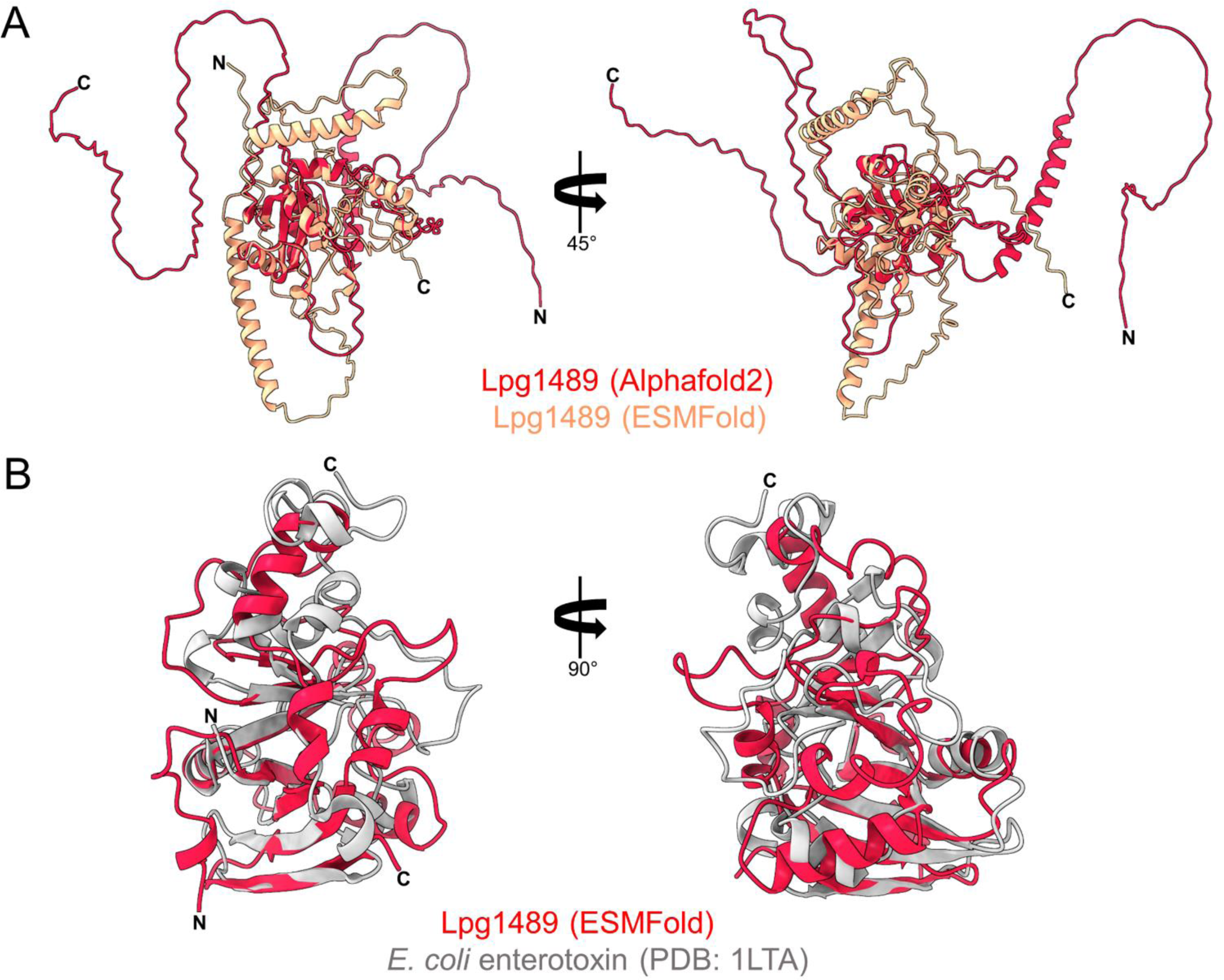
Lpg1489/RavX has a central unique fold surrounded by disordered loops. A. Structural alignment of the full-length Lpg1489/RavX models generated by Alphafold2 (red) and ESMFold (salmon). B. ESMFold model of the Lpg1489/RavX unique fold (residues 83-263, red) onto the *E. coli* enterotoxin (PDB: 1LTA, residues 1-181, grey).

**Figure EV5.**
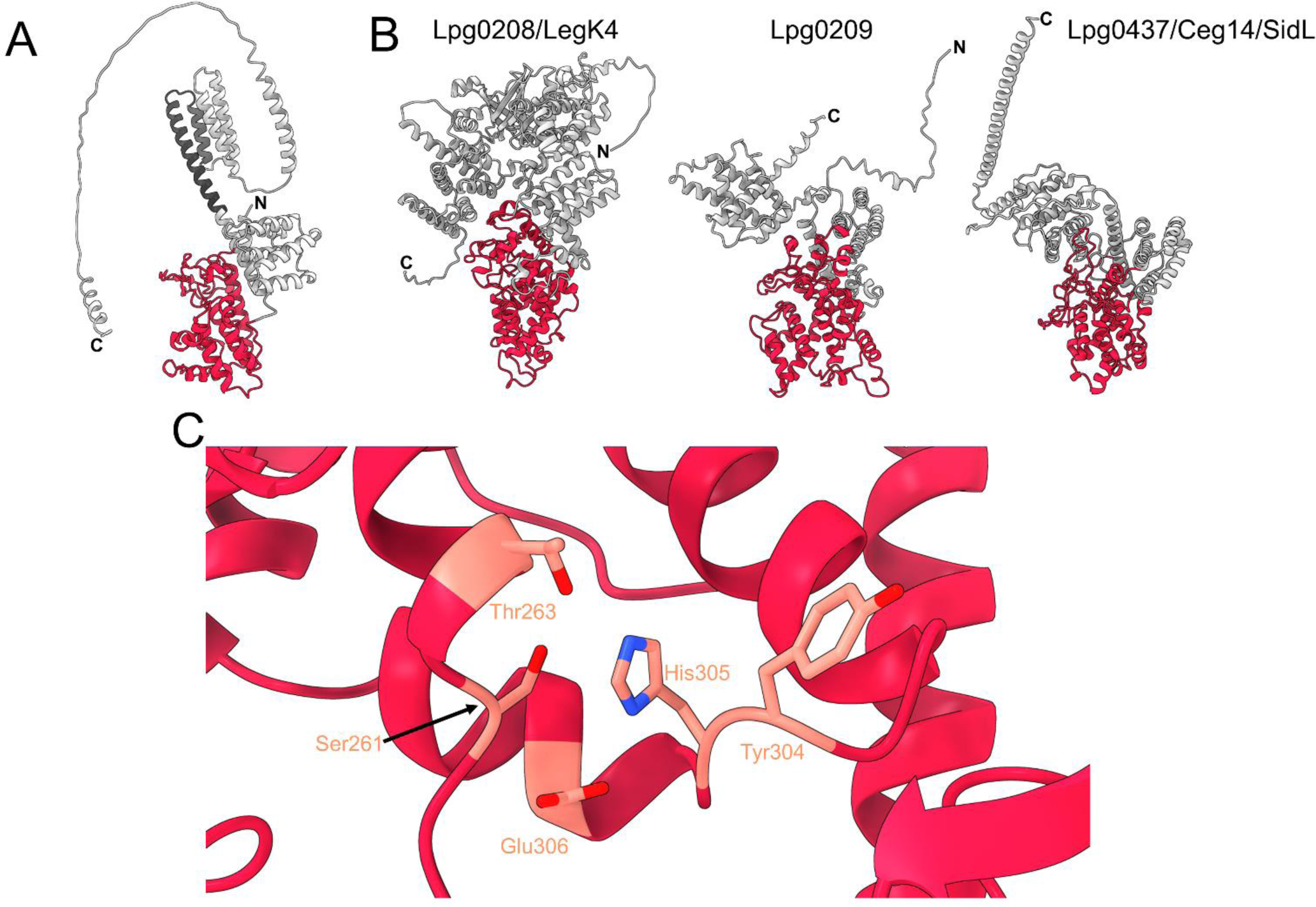
The unique domain in Lpg2527/LnaB is present in other *L. pneumophila* effectors, harboring a conserved set of residues that resemble a potential active site. A. Alphafold2 model of Lpg2527/LnaB which highlights the unique fold (red) and the helical bundle previously shown to be important in the activation of the NF-κB pathway (dark grey) (Losick *et al*., 2010). B. Representation of the Lpg2527/LnaB unique domain (shown in red) that is present in other *L. pneumophila* effectors (Lpg0208/LegK4, Lpg0209, and Lpg0437/Ceg14/SidL). C. Zoom-in image of the putative active site of Lpg2527/LnaB that is also conserved in its *Legionella* orthologs and *L. pneumophila* effectors containing this domain.

## Supplemental Information - Figure Legends

**Appendix Figure S1.**
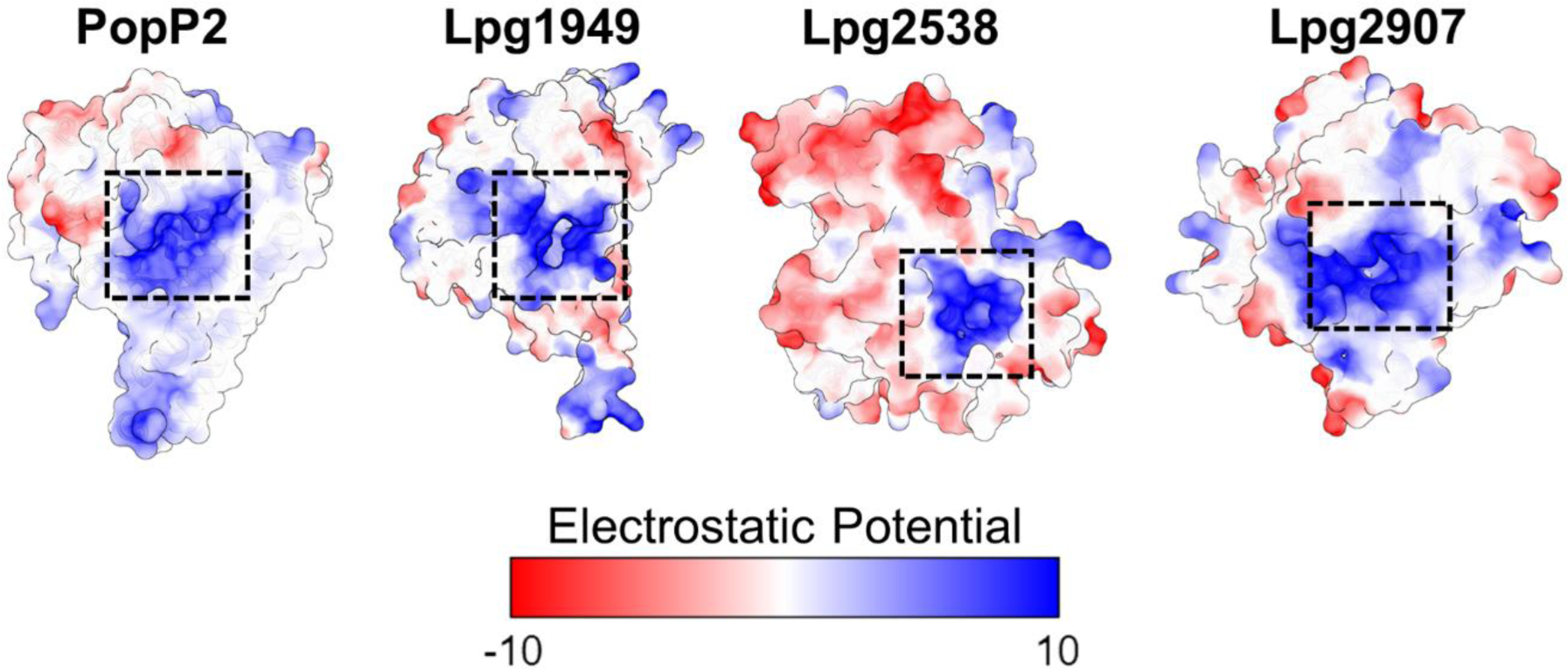
Electrostatic potential of the characterized IP6 binding site (boxed) in the PopP2 crystal structure. The corresponding IP6 sites were boxed on the surface representation of the following effectors: Lpg1949, Lpg2538, and Lpg2907. The potential IP6 binding sites were determined by structurally aligning IP6-bound PopP2 with each effector model.

**Appendix Figure S2.**
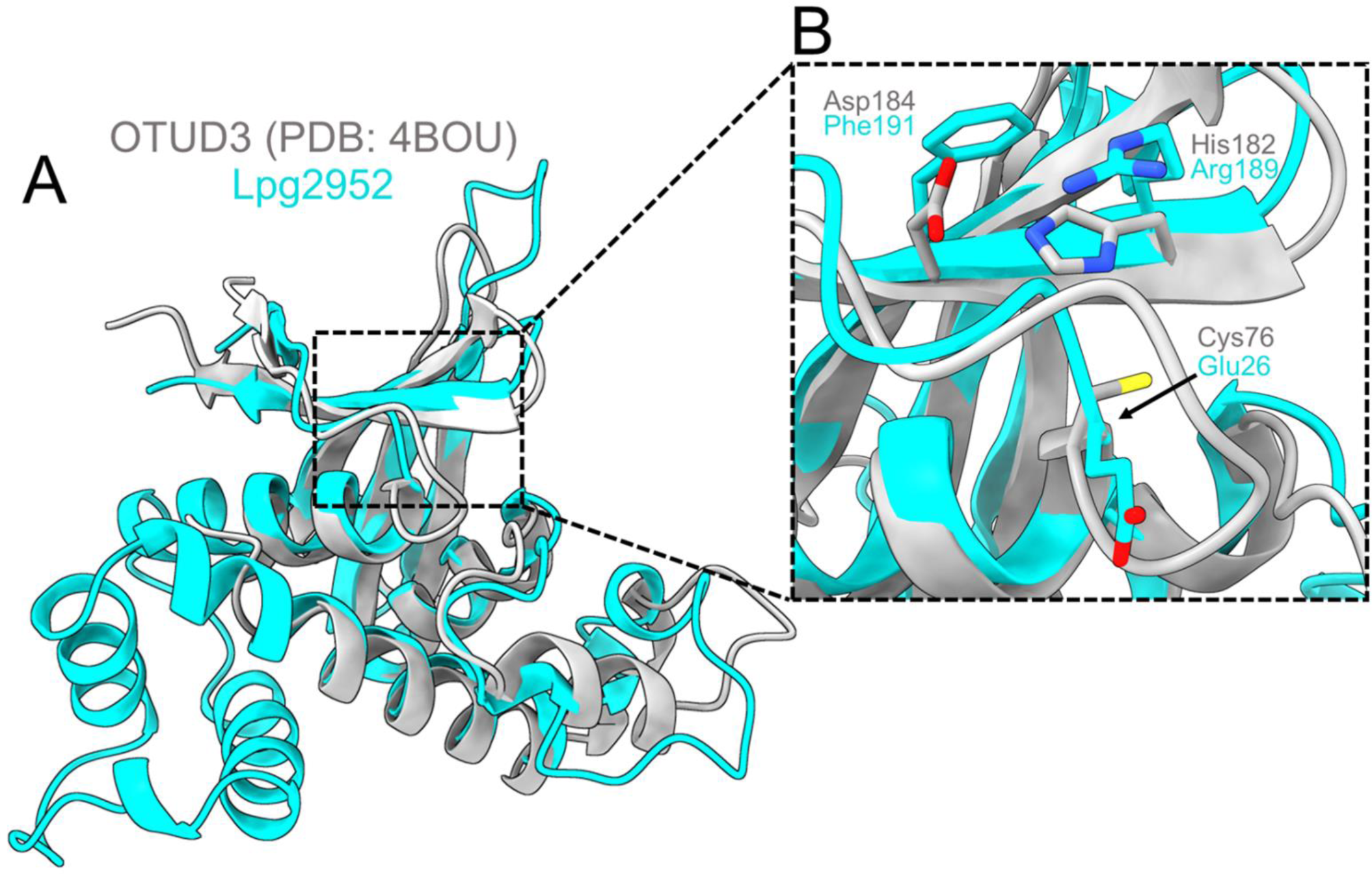
A. Alignment of the Alphafold model of Lpg2952 onto the molecular structure of the human OTU3 enzyme. B. Zoom in on the catalytic triad of OTU3 (sticks in grey) and the corresponding residues of the Lpg2952 model (sticks in cyan).

**Appendix Figure S3.**
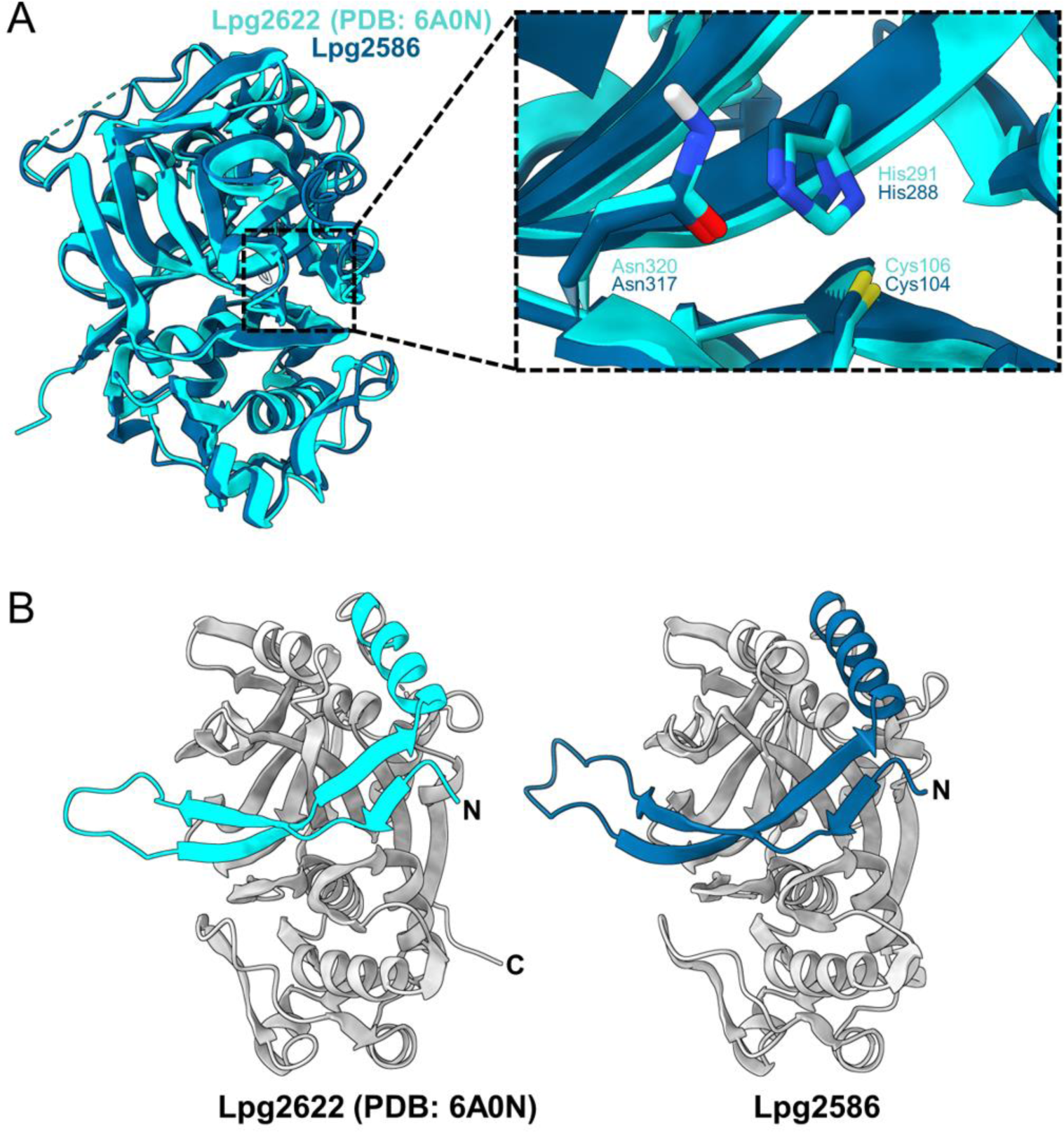
A. Structural alignment of the Lpg2592 model (navy) onto the crystal structure of T2SS *L. pneumophila* effector Lpg2622 (cyan). Inset view of the catalytic residues of Lpg2622 (sticks in cyan) and the corresponding residues of the Lpg2586 model (sticks in navy blue). B. Colored representation of the novel hairpin-turn-helix motif in the Lpg2622 crystal structure (navy) and Lpg2586 model (cyan).

**Appendix Figure S4.**
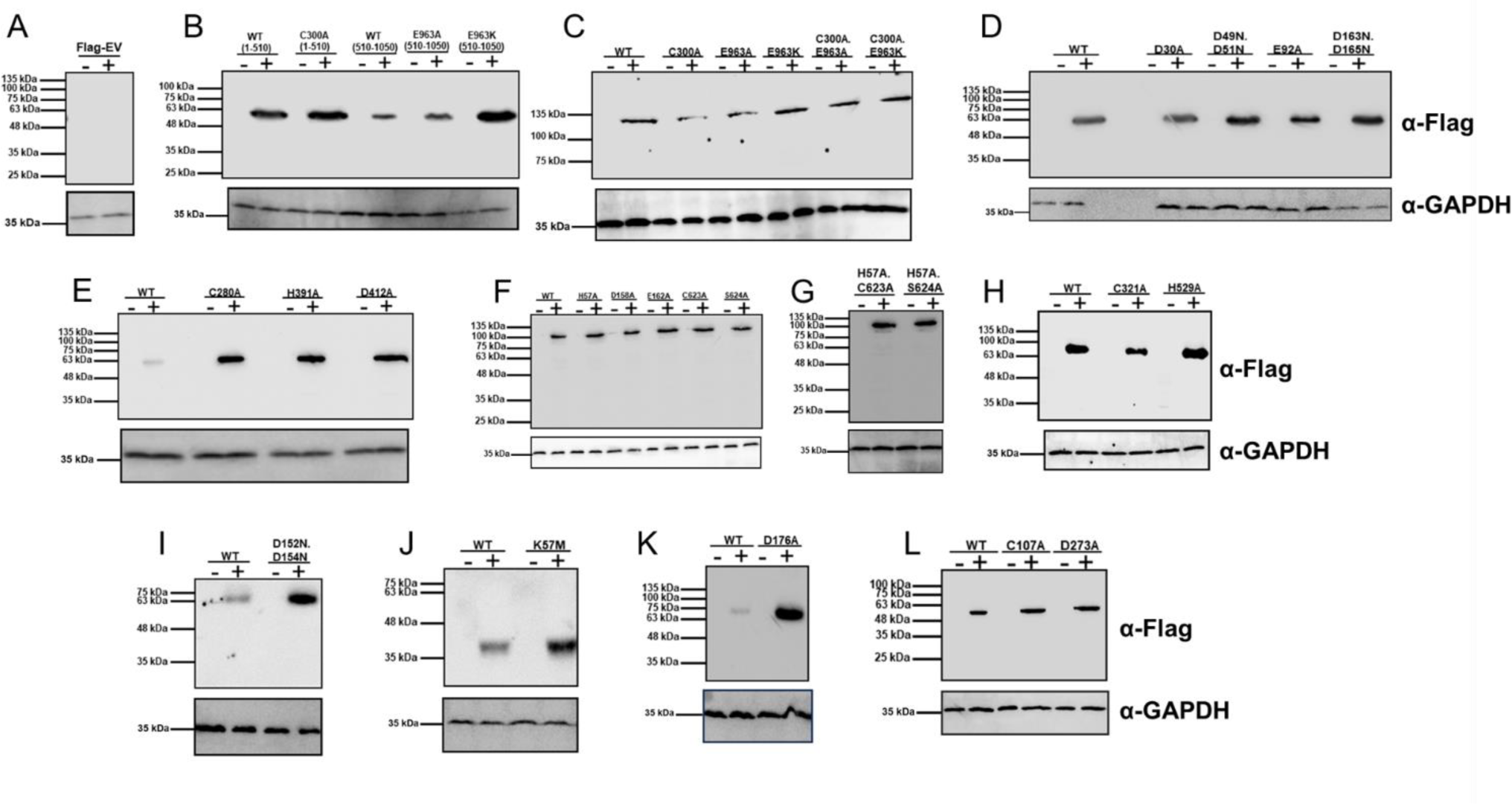
Analysis of the expression levels of (A) Flag-EV, (B) Lpg0275/SdbA fragments, and (C) Lpg0275/SdbA Full length, (D) Lpg0402/LegA9, (E) Lpg1290/Lem8, (F and G) Lpg1355/SidG, (H) Lpg1959, (I) Lpg1961, (J) Lpg2050, (K) Lpg2322/LegA5, and Lpg2482/SdbC and their mutants in the *S. cerevisiae* BY4741 strain. “-” indicates yeast incubated with dextrose SD media (non-inducing). “+” indicates yeast incubated with galactose SD media (inducing). Flag-tagged effector proteins were detected by immunoblotting with a FLAG-tag specific antibody. *S. cerevisiae* glyceraldehyde 3-phosphate dehydrogenase (GAPDH) was detected using a GAPDH-specific antibody for loading control westerns. (A) and (B) were processed on the same blot. (F) and (G) were processed on separate blots.

**Appendix Figure S5.**
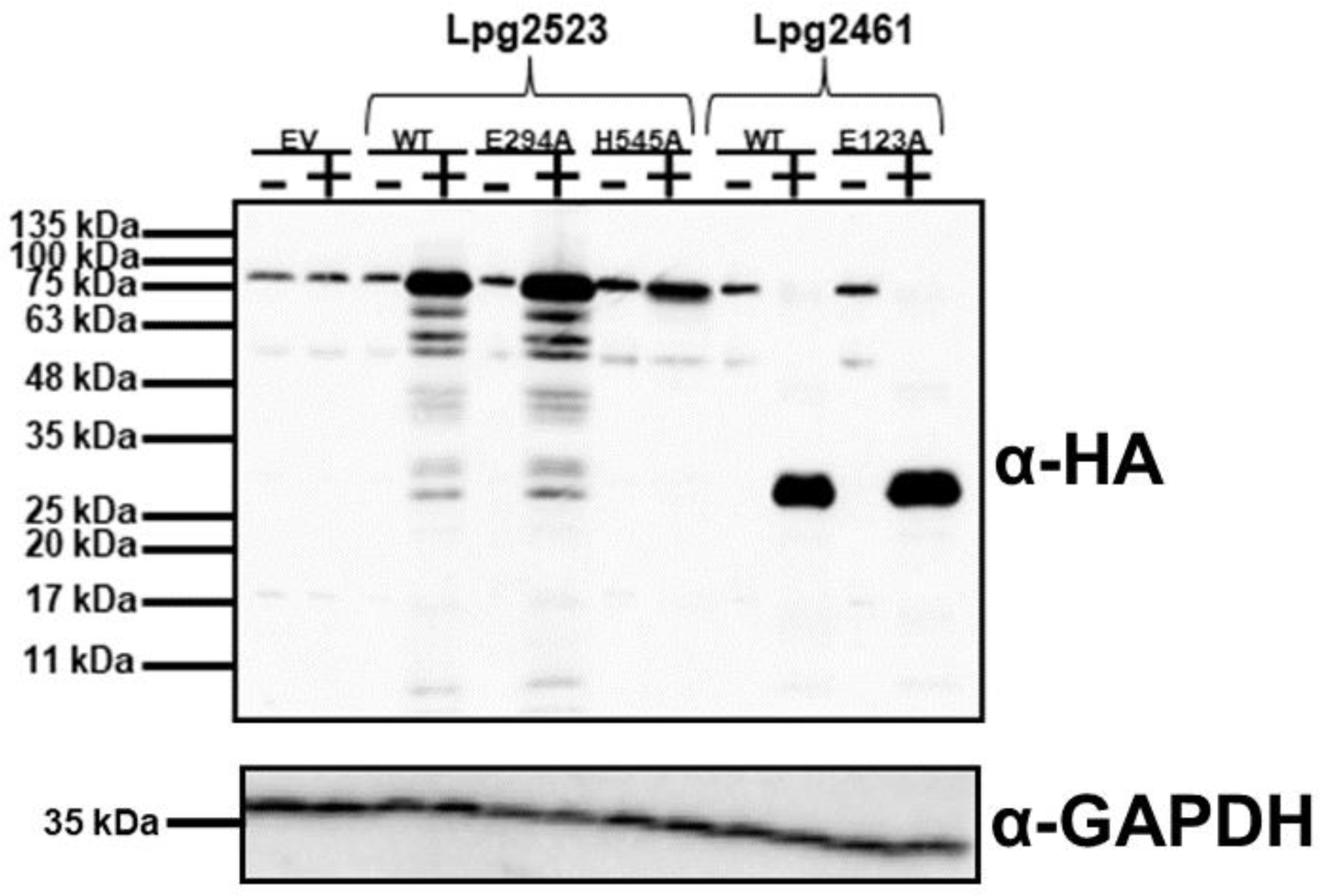
Assessment of the expression levels of Lpg2523/Lem26 and Lpg2461 and their respective mutants in the *S. cerevisiae* BY4741 strain. Yeast cells were grown in either non-inducing SD media (dextrose, “-”) or inducing SD media (galactose, “+”). An HA-tag specific antibody was used for the detection of the overexpressed HA-tagged effector protein in each sample. A GADPH-specific antibody was used in loading control westerns.

**Appendix Figure S6.**
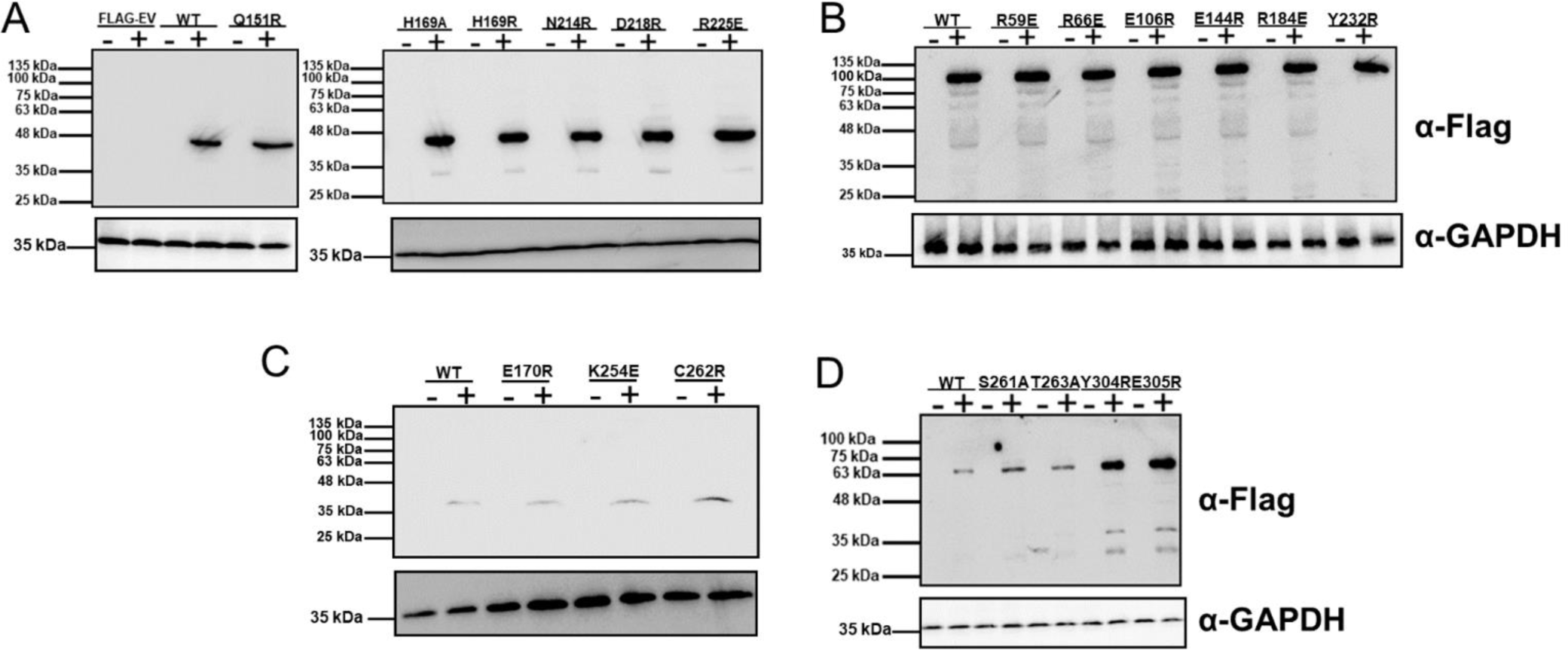
Samples of unique fold effectors and their respective mutants that were transformed in the *S. cerevisiae* BY4741 strain and were grown in dextrose-supplemented SD media (repressing, “-”) or galactose-supplemented SD media (inducing, “+”) were analyzed by Western blot using a FLAG-tag specific antibody. The loading control westerns were performed using a GADPH-specific antibody. (A) FLAG-EV and Lpg1154/RavQ, (B) Lpg1426/VpdC, (C) Lpg1489/RavX, and (D) Lpg2527/LnaB. FLAG-EV and Lpg1154 mutants were processed on separate western blots.

## Supplemental Information -Tables

**Supplementary Table 1.** The list of 368 effectors analyzed in this study with domain architectures identified in their 3D models. The legend for domain labels used in domain architectures is provided in Table 1.

**Supplementary Table 2.**
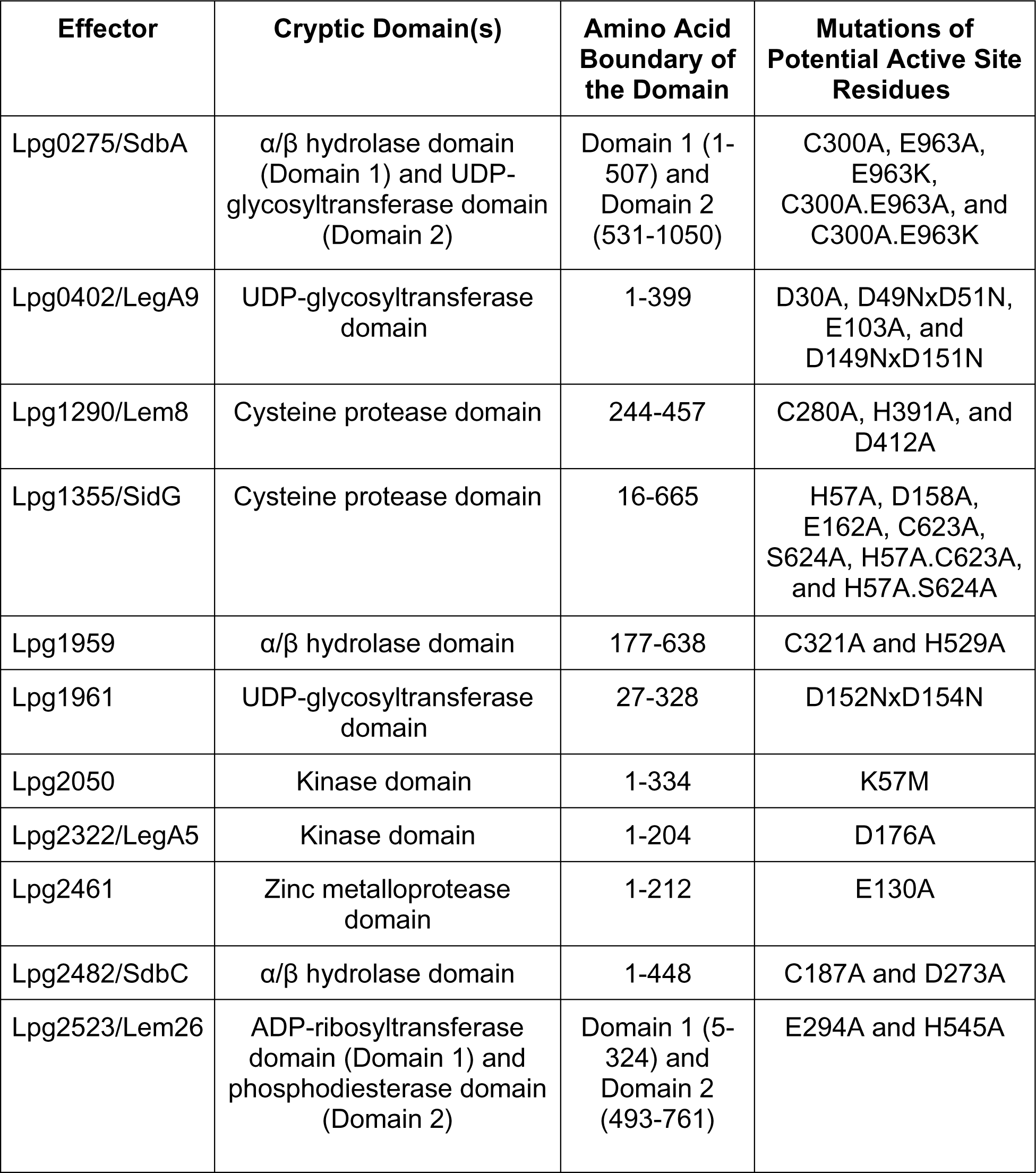
Cryptic domains of structurally uncharacterized *L. pneumophila* effectors which are known to cause a severe yeast growth defect phenotype when ectopically expressed. Point mutations targeting the putative catalytic residues are shown in Figure EV1 and tested in Figure 9.

**Supplementary Table 3.**
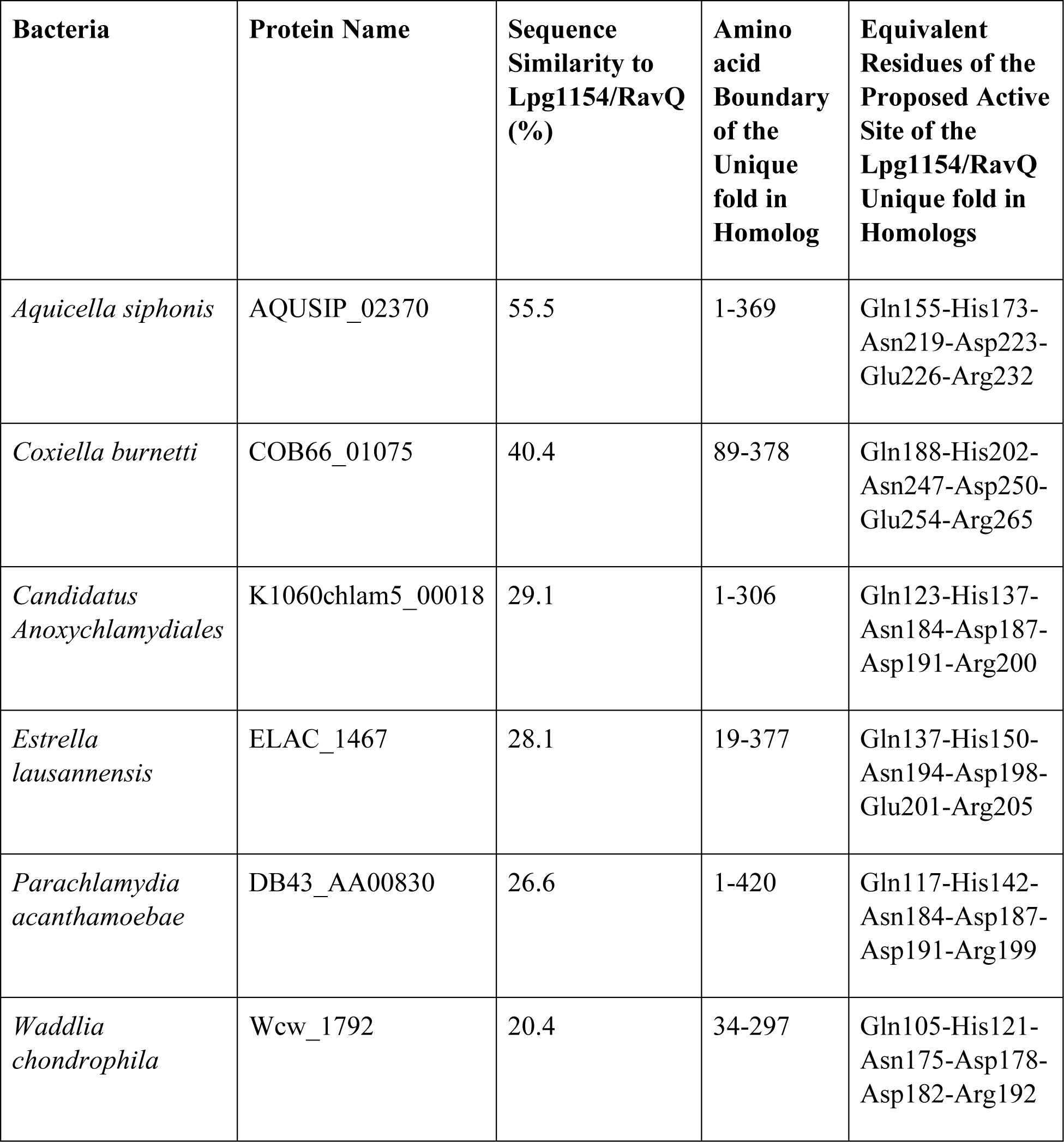
Homologs of Lpg1154/RavQ that are present in other bacterial species.

| **Bacteria** | **Protein Name** | **Sequence Similarity to Lpg1154/RavQ (%)** | **Amino acid Boundary of the Unique fold in Homolog** | **Equivalent Residues of the Proposed Active Site of the Lpg1154/RavQ Unique fold in Homologs** |
| --- | --- | --- | --- | --- |
| *Aquicella siphonis* | AQUSIP_02370 | 55.5 | 1-369 | Gln155-His173-Asn219-Asp223-Glu226-Arg232 |
| *Coxiella burnetti* | COB66_01075 | 40.4 | 89-378 | Gln188-His202-Asn247-Asp250-Glu254-Arg265 |
| *Candidatus Anoxychlamydiales* | K1060chlam5_00018 | 29.1 | 1-306 | Gln123-His137-Asn184-Asp187-Asp191-Arg200 |
| *Estrella lausannensis* | ELAC_1467 | 28.1 | 19-377 | Gln137-His150-Asn194-Asp198-Glu201-Arg205 |
| *Parachlamydia acanthamoebae* | DB43_AA00830 | 26.6 | 1-420 | Gln117-His142-Asn184-Asp187-Asp191-Arg199 |
| *Waddlia chondrophila* | Wcw_1792 | 20.4 | 34-297 | Gln105-His121-Asn175-Asp178-Asp182-Arg192 |

**Supplementary Table 4.** Homologs of Lpg2527/LnaB present in other intracellular bacterial pathogens.

| **Bacteria** | **Protein Name** | **Sequence Similarity to Lpg2527/LnaB (%)** | **Amino acid Boundary of the Unique fold in Homolog** | **Equivalent Residues of the Proposed Active Site of the Lpg2527 Unique fold in Homologs** |
| --- | --- | --- | --- | --- |
| *Aquicella siphonis* | AQUSIP_08150 | 21.3 | 266-623 | Ser513-His557Glu561 |
| *Candidatus Berkiella* | CC99x_00179 | 24.2 | 1-368 | Ser254-His298-Glu302 |
| *Pseudomonas chlororaphis* | C4K37_3919 | 26.7 | 1-434 | Ser374-His416-Glu420 |
| *Pseudomonas antarctica* | A7J50_5968 | 26.8 | 96-328 | Ser386-His430-Glu434 |
| *Vibrio caribbeanicus* | VIBC2010_01453 | 21.0 | 33-487 | Ser542-His563-Glu567 |
| *Vibrio cholerae* strain A325 | ERS013201_03035 | 22.1 | 136-490 | Ser203-His242-Glu246 |
| *Vibrio cholerae* strain IDH06781 | EYB64_18120 | 23.3 | 119-490 | Ser203-His353-Glu357 |
| *Vibrio parahaemolyticus* | MAVP-RPIeRC_00038 | 22.0 | 119-471 | Ser315-His354-Glu358 |

**Supplementary Table 5.**
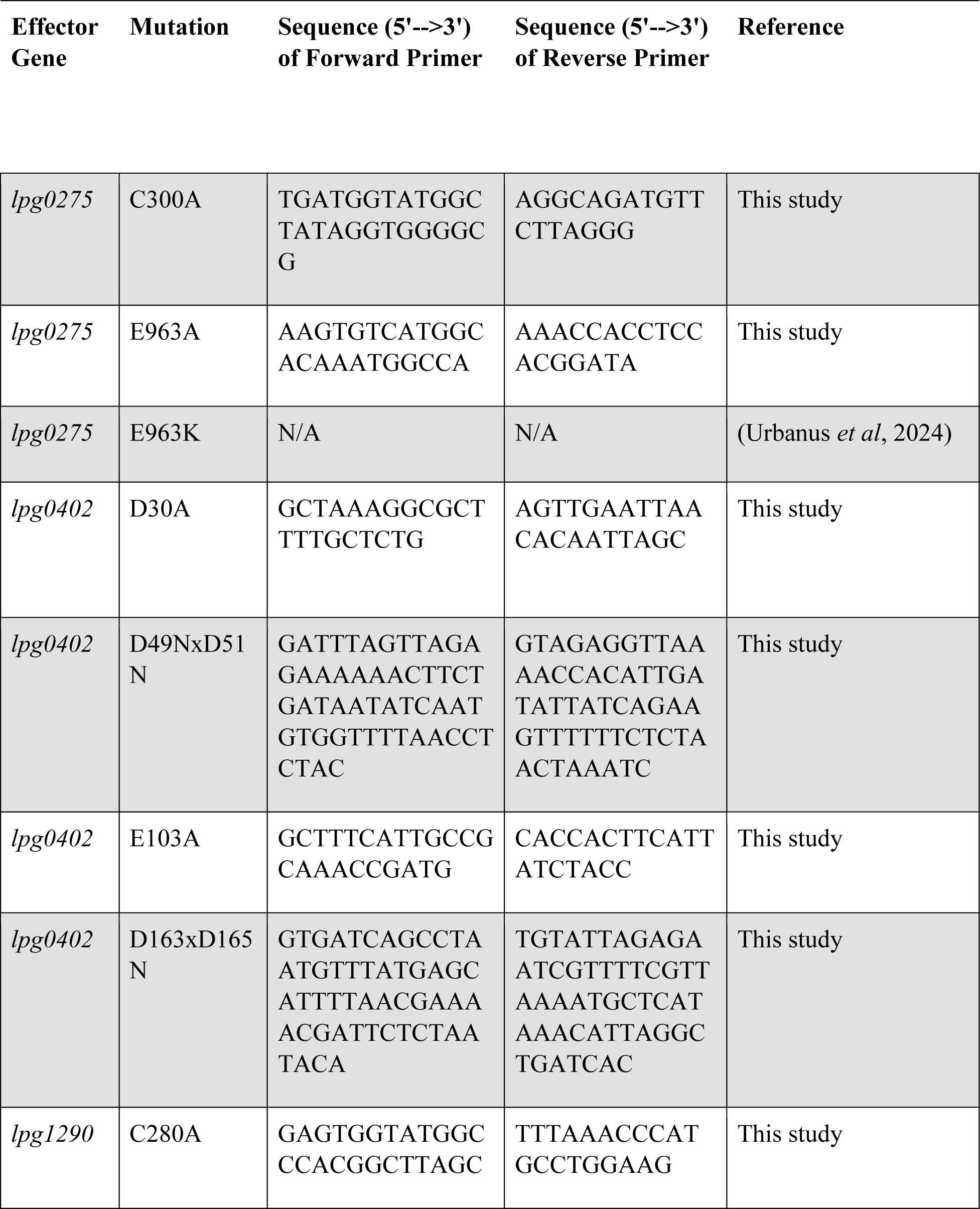
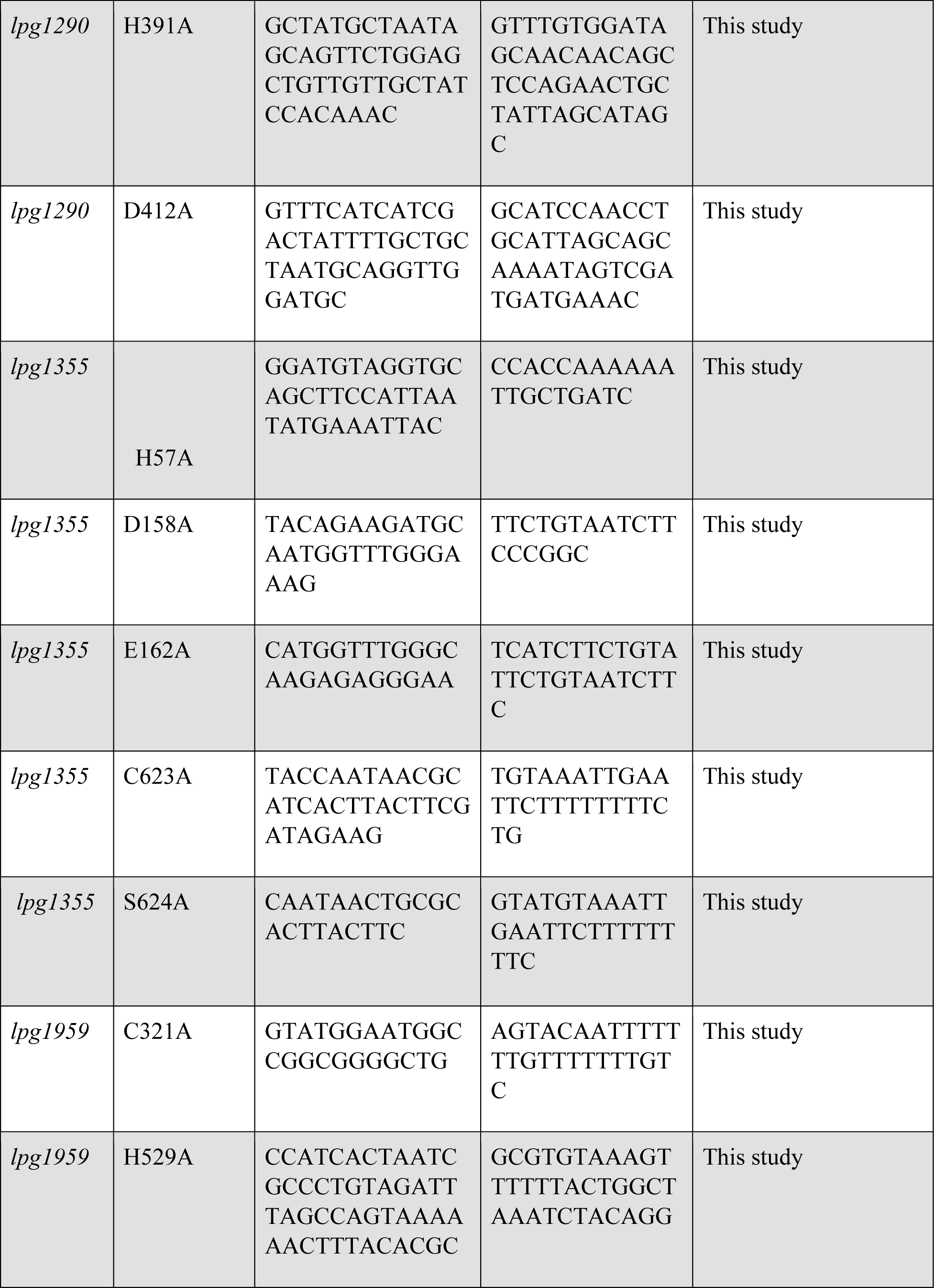
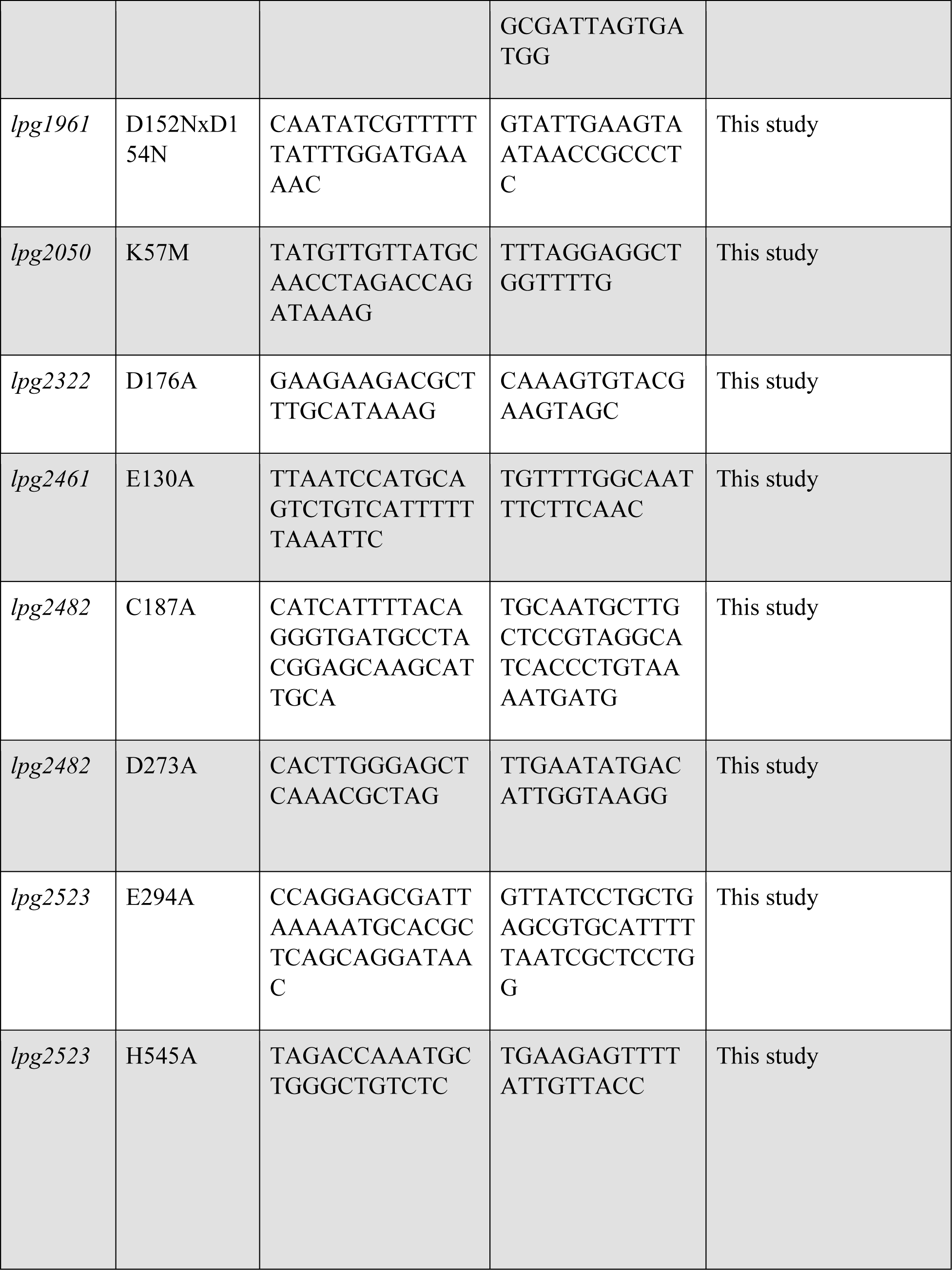
Primers used for the site-directed mutagenesis of yeast overexpression constructs of cryptic enzymatic domains.

**Supplementary Table 6.**
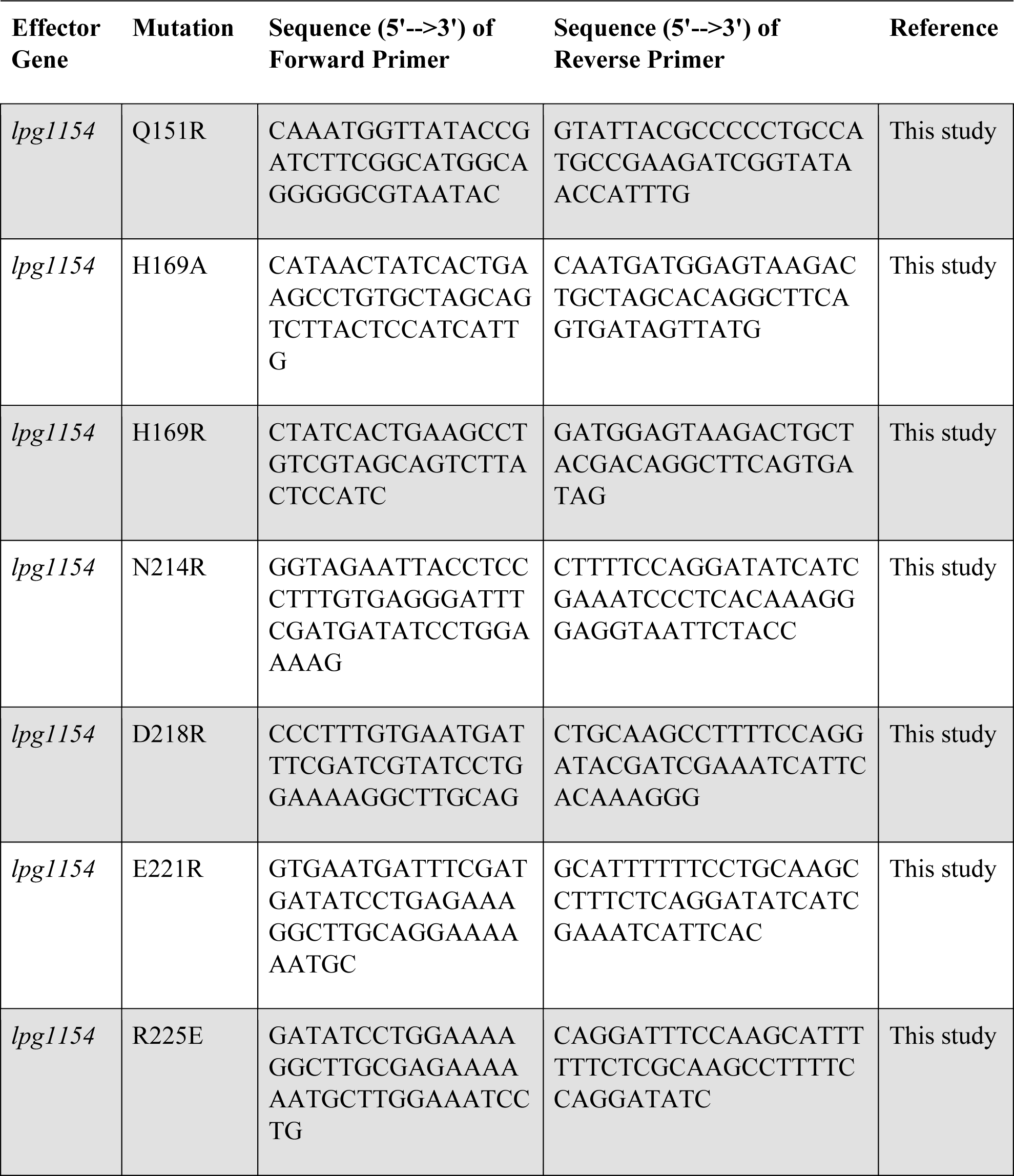
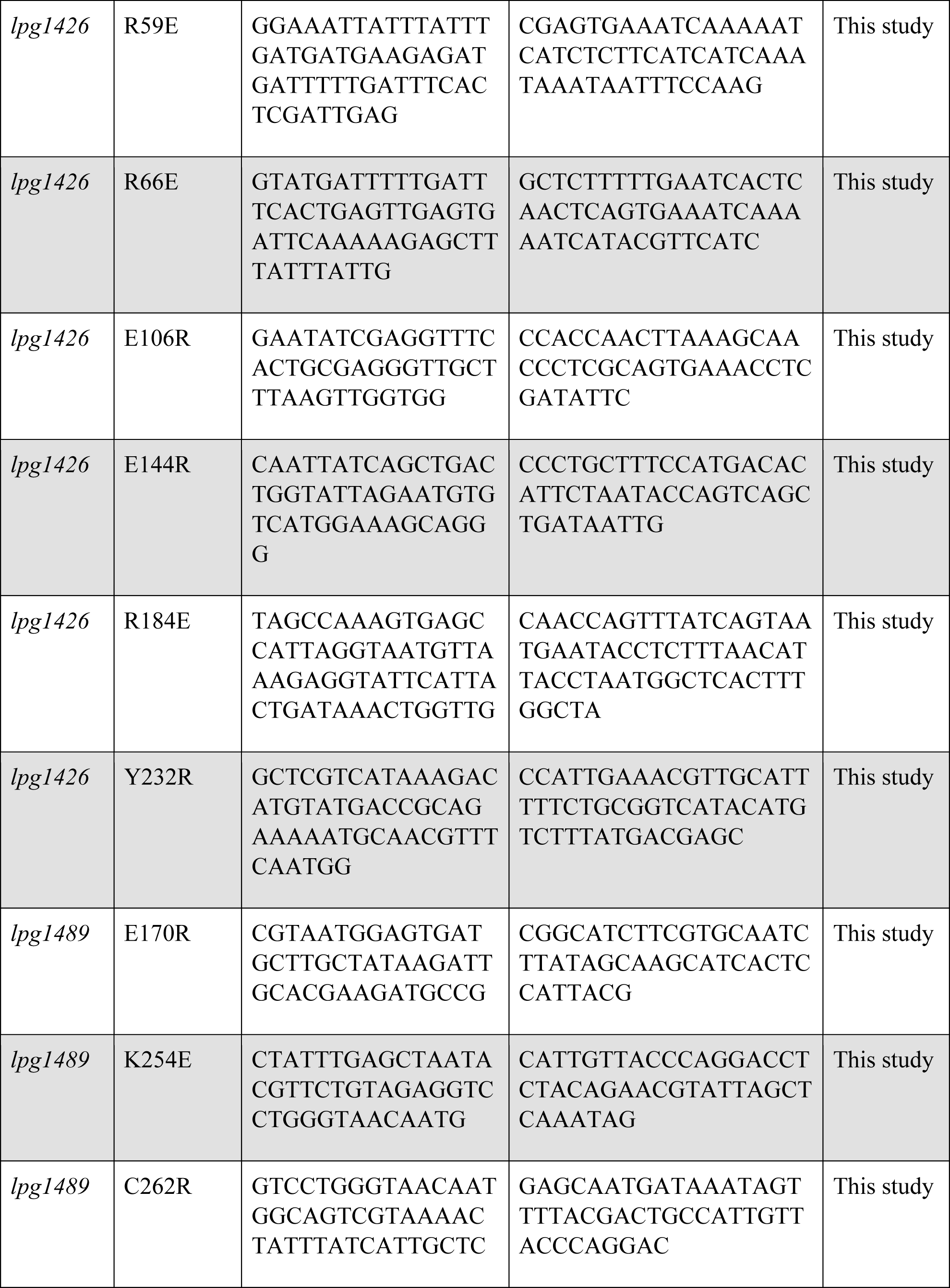
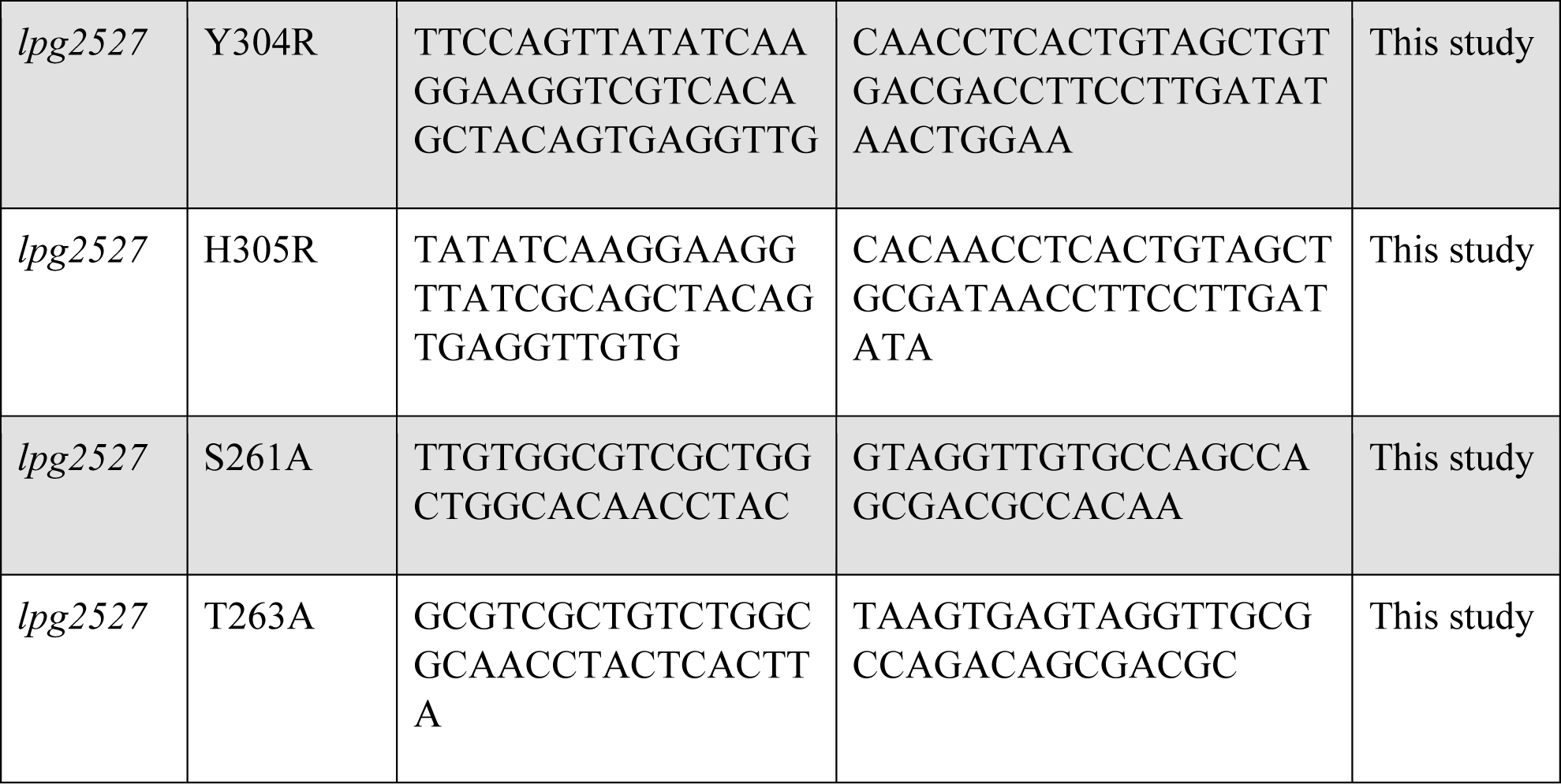
Primers used in this study for the site-directed mutagenesis of plasmids that overexpress effectors containing unique folds/domains in yeast.

